# Organic Convolution in The Ventral Visual Pathway Can Explain the Variety of Shape Tuning in Area V4

**DOI:** 10.1101/2022.05.03.490165

**Authors:** Carl Gold

## Abstract

This modeling study proposes a novel theory for how V4 neurons embody selectivity to the varying degrees of curvature in diverse receptive field arrangements reported in previous recording studies. The model shows that a simple, unsupervised approach to curvature selectivity can explain a wide variety of past observations: V1 aggregates points into selectivity for lines and edges at a variety of rotations; V2 aggregates orientated segments into selectivity for corners at a variety of angles and rotations; V4 aggregates corners into curvature selectivity in a variety of degrees, rotations and positions. The model is implemented in 900,000 integrate and fire units in standard cortical micro-circuits obeying Dale’s law. The unit model and spike timing and transmission use realistic biophysical parameters. A novel method for simulating large numbers of integrate and fire units with tensor programming and GPU hardware is employed: By combining convolution with the time course of post-synaptic potentials, computation occurs in a feedforward cascade of loosely synchronized spikes. The model has relatively few parameters which are tuned by stochastic search with manual fine tuning. A synthesis of hierarchical and convolutional network models, this study adds novel elements to both: In comparison to previous hierarchical models there is a novel V4 mechanism and a biologically realistic computation based on single spikes using tensor convolution as the simulation engine. In comparison to previous convolution models there is a much higher degree of biological realism and a novel unsupervised approach to connection formation.

## 1 Introduction

### 1.1 Organic Convolution

This study demonstrates a novel theory for how V4 pyramidal cells implement rapid recognition of curved and straight image patches by simple patterns of connections with afferent inputs from V2. A model based on the theory can reproduce diverse sets of recordings made in V4 [Pasupathy and Connor, 2001, Nandy et al., 2013]: These studies found that there are V4 neurons that are selective to either high, medium or low curvature. There seem to be some neurons selective to a constant degree of curvature in a certain position and orientation in its receptive field, and other neurons selective to a constant degree of curvature around the entire receptive field with orientation selectivity that rotates with position. Still other V4 neurons are selective to straight or nearly straight contours with an aligned orientation. The inputs to V4 were recently described in [Liu et al., 2016]: V2 contains units with receptive fields that are “complex-shaped” because they combine two Gabor-like receptive fields from V1 input sub-units with each sub-unit at a different orientation. A typical “complex-shaped” V2 unit is selective to a corner where two orientations meet. Curvature selectivity in a V4 unit may be built up from afferent corner selective inputs that have a characteristic range of separation angles and aligned orientations that rotate systematically around the center of the receptive field. The arrangement is akin to the simple arrangement of inputs that leads to motion detection described in [Kim et al., 2014, Borst and Helmstaedter, 2015]: In motion detection afferent inputs, each having a different characteristic delay, connect at different preferred distance along each dendrite of the receiving cell. In this theory for V4 the proposed V4 arrangements of afferent inputs are selective for the form of objects rather than motion, and also have simple yet purposeful arrangements along the dendrites.

If simple connection patterns lead to selectivity in diverse neurons across cortex it relies on the fact that the same connection patterns are repeated by neurons at different locations in topographically organized cortical layers. Convolution is the mathematical model for such a computation: Convolution refers to the mathematical operation common in signal processing in which one function, a filter, is applied to every point in an input signal. (see e.g. [Smith et al., 1997].) This theory for V4 curvature selectivity and the associated model are referred to as Organic Convolution (OC) because it proposes that computation equivalent to convolution can be implemented biologically, and in a realistic biophysical model, not just in V1 but also in V2 and V4.

In OC the primary mechanism for synaptic connection pattern formation is assumed to be a genetically controlled process that guides local connections during healthy development without external stimuli; it is “unsupervised” in the parlance of machine learning. The connection patterns assumed by OC are intended to meet the criteria proposed in [Zador, 2019, Shuvaev et al., 2024] in that they are simple enough to be created through mechanisms controlled by the genome. It remains an open question what forms of learning may complete our understanding of cortical processing biologically plausible learning mechanisms are beyond the scope of this study. As will be detailed in the discussion of background literature, it is assumed that the concerted action of local development rules can enable neurons distributed in a topographically organized layer to have approximately the same patterns of connections. This approach satisfies the primary objection to convolution as a result of a learning process [Bartunov et al., 2018], which is that neurons at diverse locations could not learn the same connection strengths from experience.

All modeling in this study refers to cascades of individual spikes, defined as a computation in which every neuron in every layer fires at most once. Cascades are of interest to biophysicists seeking to explain neural computation for several reasons: As described in [Thorpe and Imbert, 1989, VanRullen et al., 2005], object recognition can be achieved by a single cascade. The fastest possible multi-area cortical computation, a cascade is an elementary unit of computation from which higher level computations may be built. Computation involving rates or bursts and lateral feedback within a brain area or long range feedback between brain areas take place on longer time scales. These more complex modes of computation must build on an initial feedforward cascade that occurs when a stimulus is first presented. While bursts, rates, lateral feedback and long range feedback may be requirements of a fully realistic model of visual recognition, according to [Thorpe and Imbert, 1989, VanRullen et al., 2005] they are not necessary for the fastest form of visual recognition. For these reasons this study focuses on cascades as a useful building block in understanding those other more complex phenomena.^1^

To make a biophysically realistic model based on a convolution calculation, the implementation of OC uses tensor based computation technologies from Deep Learning and Deep Convolutional Neural Networks (DCNN) [LeCun et al., 2015] and adapts it to create a of model individual spikes. The Spike Time Convolution algorithm (STC), adds the time course of post synaptic potentials (PSP’s) onto a standard DCNN model, as described in detail in the Methods Section 4.3. At the same time, the selectivity of the convolution filters in the V1 and V2 models of this study take a standard form for a Hierarchical Model (HM) of cortex [Fukushima, 1980, Riesenhuber and Poggio, 1999]: V1 includes orientation selective units, there are layers of invariance operations, and V2 units have selectivities combining orientations.

The model in this study is biophysically realistic to an intermediate level of neuronal detail: It includes the time course of post-synaptic potentials at the soma of excitatory and inhibitory model neurons obeying Dale’s Law [Dale, 1935] but not details of dendritic compartments or different types of ionic channels. The excitatory and inhibitory units are arranged into standard cortical microcircuits [Douglas and Martin, 2004]. Biophysically realistic time constants are used for the arrival time distribution of spikes in V1, the time course of PSP’s, and transmittal latency between layers and areas. These result in simulations of about 30 ms, the time necessary for a cascade of spikes arriving in V1 to propagate from V1 to V4, as detailed in the Results and Methods (Sections 2.5 and 4.1.1).

There are multiple sources of randomness in the model, detailed in the Methods: The probability of spikes arriving in V1 is proportional to the input intensity, the arrival time of spikes in V1 and the spike transmission time between cortical layers and areas are modeled using Gamma distributions, and the size of PSP’s are drawn from Normal distributions. (Sections 4.1 and 4.1.1) However, the model in this study has far fewer neurons than are present in the ventral visual pathway of mammals, and it does not include a model of transmission failure; for these reasons the model units in this study are proposed as models for averages of redundant faulty units in real cortex. Still, the OC model presented here goes beyond the typical level of “biological plausibility” in a neural network study: This is a model for the biomechanics of computation at a very low level of detail, and may serve as a foundation from which biophysical parameters can be related to high level computational properties.

The network architecture is summarized in Figure 1. As detailed in the Methods (Section 4.1.1), the model starts with V1 L4 units (Figure 1C.) which spike with a probability proportional to the input image intensity at times which are randomly, but tightly synchronized as in real cortex [Singer, 1999]. This input layer serves as a lumped model of the visual pathway from the retina to V1 L4. These spikes are the input to the V1 L2/3 pyramidal units (Figure 1 E,F) and also L2/3 inhibitory units (D), which also send their spikes to the L2/3 pyramidal units. The inhibitory units match the excitatory units one to one and each has enough excitatory input to fire shortly after receiving an input spike, its output matching the input with a delay introduced by its own PSP timing properties - these properties are common to the inhibitory units throughout the model.

**Fig. 1.**
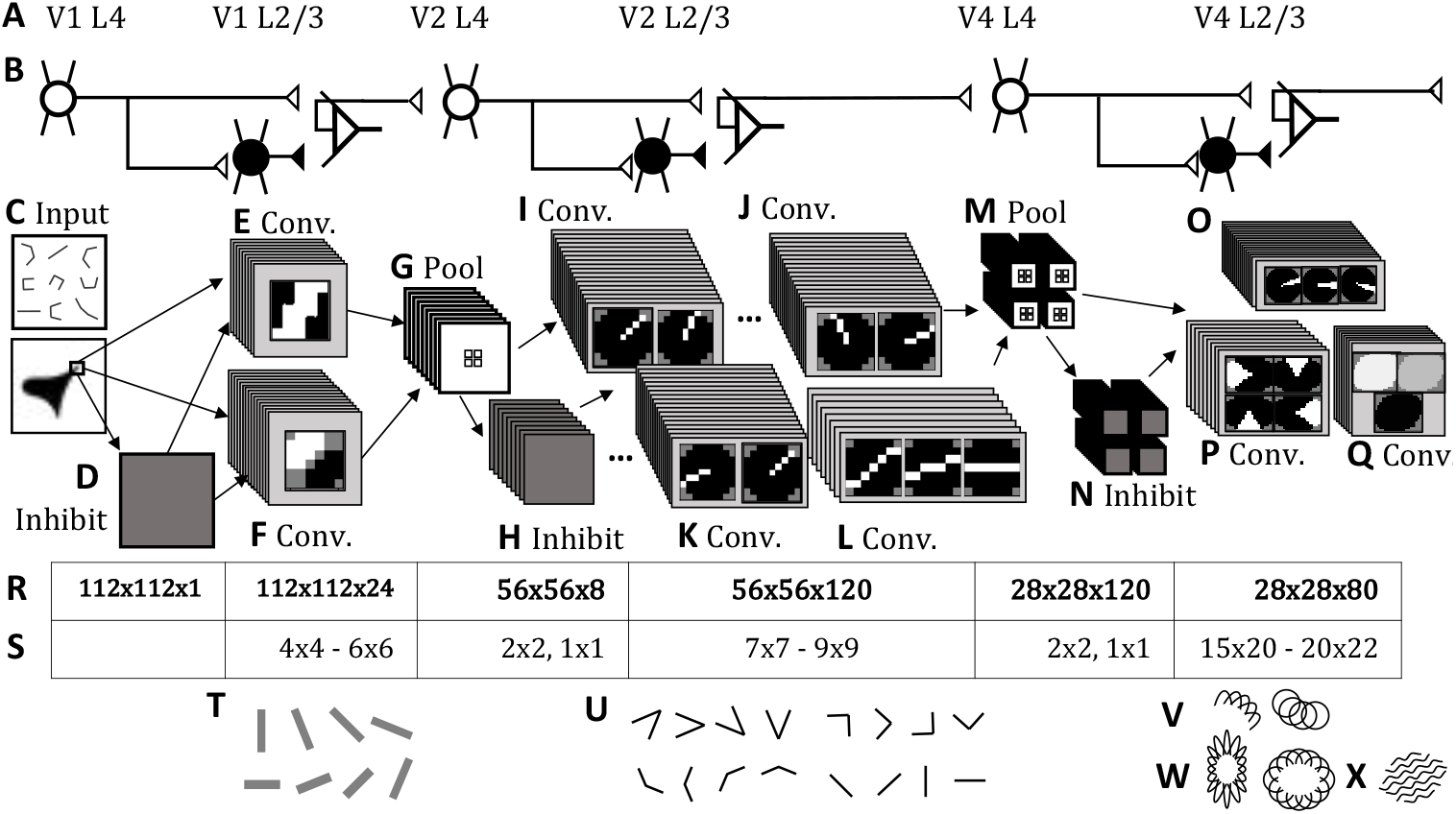
Organic Convolution Model Architecture - see the accompanying text for a detailed explanation. **A** Labeling of network parts by cortical area. **B** Identification of cell types represented by different parts of the network: Stellate cells in L4, Inhibitory and Pyramidal and cells in L2/3. L2/3 pyramidal cells are modeled with a convolution layer (Conv.) while L4 stellate are modeled with a pooling (Pool) layer. Inhibition are modeled as an inversion layer (Inhibit). **C** Input layer, corresponding to V1 L4 stellate cells. D Inhibition of input layer corresponding to V1 L2/3 inhibitory interneurons. **E** V1 L2/3 pyramidal neurons selective to oriented lines represented by eight channels of convolution. **F** V1 L2/3 pyramial neurons selective to oriented edges represented by 16 channels of convolution. **G** The V2 L4 stellate model is 24 channels of pool operations corresponding to the V1 L2/3 channels. **H** Inhibition layer representing V2 L2/3 inhibitory interneurons. **I-L** The V2 L2/3 pyramidal neurons are represented by convolutional channels that combine inputs at a range of angles: acute (I), right (J), obtuse (K), fully straight (L); additional angles are omitted for brevity. **H** The V4 L4 stellate model is 120 channels of pool operations corresponding to the V2 L2/3 channels. **N** Inhibition layer representing V4 L2/3 inhibitory interneurons.**O** V4 L2/3 neurons sensitive to a medium or high degree of curvature over an asymmetric portion of the receptive field represented by 48 channels of convolution. **P** V4 L2/3 neurons sensitive to a medium or high degree of curvature over an a a fully symmetric receptive field represented by 24 channels of convolution. **Q** V4 L2/3 neurons sensitive to low curvature are represented by 8 channels of convolution. **R** The size and number of channels for the layers above. Note that L4 stellate pooling layers and L2/3 inhibition layers share common dimensions and numbers of channels and are presented together. **S** The sizes of the convolution filters for the layers above. **T** Preferred stimuli for the V1 L2/3 units: short, oriented images segments. **U** Preferred stimuli for V2 L2/3 units: small corners at a range of angles, from acute to fully straight lines. **V** Preferred stimuli for asymmetric V4 medium and high curvature selective units. **W** Preferred stimuli for fully symmetric V4 medium and high curvature selective units. **X** Preferred stimuli of the V4 low curvature selective units.

The L2/3 pyramidal units have convolution filters created to resemble the receptive fields of V1 simple cells [Hubel and Wiesel, 1962, Ferster and Miller, 2000]: excitation juxtaposed with inhibition to form either a band (Figure 1E) or an edge (Figure 1F). The band filters have 8 versions at 8 orientations (every 22.5^°^ up to 180^°^) and the edge filters at 16 orientations (every 22.5^°^ up to 360^°^.) Each orientation of edge or band forms a “channel” in the terminology or DCNN: a layer of units having identical input/output properties - in this case, the oriented simple convolution filters. Figure 1T illustrates the preferred stimuli of the V1 L2/3 pyramidal cells. See Section 4.4.2 for details on the V1 model.

The spikes from the V1 L2/3 units are the inputs to the V2 L4 units (Figure 1G). In the model, the L4 units perform a downsampling function known as “pooling” in DCNN or “invariance” in the terminology of HM: These units have 2 × 2 receptive field and sufficient excitation to fire whenever any input fires. Matching orientations of edge selective and pool selective V1 L2/3 units are combined into eight orientated channels in the V2 L4 model (one channel every 22.5^°^.)

The output of the V2 L4 units travels to both the V4 L2/3 inhibitory units (Figure 1H) and the V4 L2/3 pyramidal units. The convolution filters for the V4 L2/3 pyramidal units are based on “complex-shaped” units described [Liu et al., 2016] in that there are units that combine two orientations at a particular angle (Figure 1I-K). In the model there are 6 separation angles, every 22.5° from 45° to 180° (45°, 67.5°, 90°, 112.5°, 135°, 157.5°.). There are also units that have selectivity to long straight line segments, “ultra-long” units in the terminology of [Liu et al., 2016]. The corner selective units have 16 channels per separation angle, with the pattern rotated by 22.5° in each channel. The units selective to extended straight inputs have 8 rotations rotated by 22.5°. Figure 1U illustrates the preferred stimuli of the V2 L2/3 pyramidal cells. See Section 4.4.3 for details on the V2 model. This definition of V2 selectivity is substantially similar to that proposed in [Riesenhuber and Poggio, 1999], albeit in a spiking convolutional form.

The models for V4 L4 (Figure 1 M) and V4 L2/3 inhibitory units (Figure 1 N) are the same as those for V2 L4 and V2 L2/3 inhibition: The V4 L4 units pool and downsample the V2 L2/3 outputs. In the V4 model each separation angle and orientation of V2 L2/3 unit has its own separate channel of units performing pooling and inhibition.

The V4 L2/3 pyramidal cell model uses convolution filters that embody a novel proposal for how V4 units may create curvature selectivity from V2 afferents (Figure 1O-Q): Selectivity to curvature of a certain degree is formed by pooling afferent inputs having a degree of curvature in a particular range: High curvature corresponds to pooling inputs having selectivity to a separating angle less than 90^°^; medium curvature selectivity corresponds to pooling afferent units with selectivity to a separating angle from 90^°^ to 135^°^; low curvature selectivity corresponds to pooling 158^°^ selective units along with straight selective units (Figure 1Q). To achieve curvature selectivity, only similarly oriented V2 corner selective units are pooled in any part of the V4 L2/3 units receptive field. Importantly, the selected orientation of afferent V2 inputs shows a systematic rotation around the receptive field of V4 L2/3 pyramidal neurons - this is more apparent in V4 units with curvature selectivity throughout their receptive field (Figure 1P), and also matches the results for V4 units with curvature selectivity in a particular portion of the receptive field (Figure 1O). Illustrations of the preferred stimuli of the V4 L2/3 pyramidal cells with curvature selectivity in a particular region of the receptive field are shown in Figure 1 V; the preferred stimuli of V4 L2/3 pyramidal cells with curvature selectivity that is symmetric around the receptive field are shown in Figure 1 W; and the preferred stimuli of V4 L2/3 pyramidal cells with selectivity for straight or mostly straight stimuli is shown in Figure 1 X. Details of the models for different types of V4 units are discussed in the Results Sections 2.1 (acute/high curvature selectivity), Section 2.2 (medium curvature selectivity), Section 2.3 (low curvature selectivity) and the Methods Section 4.4.4.

As will be detailed in the Methods Section 4.4, the form of the excitatory and inhibitory patterns in the convolution filters are defined by rules embodying the theory of their function. Those rules have 142 parameters that need to be tuned including 73 parameters that set the strength of excitatory and inhibitory connections and another 68 parameters that set the size of the receptive fields and free parameters of the connection rules (see Table 7.) The majority of the connection weights, for the V1 and V2 convolution models, are set with a novel unsupervised learning algorithm that stochastically optimizes the parameters to minimize the error in response to stimuli sets consisting of preferred and non-preferred stimuli, as described in Section 4.5. The connection strengths of the V4 convolution model were set manually.

The remaining sections of the Introduction provide further details about the prior research that was introduced briefly in this section. The Results section shows the reproduction of the recordings by the model, provides further details about the V4 unit connection patterns that reproduce the recordings, presents data on the propagation and timing of spike cascades in the network, and demonstrates predictions that the model makes about novel stimuli. The Methods section provides details about the spiking neuron model, the Spike Time Convolution computation, the constants of the biophysical model, and the construction and tuning of the convolution filters. The Appendix contains illustrations of the propagation of spikes in every part of the network and details of methods omitted from the main Methods section for brevity.

### 1.2 Visual Area V4

Visual object identification is processed in the ventral visual pathway through primary visual areas V1 and V2, and an intermediate area V4, before object recognition is processed in area IT, or inferior temporal cortex (IT) [Tsao et al., 2006, Kobatake and Tanaka, 1994]. It has been known for many years that there are neurons in area V1 that encode local orientation [Hubel and Wiesel, 1962, Ferster and Miller, 2000].

More recent studies [Liu et al., 2016] have found that V2 contains neurons in which V1 receptive fields converge into two main types: The first type of V2 selectivity is described as “ultralong” (aligned) which combines afferent inputs from orientation selective V1 neurons that all share the same orientation, effectively extending the short oriented selective patches into selectivity for longer straight contours in V2. The second type of selectivity seen in V2 neurons is “complex-shaped” which combines multiple oriented sub-unit at an angle, forming a corner where two orientations meet.

The functional properties of V4 are less well understood: studies suggest that area V4 occupies an intermediate level in the hierarchy of visual processing, responding to moderately complex stimuli, but there is not consensus on the precise nature of V4’s processing [Roe et al., 2012]. In this last two decades, multiple streams of evidence suggest that V4 neurons are selective to different degrees of curvature in a variety of configurations. [Pasupathy and Connor, 2001, Nandy et al., 2013, Hu et al., 2020]

Two examples of stimuli used in studies recording from individual V4 neurons are shown in Figure 2. The stimuli from [Pasupathy and Connor, 2001] are shown in Figure 2 A: The stimuli consist of systematic combinations of convex and concave boundary elements, yielding 366 total stimuli. [Pasupathy and Connor, 2001] found that individual cells showed consistency to preferring one type of boundary element, for example an acute convex curvature, in a specific position. (See Figures 3 and 7) Later studies using similar methods found that some V4 neuron seem to prefer texture [El-Shamayleh and Pasupathy, 2016]. More recently, it was found that a given convex or concave shape preference tends to be selective for similar stimuli with a range of sizes [Kim et al., 2019].

**Fig. 2.**
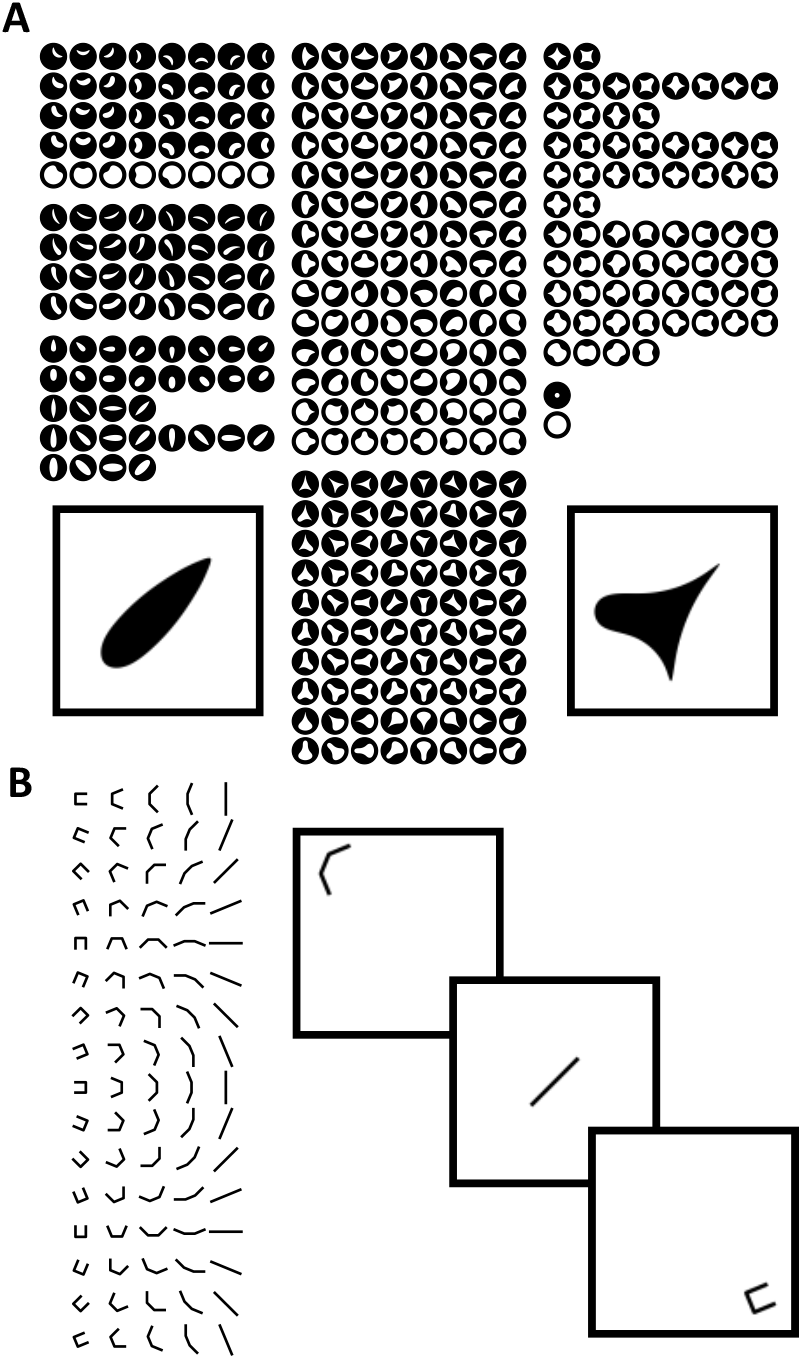
Stimuli used to probe the response of V4 neurons. **A** [Pasupathy and Connor, 2001] used systemtatic configurations of convex and concave boundary elements included sharp convex angles, medium and broad convex curves, and medium and broad concave curves. Each stimulus is represented by a white icon positioned within a black disk that represents the cell’s RF. The stimuli are arranged into blocks according to the number and configuration of convex projections. **B** [Nandy et al., 2013] used composites of the 3 bars at five different conjunction angles (0^°^, 22.5^°^, 45^°^, 67.5^°^ and 90^°^). The five conjunction angles are presented at 16 orientations for a total of 72 unique stimuli. The composite shapes are presented on a grid that spans the receptive field. Insets: Examples of stimuli as presented.

**Fig. 3.**
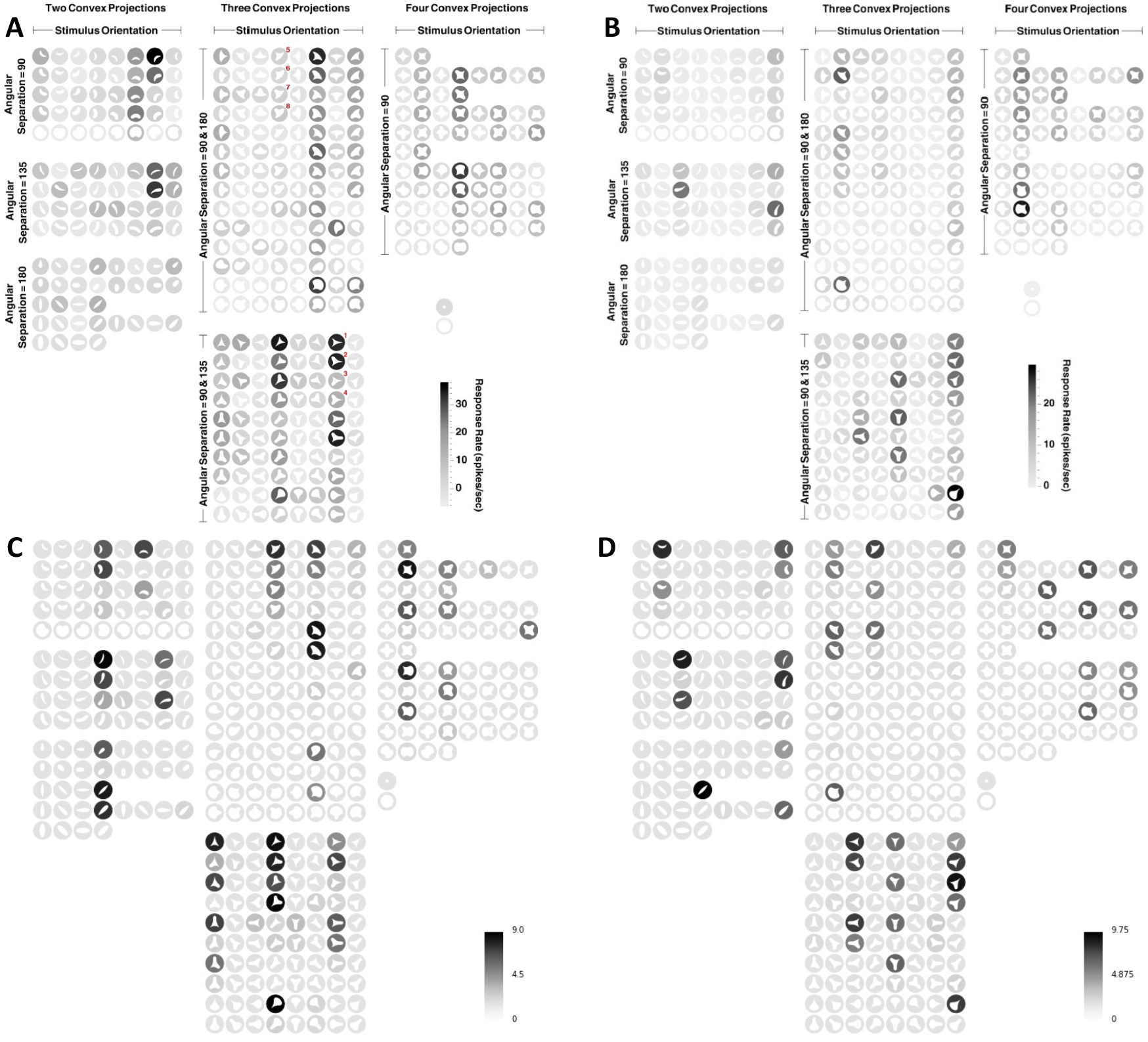
Recording and simulation of Acute convex asymmetric selectivity in V4: The white shape in each location represents the the stimuli shape, which is actually presented as a black silhouette on a white background. The shading of the disc around the shape represents the intensity of the response as indicated by the legend. **A** Recording reproduced from [Pasupathy and Connor, 2001] showing a neuron tuned for acute convex curvature at the bottom left. **B** Recording reproduced from [Pasupathy and Connor, 2001] showing a neuron tuned for acute convex curvature at the top right. **C** Result of a model simulation qualitatively matching **A**. The results shown are the number of spikes out of 10 simulations. **D** Result of a model qualitatively matching **B**.

The stimuli from [Nandy et al., 2013] are shown in Figure 2B: the stimuli consisted of oriented bars combined to form approximate curves ranging from straight to tightly curved at a range of orientations. These elements were sized to be smaller than the receptive field of a single V4 neuron and presented at multiple locations throughout the receptive field. This approach allowed the detailed investigation of shape tuning throughout the receptive field. [Nandy et al., 2013] found that the V4 neurons responsive to these stimuli can be categorized into those selective for low curvature, medium curvature and high curvature. Only low curvature selective V4 neurons were found to have spatially invariant shape tuning: V4 neurons selective to medium and high curvatures had spatially variable tuning. [Nandy et al., 2013] showed units that were selective to a consistent degree of curvature but at varying orientations throughout the receptive field. (See Figures 5, 9 and 11.)

Recently [Srinath et al., 2021] measured V4 neuron responses with a dataset generated on the fly by a genetic algorithm designed to find shapes that had the highest neuronal response. This dataset was more complex than those of [Pasupathy and Connor, 2001] and [Nandy et al., 2013] in that it included not only flat imagines (against dark and light backgrounds), but also three dimensional shapes. However, the stimuli of [Srinath et al., 2021] did not include lines or outlines of shapes. [Srinath et al., 2021] found that many V4 neurons seem to be selective to three dimensional shapes more than a flat projection of a similar shape. This suggests that the form of complex objects is already represented to some degree in V4 and not only in IT. However, [Srinath et al., 2021] did not characterize the shape selectivity of V4 neurons by categories like the acuteness or orientation of the selected shapes. Applying the model of this study to the data of [Srinath et al., 2021] is an attractive area for future research.

Another recent study supports the notion of orientation and curvature tuning in V4: [Hu et al., 2020] used intrinsic optical imaging of V4 and found a pattern of curvature domains that were tuned to respond to either straight, medium curvature or high curvature. In [Hu et al., 2020] the stimuli consisted of conventional straight line gratings along with gratings having different degrees of curvature, all presented at different orientations. The results in [Hu et al., 2020] are complimentary to the results in [Nandy et al., 2013] and [Pasupathy and Connor, 2001], suggesting that neurons sensitive to different degrees of curvature are in fact central to the functioning of V4.

### 1.3 Hierarchical Models of Visual Cortex

Hierarchical models (HM) of visual cortex have a long history of explaining how cells that respond to complex visual stimuli may be built from cells that respond to simple stimuli. The most well known early example was the Neocognitron [Fukushima, 1980] which used an alternation between feature extracting cells, modeled after V1 simple cells, and cells modeled after complex units that pool together the input from multiple feature recognizing cells. The feature selective units in the first layer of this and later models correspond in some form to orientation selective units first identified in [Hubel and Wiesel, 1962]. The idea was further developed in [Perrett and Oram, 1993] which extended the idea to achieve invariance over a wider range of visual features. [Wallis and Rolls, 1997] used a four level network named Visnet and showed that a Hebbian inspired learning rule could train units to have invariance to the position and size of faces and other simple stimuli.

Another example of a mathematical model using alternating selectivity and invariance operations was the HMAX model [Riesenhuber and Poggio, 1999]. The HMAX model had two layers of simple (feature recognizing) and complex (feature pooling) cells, followed by a learned layer of view tuned cells. The view tuned cells were trained to have invariance to moderately complex shapes. The HMAX model was later extended specifically to match the V4 shape tuning data of [Pasupathy and Connor, 2001] in [Cadieu et al., 2007]: this version of the model includes orientation selective units at a variety of orientations and scales to represent V1, followed by a pooling layer creating invariance. Another layer of features selective to combinations of orientations represents V2, and a second layer of invariance generating combinations is identified with V4. In [Cadieu et al., 2007], the V4 combination layer was fit using a selection algorithm to match the data from [Pasupathy and Connor, 2001].

[Hubel and Wiesel, 1965] suggested that end stopped simple cells would have a degree of curvature selectivity, and the idea was developed into a mathematical model by [Dobbins et al., 1987]. The idea that end stopping was related to curvature selectivity received little further attention until recently [Ponce et al., 2017] showed that in V1, V4 and IT end stopping is correlated with curvature selectivity in microelectrode array recordings in behaving monkeys. This study takes a minimalist approach to the model for V1 and does not assume end stopping, and whether end stopping may enhance curvature selectivity is an attractive area for future research.

In comparison to [Cadieu et al., 2007], the most similar prior work, the Organic Convolution model replaces learning in the creation of V4 selectivity with an unsupervised mechanism that assumes a large number of generic V4 selectivities for different curvatures and rotations are present. Further, this study proposes that an important unsupervised connection pattern in creating curvature selectivity in V4 is that the V2 afferents orientations preferred by the V4 neuron rotates with the topographical position around the center of the receptive field.

The HM’s described here use a variety of mathematical descriptions for the activity of the neuronal representations. The models are generally interpreted as representing neuronal firing rates in response to a constant stimuli. From [Fukushima, 1980] to [Cadieu et al., 2007], the typical approach is some form of normalized dot product to represent the activity of a feature selective units, and then a non-linearity such as a hinge or a sigmoid function to determine the resulting activity level (firing rate) of the unit.

This model embodies a novel biologically realistic cascade spike computation and a simulation implemented using tensor computational tools. None of the HM’s surveyed here used a representation of individual spikes, although it has been shown that object recognition in humans and animals is too fast to rely on a rate code [Thorpe and Imbert, 1989]: rapid recognition happens so fast that there is only time for a single spike per layer in each brain area. This means a single cascade is sufficient to perform recognition. The issue of recognition speed and implications for spiking computation was also described in [Wallis and Rolls, 1997].

### 1.4 Deep Neural Networks

In the last decade deep convolution neural networks (DCNN’s) [LeCun et al., 2015] were developed by artificial intelligence researchers that dramatically improved performance in visual recognition tasks. In fact, the idea of convolution in a neural network is inspired by the same observations about cortex as HM’s : First, there is the observation that selectivity to simple oriented stimuli in earlier layers seem to repeat more or less interchangeably over positions in topographically organized visual space. That makes it well represented by convolution, and even HM’s have used convolution to implement their selectivity mechanisms [Fukushima, 1980]. Also the alternation of filter operations with non-linear invariance operations is taken from the idea of complex cells which became “pooling” layers in DCNN [Carandini et al., 2005]. However, modern DCNN use stochastic gradient descent training algorithms broadly known as deep learning (DL) to determine the selective unit properties rather than design them to match neurophysiological data.

Modern DCNN’s have converged on a hinge activation function, known as RelU which is shortfor Rectified Linear Unit. The RelU activation function has desirable computational properties and is taken to be a model of a neuron where the firing *rate* increases as a function of the total input [Glorot et al., 2011]. However, RelU activation does not provide a model for computation with individual spikes.

There has been considerable interest among computational neuroscientists in whether DCNN and DL can be the basis for explaining the object recognition pathway [Yamins et al., 2014, Kietzmann et al., 2019]. However, there are questions as to whether DL models can serve as useful models for neuroscience: DL requires backpropagation of error signals [Rumelhart et al., 1986] carrying gradient information with the same weight as the feedforward signal, but such signals have not been observed [Bengio et al., 2015]; as a result DL in the form originally proposed is biologically implausible. It has also been noted, even by DL researchers, that no learning system can guarantee that the same weights will be learned by neurons at different locations [Bartunov et al., 2018] raising questions about the plausibility of the convolution model. Like hierarchical cortical models, DL models do not model spikes but instead rely on non-linear transfer functions, which are described as representing firing rates [Glorot et al., 2011]. DL models also ignore the fact that real neurons are constrained by their type to have entirely excitatory or entirely inhibitory influence on efferent connections [Cornford et al., 2020]. At the same time, there have recently been some serious attempts to investigate whether biological learning mechanisms like spike timing dependent plasticity (STDP) in a rate coded model can implement biological deep learning [Bengio et al., 2017, Kheradpisheh et al., 2018].

A few studies have attempted to use DL techniques to understand neurophysiological data from V4: [Pospisil et al., 2018] examined a state of the art DL model and found units that respond to the shapes used in [Pasupathy and Connor, 2001]. [Yamins et al., 2014] proposed a goal driven approach and trained a DL model to match the response of IT neurons; the network trained to match IT also matched V4 response properties. These studies are described by their authors as “top-down” in the sense that they create models to perform the functions of higher cortical areas and then try inspect the resulting learned connections to gain insight into lower cortical areas. [Cadena et al., 2019] also took the goal driven approach by analyzing a pre-trained DCNN trained with DL and added output non-linearities and noise to the unit activations to come up with spike counts. They found that intermediate convolution layers in the network best match V1, but did not consider visual areas beyond V1.

Despite high performance on AI vision benchmarks, there are several ways in which DL models are a poor match for visual processing in the brain: DL networks seem to rerquire much more data to learn concepts than biological brains, DL networks suffers from so-called “adversarial” attacks (that degrade performance with almost imperceptible changes in an image) and failure to generalize that are inconsistent with human behavior[Serre, 2019]. [Jacob et al., 2021] found that DCNN trained with DL can reproduce some qualitative properties of natural vision (e.g. Thatcher effect), but others were absent (e.g. surface invariance). [Feather et al., 2023] looked at what images are seen as invariant to humans and DL networks (termed metamers) and found distinctly different patterns of invariance in DL and biological networks.

### 1.5 Learning vs. Unsupervised Connection Formation

A more meta problem with not only DL but also many HM’s is the assumption that the neural network is initially in a random state and useful behavior is learned from experience. While the requirement to “learn” connections through an active process has long been assumed in neural modeling, [Zador, 2019] argues that the competitive demands on newborns require brains to be “initialized” to perform significant functions. While true in mammals, this is even more the case in non-mammalian species with complex behavioral repertoires. [Zador, 2019] concludes that there must be an efficient encoding for the generation of such structure in the genome because the genome is not large enough to specify individual connections in complex brains, an idea that is developed in a mathematical model in [Shuvaev et al., 2024].

Studies have already demonstrated this kind of structure : For example, pyramidal cells in mouse V1 display orientation selectivity at the time of *first eye opening* [Ko et al., 2013]. If orientation selectivity is present at that time its development must have been largely innate, or “unsupervised” in the parlance of machine learning. Another example is that both mammalian [Kim et al., 2014] and fly [Borst and Helmstaedter, 2015] visual systems detect motion by having specific sub-types of bipolar cells, each having a different characteristic delay, connect at different preferred distance along each dendrite of the receiving amacrine cell. This is an example of a “pre-programmed” computation defined by simple arrangements of connections from different types of inputs at different distances from the soma. A variety of pathways and signals control the shape of axonal structures and even the specific connections between neurons of different types [Mueller, 1999] as well as the specific subtypes making up topographically mapped neuronal structures [Shirasaki and Pfaff, 2002].

While [Shuvaev et al., 2024] provides a mathematical framework for how complex learned weights may be compressed through a “genomic bottleneck” the approach here is more heuristic, leaning heavily on the principle known as Occam’s razor: At every stage, the patterns are simple and clearly among the least complex forms that can reproduce the theorized biological function. The resulting model has several orders of magnitude fewer free parameters than a comparable DCNN (for details see Section 4.4). It is concluded to be self-evident that genes *could* control the generation of such simple structures, although proof and elucidation of the mechanism is beyond the scope of this study.

## 2 Results

### 2.1 Acute Convex Selectivity

#### 2.1.1 Composite Boundary Stimuli

[Pasupathy and Connor, 2001] showed that some V4 neurons are selective to acute convex curvature at different positions relative to the center of the receptive field; Figure 3A and 3B are reproduced from [Pasupathy and Connor, 2001]. The stimuli were previous shown in Figure 2A and consist of 366 combinations of convex and concave boundary elements. While the actual stimuli are presented as a silhouette against a light background, in Figure 3 the shape is shown in white and the shading represents the intensity of the neuron response as detailed in the legend. [Pasupathy and Connor, 2001] described the neuron in Figure 3A as “V4 neuron tuned for acute convex curvature at the bottom left” based on the consistent element among all of the stimuli to which the unit has a high response. Similarly, the neuron in Figure 3B is described as a “V4 neuron tuned for acute convex curvature at the top right” on the basis of the fairly obvious common elements among all of those stimuli to which the neuron responds.

Figure 3C and 3D are results for simulated V4 neurons with similar response properties to Figure 3A and 3B respectively. The simulated neuron in Figure 3C is qualitatively similar to Figure 3A in that both consistently respond to shapes with an acute convex curvature located in the bottom left. Although not a perfect match, the model matches the primary feature of the recording; the main weakness of the model is in being a bit more consistent in firing to some acute convex curvatures that the real neuron ignores. The same qualities also apply to the simulated recreation of the recording in Figure 3B with the simulation in 3D which is the neuron tuned for acute convex curvature in the top right. The model was manually tuned based on the published image without the data used to produce Figure 3A and 3B, as described in the Methods.

The connection pattern resulting in the selectivity of Figure 3B are shown in Figure 4. The figure shows convolution filters which implement the synaptic weights to the topographically arranged inputs, with the center of the unit receptive field represented at the center of the filters. The convolution filters are the model embodiment of the connection patterns that give rise to selectivity for acute convexity in a particular position. For further explanation of how connection patterns to topographically organized afferents manifest and how they are represented by the con-volution filters in Figure 4, see the Methods Figure 19 and the accompanying discussion.

**Fig. 4.**
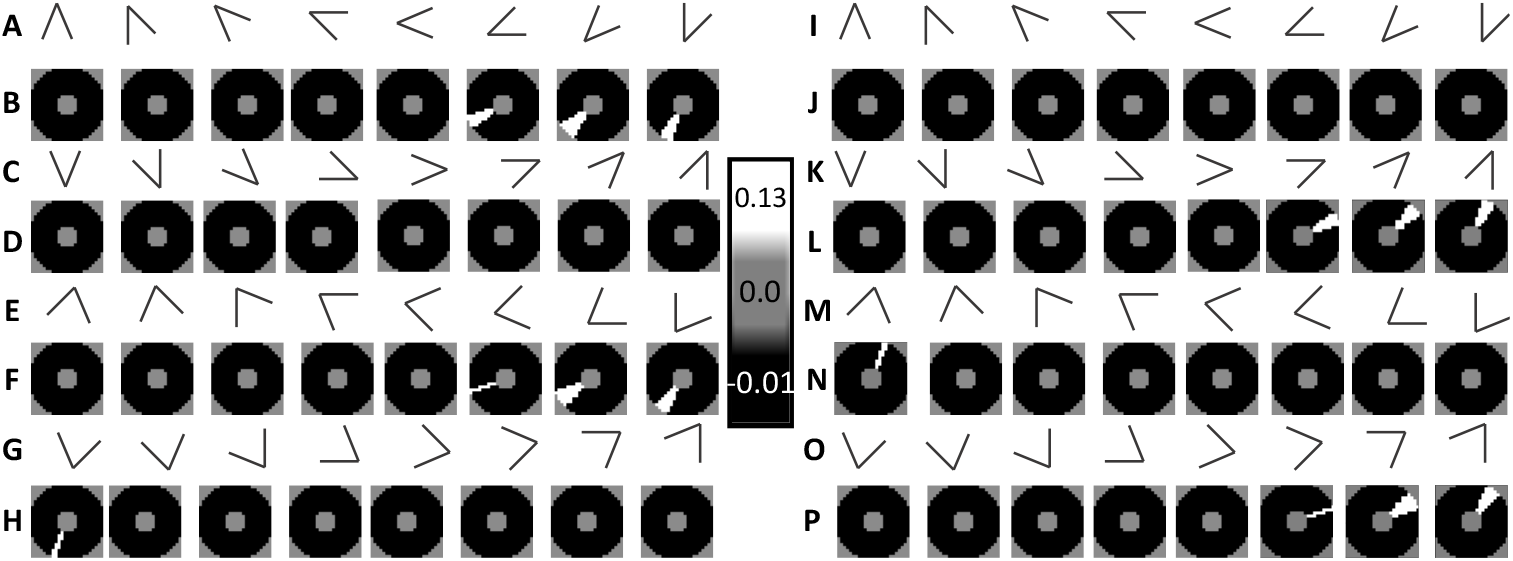
Afferent Connections for model V4 L2/3 neurons selective to high curvatures in asymmetric zones. **A - H:** Connections for the model V4 neuron selective to acute convex curvature at the lower left. **I - P:** Connections for the model V4 neuron selective to acute convex curvature at the top right. **A & C:** Pictorial representation of the preferred stimuli of V2 afferents inputs to V4 having a 45° interior angle at a complete range of rotations every 22.5°. Details of V2 neuron selectivity are described in the Methods 4.4.3. **E & G:** Pictorial representation of the preferred stimuli of V2 afferents inputs to V4 having a 67.5° interior angle at a complete range of rotations every 22.5°. **B, D, F, H:** The synaptic connection strength versus position in topographic space relative to the soma at the center for a V4 neuron selective to acute convex curvature at the bottom left. Each square is a single 9×9 convolution filter implementing the weights for the V2 afferent inputs indicated by the pictogram above it. White=excitatory, Black=Inhibitory, Gray=No connection. All of the weights in B-G together are the connections to the topographically arranged inputs for one V4 neuron. The selectivity is due to excitatory inputs on afferents from corner selective V2 units that are convex and in the bottom left portion of the receptive field. There are also excitatory inputs from slightly rotated orientations of the V2 unit, at positions rotated around the receptive field so that the preferred stimuli remain convex. Other connections to afferent inputs are weakly inhibitory. **I, K, M, O:** Pictorial representation of the preferred stimuli of V2 afferents inputs to V4 as in A/C/E/G. **J, L, N, P**: The synaptic connection strength versus position in topographic space relative to the soma at the center for a V4 neuron selective to acute convex curvature at the top right as in B/D/F/H. There are excitatory inputs on afferents from corner selective V2 units that are convex and in the top right portion of the receptive field. There are also excitatory inputs from slightly rotated orientations of the V2 unit, at positions rotated around the receptive field so that the preferred stimuli remain convex. Other weights on afferent inputs are weakly inhibitory.

**Fig. 5.**
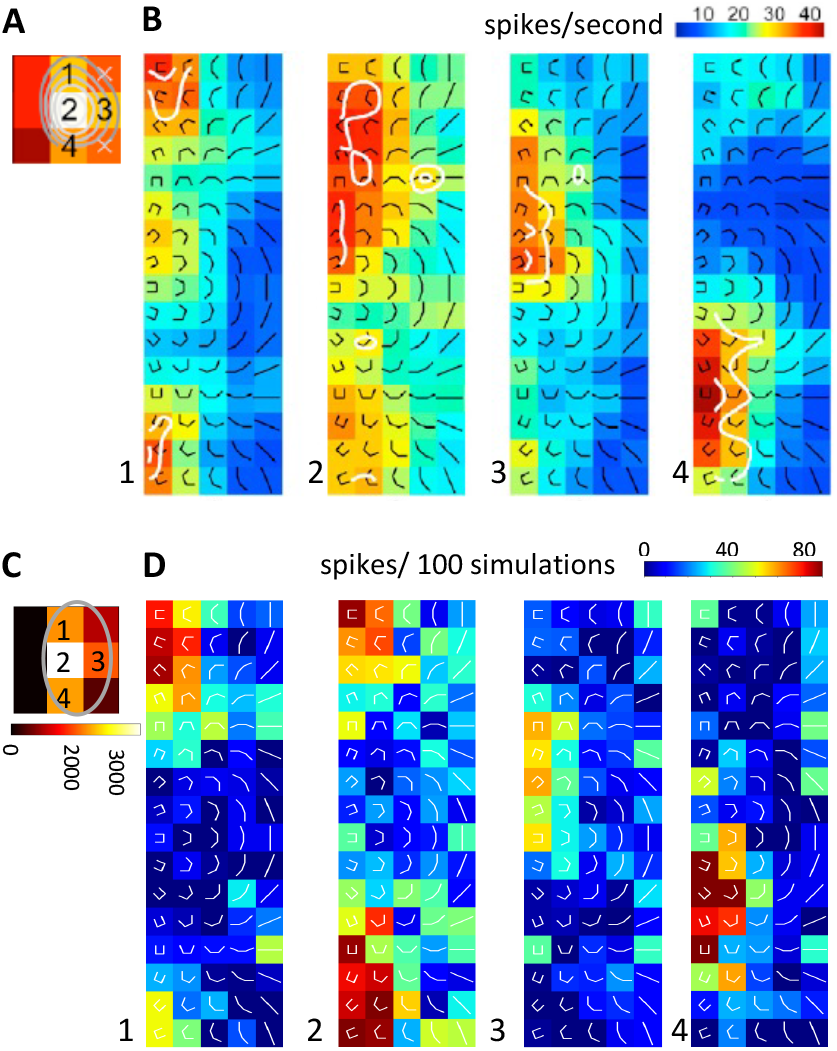
High curvature selective unit recording (top) and simulation (bottom) **A** Intensity of response at 9 positions in the stimuli grid. The numbers index the four positions with the highest response. The shading indicates the overall response intensity. **B** Shape tuning for recording, 1-4 show the shape tuning at the indexed positions in the receptive field. Each pictogram represents a composite bar stimuli that varies by curvature (x-axis) and orientation (y-axis). The color coded scale shows the response firing rate. **C** Receptive field for simulation. The approximate location of the simulated neurons dendritic tree in the topographic visual field is shown. **D** Shape tuning for simulation, as in B, at positions in the simulated receptive field (C).

The afferent inputs relevant to the V4 L2/3 neurons selective to acute convex curvature are represented by pictograms Figure 4A,C, E and G: These are the outputs of V2 neurons selective to corners having interior angles 45 ° (A,C) and 67.5 ° (E,G). The corresponding convolution filter weights are shown below each pictogram, Figure 4 B,D, F and H. The actual inputs to V4 L2/3 have passed through the V4 L4 neurons which reduced the dimension of the representation by two, and the V4 inhibitory neurons which provide the inhibitory version of the same selectivity slightly delayed. So areas shown as excitatory in Figure 4 are actually connections to the V4 L4 neurons, and those that are inhibitory are connections to the V4 inhibitory neurons. As detailed in the methods, these are implemented as two separate channels and two separate convolution filters but for simplicity and conciseness the strength of input connections are shown together and described by the selectivity of the V2 L2/3 neuron from which the afferents inputs originate.

The pattern of excitatory connections is simple: convex angles selective inputs on the bottom left are strongly excitatory and to an equal degree; all other angles are inhibited with a weaker and more diffuse strength. The primary connection is to convex 45^°^ and 67.5^°^ interior angle corners pointing at a 225^°^ angle clockwise from horizontal. At the same time, there are connections to corners angled 202.5^°^ and 247.5^°^ - these connections are in strips that rotate around the center of the receptive field so that the afferent inputs always have a selectivity that is convex. The center of the receptive field lacks excitatory connections - excitatory connections near the center of the V4 L2/3 pyramidal model neurons lead to poor matches with almost all the V4 recordings and this rule was followed to varying degrees in all of the models. This pattern of afferent connection is meant to represent a simplified or average pattern that might be seen in many neurons, each of which would have subtly different selectivity resulting from its own particular dendritic structure and modifications of synaptic weights acquired from experience. The connection pattern for acute convex curvature at the top right is shown in Figure 3D and is qualitatively similar to the neuron tuned for acute convex curvature at the bottom left. The overall preferred stimuli set for the pattern were illustrated in Figure 1V: it is a set of high curvature overlapping curves in a particular region of the receptive field.

#### 2.1.2 Grid of Composite Bar Stimuli

[Nandy et al., 2013] also found that some neurons are selective for high degrees of curvature, and also that some units were selective to high degree of curvature throughout the receptive field with an orientation preference varied with position. Figure 5A and B are reproduced from [Nandy et al., 2013] and represent the output of a V4 L2/3 neuron that is selective to a high degree of curvature throughout its receptive field. Figure 5A shows the intensity of the response at each location in the stimuli grid and the numbered positions identify the highest response; the numbers index the position from top left to bottom right. The color in 5A represents the overall intensity of the response. Figure 5B details the response at each of the four indexed positions from Figure 5A: Each set of icons represents one position, and each icon represents a single stimuli. The color shows the intensity of the response as indicated by the legend.

For this neuron, the stimuli locations triggering the highest response lie along a vertical strip of three locations (Fig 5B1, 5B2 and 5B4), with one additional high response receptive field location on the right side of the main strip (Fig 5B3). The neuron responds primarily to highly curved stimuli, with broad tuning that includes moderate responses to low curvature and straight lines, especially for the most responsive stimuli locations (Figure 5B2). The orientation of the preferred stimuli varies by stimuli location. The response tuning curve at each stimuli location is broad but it peaks around curves oriented convexly towards the center of the receptive field: Location 1 (top) is more strongly tuned to curves open downward, location 2 (center) is tuned strongly for both upward and downward facing curves, location 3 (on the right) is responsive to left opening curves and location 4 (on the bottom) is responsive to upward facing curves. But the convex orientation of the most preferred stimuli does not appear consistent because of the broad tuning of the response.

The output of a V4 L2/3 model neuron with similar high curvature selective response is shown in Figure 5C and Figure 5D. The model response is qualitatively similar to that of the recording in the preference for high curvatures and it approximately matches the orientation selectivity of the recording as well. The main mismatch between the model and recording is that the real neuron shows broader tuning to orientations with high curvature. The model was manually tuned based on the published image without the data used to produce Figure 5A and 5B, as described in the Methods.

The connection pattern to afferent inputs for for the [Nandy et al., 2013] high curvature selective V4 L2/3 cell model is shown in Figure 6. The afferent inputs to the V4 L2/3 neuron are represented by pictograms Figure 6 A,C, E, G, I and K: These are the outputs of V2 neurons selective to corners having interior angles 67.5 ° (A,C), 90 ° (E,G), and 112.5 ° (I,K). The corresponding convolution filter weights are shown below each pictogram, Figure 6 B,D, F, H, J and L. For a detailed explanation, see the text accompanying Figure 4 in Section 2.1.1. The pattern is that for every type of afferent input, half of the receptive field is excitatory and half is inhibitory, and a small region at the center is neutral. Afferent inputs that are approximately convex to the center of the receptive field are excitatory, and afferent inputs that are approximately concave to the center of the receptive field are inhibited. To achieve this, the excitatory/inhibitory boundary rotates around the receptive field, depending on the orientation of the afferent inputs. The boundary is not exactly even - to better match the recording, there is a rotational offset of 45^°^ from perfectly convex orientation of the excitatory inputs. For details, see the Methods section 4.4.4 and Table 11. Non-preferred orientations are inhibited and afferent inputs with higher interior angle complex shape preferences are ignored. The preferred stimuli for the pattern were illustrated in Figure 1W: it is a set of high curvature overlapping curves all around the receptive field.

**Fig. 6.**
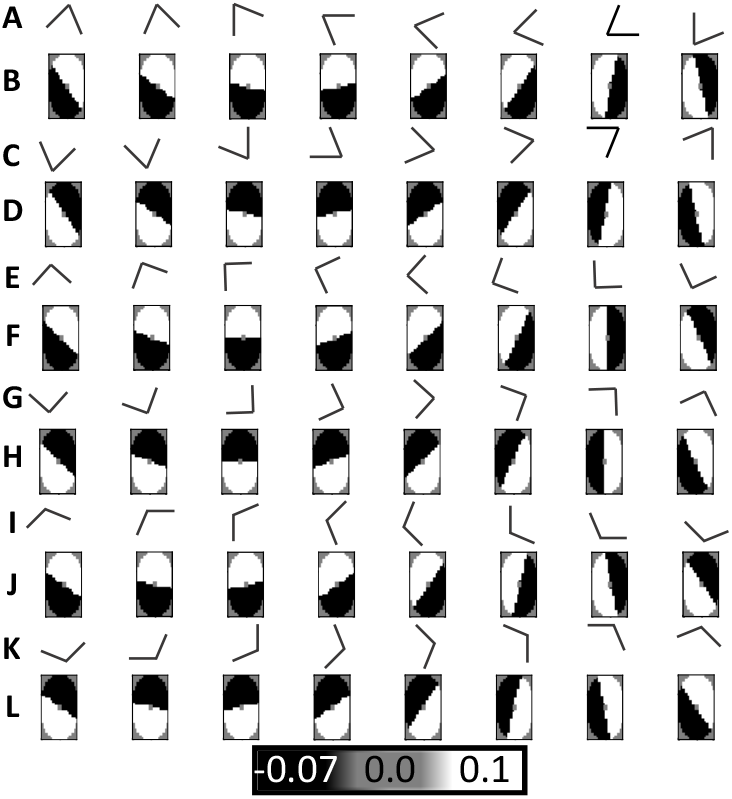
Afferent Connections for model V4 L2/3 neurons selective to mostly convex high curvatures throughout the receptive field. **A & C**: Pictorial representation of the preferred stimuli of V2 afferents inputs to V4 having a 67.6 ° interior angle at a complete range of rotations every 22.5 °. Details of V2 neuron selectivity are described in the Methods 4.4.3. **E & G**: Pictorial representation of the preferred stimuli of V2 afferents inputs to V4 having a 90 ° interior angle at a complete range of rotations every 22.5 °. **I & K**: Pictorial representation of the preferred stimuli of V2 afferents inputs to V4 having a 112.5 ° interior angle at a complete range of rotations every 22.5 °. **B, D, F, H, J, L**: The synaptic connection strength versus position in topographic space relative to the soma at the center for a V4 neuron selective to high curvature throughout the receptive field. Each square is a single 16 ×28 convolution filter implementing the weights for the V2 afferent inputs indicated by the pictogram above. White=excitatory, Black=Inhibitory, Gray=No connection. All of the weights in B-L together are the connections to the topographically arranged inputs for one V4 neuron. There are excitatory connections to every orientation of the V2 afferents, and the position of the excitation rotates around the receptive field with the orientation of the inputs so that convexly oriented inputs are excited and other inputs are inhibited.

### 2.2 Medium Curvature Selectivity

#### 2.2.1 Composite Boundary Stimuli

[Pasupathy and Connor, 2001] found that some V4 neurons are selective to a medium degree of curvature at specific positions relative to the center of the receptive field, and that while some units were selective to curvature that was convex to the center of the receptive field others were selective for curvature that is concave to the center of the receptive field. Figure 7A and 7B are reproduced from [Pasupathy and Connor, 2001] and each shows the response of a different identified unit to the boundary based stimuli set. See the beginning of section 2.1.1 for a detailed description. Figure 7A represents the recorded selectivity of a neuron described as a “V4 neuron tuned for concave curvature at the right” which describes the common element among those composite boundaries to which it responds. Figure 7B represents the recorded selectivity of a neuron described as “V4 neuron tuned for broad convex curvature at the top.”.

**Fig. 7.**
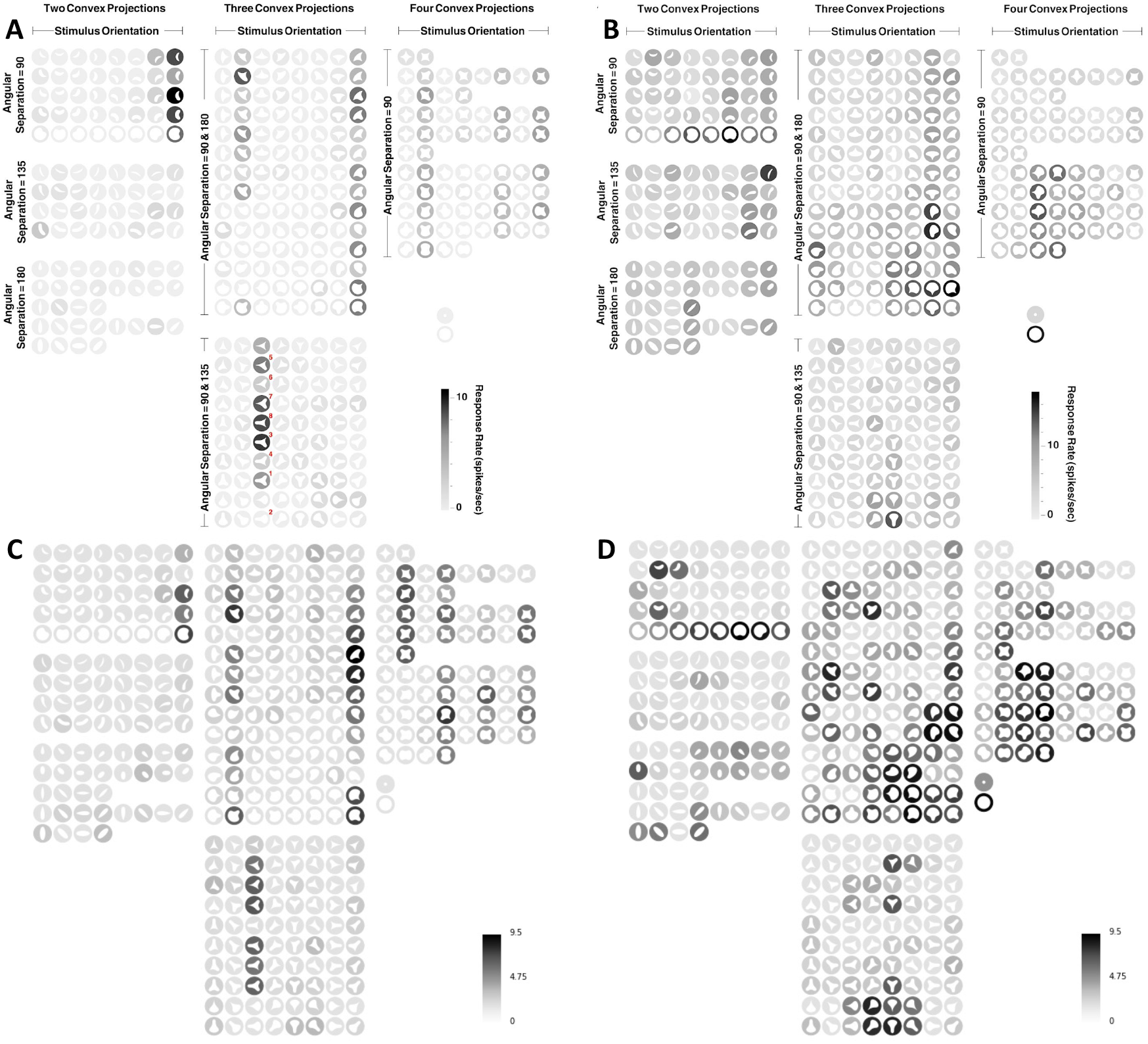
Recording and simulation of Medium curvature selectivity in V4: The white shape in each location represents the the stimuli shape, which is actually presented as a black silhouette on a white background. The shading of the disc around the shape represents the intensity of the response as indicated by the legend. **A**Recording reproduced from [Pasupathy and Connor, 2001] showing a V4 neuron tuned for concave curvature at the right. **B** Recording reproduced from [Pasupathy and Connor, 2001] showing a V4 neuron tuned for broad convex curvature at the top. **C** Result of a model simulation qualitatively matching **A**. The results shown are the number of spikes out of N simulations. **D** Result of a model qualitatively matching **B**.

The corresponding model simulations reproducing the medium curvature selective unit recordings are shown in Figures 7C and 7D. The model simulation matches the selectivities of the real neurons qualitatively, but not exactly. For the concave curvature selective unit (C), the selectivity is very close but the magnitude of responses are somewhat unequal.^1^ For the convex curvature selective unit, the model response is stronger at a broader range of stimuli with medium convex curvature at the top and has somewhat less response to other less related stimuli. The model was manually tuned based on the published image and without the data used to produce Figure 7A and 7B, as described in the Methods.

The model shows that a simple, systematic arrangement of afferent connections captures most of the properties of the selectivity: The convolution filters are the model embodiment of the connection patterns that give rise to selectivity for medium curvature in a particular position. The convolution filters resulting in the selectivity of Figures 7C and 7D are shown in Figure 8. Figure 8 A-F show the connection pattern for the neuron tuned to concave curvature at the right, and Figure 8 G-L show the connection pattern for the neuron tuned to broad convex curvature at the top. The selectivity of the afferent inputs to the V4 L2/3 neurons are represented by pictograms in Figure 8: These are the outputs of V2 neurons selective to corners having interior angles 90^°^ (A,G), 112.5^°^ (C,I) and 145^°^ (E,K). The corresponding convolution filter weights are shown below each pictogram, Figure 8 B,D, F (concave at the right) and H, J and L (broad convex at the top). For a complete explanation of the meaning of each part of the Figure, see the text accompanying Figure 4.

**Fig. 8.**
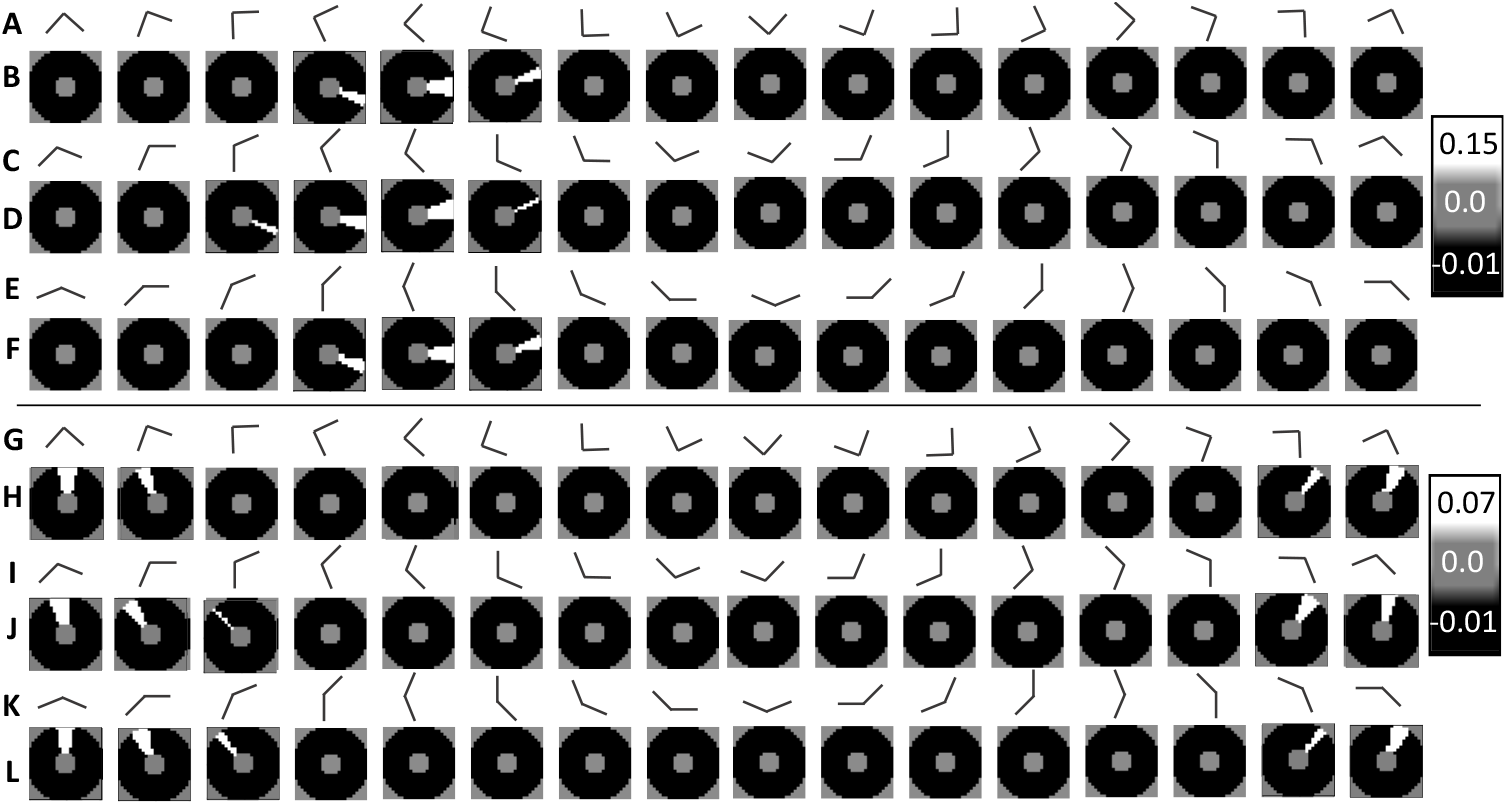
Afferent Connections for V4 neurons selective to medium curvatures in asymmetric zones. **A - F**: Connections for the model V4 neuron selective to broad convex curvature at the top. **G - L**: Connections for the model V4 neuron selective to concave curvature at the right. **A, C & E**: Pictorial representation of the preferred stimuli of V2 afferents inputs to V4 having a 90 ° (A), 112.5 ° (C) and 145 ° (D) interior angles at a complete range of rotations every 22.5 °. Details of V2 neuron selectivity are described in the Methods 4.4.3. **B, D & F**: The synaptic connection strength versus position in topographic space relative to the soma at the center for a V4 neuron selective to broad convex curvature at the top. Each square is a single 9×9 convolution filter implementing the weights for the V2 afferent inputs indicated by the pictogram above. White=excitatory, Black=Inhibitory, Gray=No connection. All of the weights in A-F together are the connections to the topographically arranged inputs for one V4 neuron. There are excitatory weights on the inputs that are convexly oriented in the top part of the receptive field; other afferent inputs are weakly inhibitory. The orientation of inputs that are excitatory rotates with position around the receptive field. **G, I & K**: Pictorial representation of the preferred stimuli of V2 afferents inputs to V4 having a 90 ° (G), 112.5 ° (I) and 145 ° (K) interior angles at a complete range of rotations every 22.5 °. **H, J & L**: The synaptic connection strength versus position in topographic space relative to the soma at the center for a V4 neuron selective to concave curvature at the right, as in B/D/F. There are excitatory weights on the inputs that are concavely oriented in the right part of the receptive field; other afferent inputs are weakly inhibitory. The orientation of inputs that are excitatory rotates with position around the receptive field.

For the neuron selective to concave curvature at right, the excitatory connections are to right opening corner selective afferents. For the neuron selective to broad convex curvature at the top, the excitatory connections are to downward opening corner selective afferents. In both cases, there are excitatory connections to a small range of orientation of afferents in a wedge shaped zone outside the center of the receptive field. The precise orientation selectivity of the afferent inputs rotates as the position of the inputs rotates around the receptive field. The overall preferred stimuli set for the pattern were illustrated in Figure 1V: it is a set of medium curvature overlapping curves in a particular region of the receptive field.

#### 2.2.2 Grid of Composite Bar Stimuli

[Nandy et al., 2013] also found that some V4 units are selective exclusively to medium curvature. And [Nandy et al., 2013] found that some units selective to medium curvature had such selectivity over a broad range of positions, and the orientation of the preferred curvature varied with position around the receptive field. The recording from [Nandy et al., 2013] of a medium curve selective V4 L2/3 unit with selectivity through its receptive field is shown in Figure 9 and compared to a model. For a detailed explanation of the parts of Figure 9 see the text accompanying Figure 5 at the beginning of Section 2.1.2. For both the model and recording the receptive field is highly varied over multiple stimuli locations. There is strong response to medium curvatures, though also broad tuning that includes activation to straight and highly curved stimuli. The orientation of the most preferred stimuli are convex around the center of the receptive field. The main shortcoming of the model to reproduce the recording is that the orientations of some of the preferred stimuli are different by around 10^°^-20^°^, and also the model lacks background firing to the less preferred stimuli found in the recordings.

**Fig. 9.**
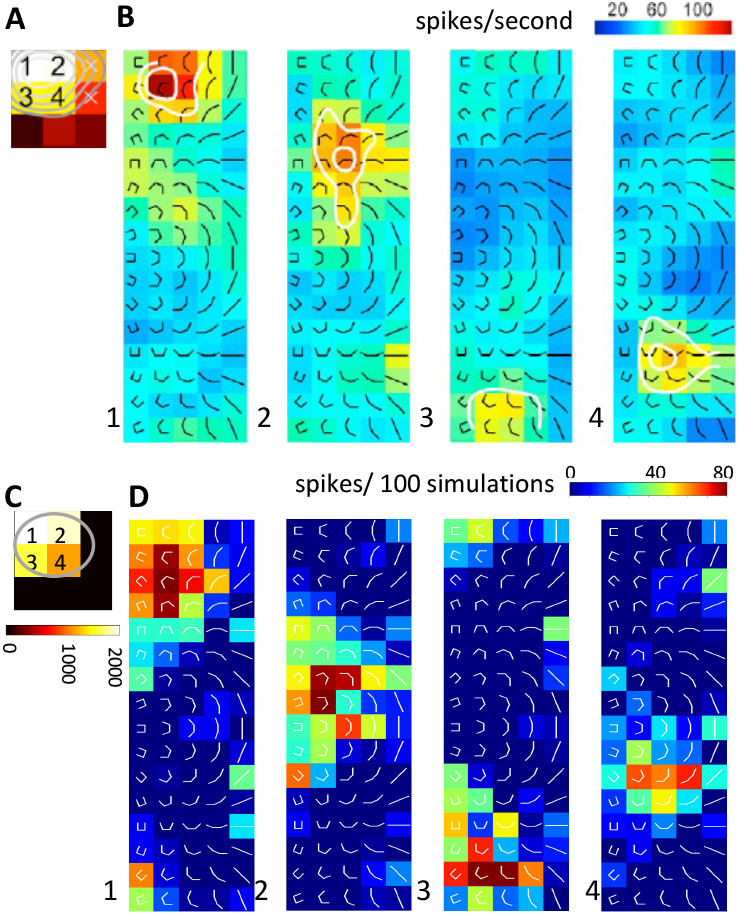
Medium curvature selective unit recording (top) and simulation (bottom) **A** Receptive field for recording. **B** Shape tuning for recording, 1-4 show the shape tuning for curvature (x-axis) and orientation (y-axis) at indicated positions in the receptive field (A). **C** Receptive field for simulation. The approximate location and extent of the simulated neurons dendritic tree in the topographic space of the afferent inputs is shown. **D** Shape tuning for simulation, as in B, at positions in the simulated receptive field (C). The simulation qualitatively reproduces the most significant features of the medium curvature selective neuron recording.

The connection pattern to affrent inputs for for the [Nandy et al., 2013] medium curvature selective V4 L2/3 cell model is shown in Figure 10. The afferent inputs to the V4 L2/3 neuron are represented by pictograms Figure 10 A,C, E, G, I and K: These are the outputs of V2 neurons selective to corners having interior angles 90 ° (A,C), 112.5 ° (E,G), and 145° (I,K). The corresponding convolution filter weights are shown below each pictogram, Figure 10 B,D, F, H, J and L. The pattern is that for every type of afferent input, a wedge covering around 20% the receptive _eld is excitatory and the remainder is inhibitory, and a small region at the center is neutral. The direction of the excitatory wedge is such that the excitatory afferent inputs are approximately concave to the center of the receptive field. To achieve this, the excitatory wedge boundary rotates around the receptive field, depending on the orientation of the afferent inputs. Non-preferred orientations are inhibited and afferent inputs with both lower interior angle (more acute) and higher interior angle (more straight) complex shape preferences are ignored. The overall preferred stimuli set for the pattern were illustrated in Figure 1W: it is a set of medium curvature overlapping curves, convex to the center all around the receptive field. For a detailed explanation of the meaning of each part of Figure 10, see the text accompanying Figure 4 in Section 2.1.1. For details of the how the connection pattern for the V4 neurons selective to medium curvature were constructed see the Methods Section 4.4.4.

**Fig. 10.**
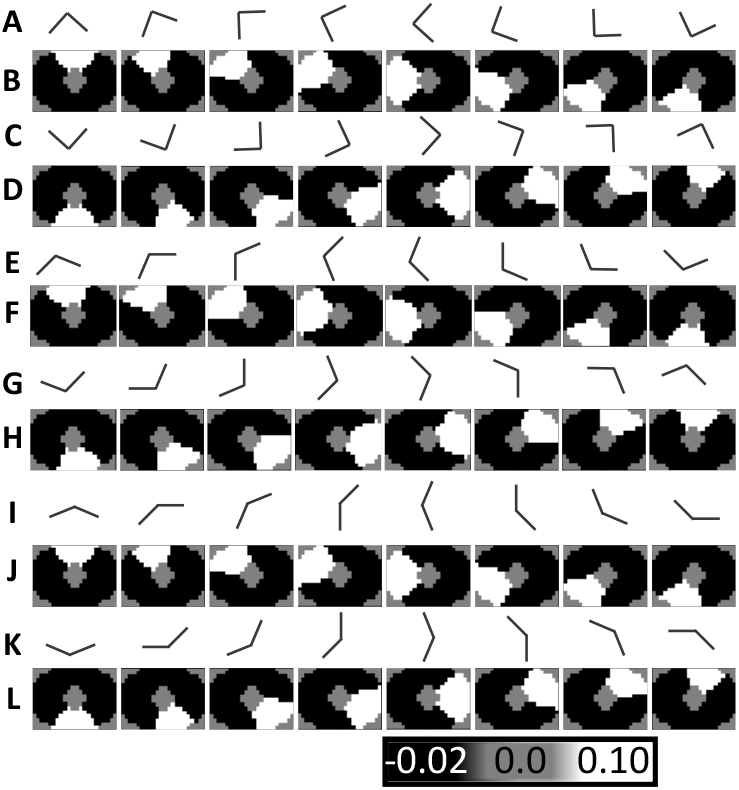
Afferent Connections for model V4 L2/3 neurons selective to convex medium curvatures throughout the receptive field. **A & C**: Pictorial representation of the preferred stimuli of V2 afferents inputs to V4 having a 90^°^ interior angle and a complete range of rotations every 22.5^°^. Details of V2 neuron selectivity are described in the Methods 4.4.3. **E & G**: Pictorial representation of the preferred stimuli of V2 a_erents inputs to V4 having a 112.5 ^°^ interior angle and a complete range of rotations every 22.5^°^. **I & K**: Pictorial representation of the preferred stimuli of V2 afferents inputs to V4 having a 145 ^°^ interior angle and a complete range of rotations every 22.5^°^. B, D, F, H, J, L: The synaptic connection strength versus position in topographic space relative to the soma at the center for a V4 neuron selective to medium convex curvature throughout the receptive _eld. Each square is a single 20 × 24 convolution filter implementing the weights for the V2 afferent inputs indicated by the pictogram above. White=excitatory, Black=Inhibitory, Gray=No connection. All of the weights in B-L together are the connections to the topographically arranged inputs for one V4 neuron. There are excitatory connections to every orientation of the V2 afferents, and the position of the excitation rotates around the receptive field with the orientation of the inputs so that convexly oriented inputs are excited and other inputs are inhibited.

### 2.3 V4 Low Curvature Selective Model

[Nandy et al., 2013] also found that some V4 units are selective to low curvature or straight stimuli with a consistent orientation tuning throughout the receptive field - for these V4 neurons, the selectivity was noted to be invariant with position in the receptive field. Figure 11 compares the shape tuning of the V4 L2/3 neuron model with the recording. For a detailed explanation of Figure 9 see the text accompanying Figure 5 at the beginning of Section 2.1.2. For both recording and simulation, the most signi_cant activity for the unit occurs at 3 stimuli locations. The most preferred stimuli are straight or minimally curved and at a small angle counter clockwise from horizontal. However, there is broad tuning around the most preferred stimulus and non-zero activity as high as 50% of the maximum for stimuli that are at orientations as much as 22.5^°^ away and at medium curvatures.

**Fig. 11.**
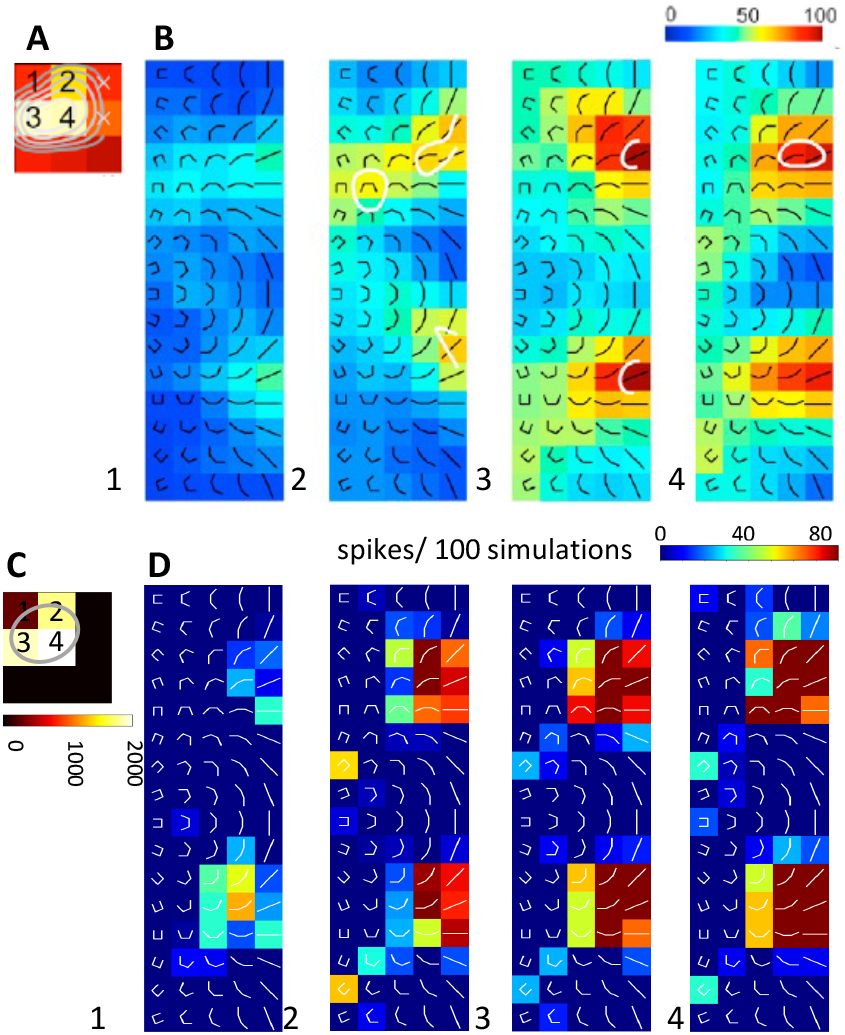
Low curvature selective unit recording (top) and simulation (bottom) **A** Receptive field for recording. **B** Shape tuning for recording, 1-4 show the shape tuning for curvature (x-axis) and orientation (y-axis) at indicated positions in the receptive field (A). **C** Receptive field for simulation. The approximate location and extent of the simulated neurons dendritic tree in the topographic space of the afferent inputs is shown. **D** Shape tuning for simulation, as in B, at positions in the simulated receptive field (C). The simulation qualitatively reproduces the main features of the low curvature selective neuron recording.

The main deficiency of the model in reproducing the recording is that the model does not have very weak responses (25% of the maximum) to as wide a range of orientation and curvatures as is seen in the recording. Also, the model neuron does not have any responses in the stimuli locations outside of top 4, while the recordings show weak responses.

The connect patterns embodied by convolution filters for the unit that produces the simulation result in Figure 11 is shown in Figure 12. The afferent inputs to the V4 L2/3 neuron are represented by pictograms Figure 12 A,C and E: These are the outputs of V2 neurons that have “ultra-long” selectivity (A) and complex selectivity at very high interior angles of 157.5 ^°^ (nearly straight). The corresponding convolution filter weights are shown below each pictogram, Figure 12 B,D, and F. The connection pattern is to have the strongest connections to afferent inputs from ultra-long units with an orientation of 22.5 ^°^ from horizontal, but also with broad tuning including orientations from 0 ^°^ to 45 ^°^. Afferents for nearly straight complex units make excitatory connections at when the orientation of the selectivity is close to 22.5 ^°^ from horizontal only. The connections to the nearly straight selective V2 afferents are at a lower strength than the most preferred orientation ultra-long units. Afferents from inputs selective for other orientations of nearly straight stimuli are accepted from inhibitory L4 units, as shown in Figure 12D, while afferents from inputs selective for more acute angled complex-shaped units are ignored.

**Fig. 12.**
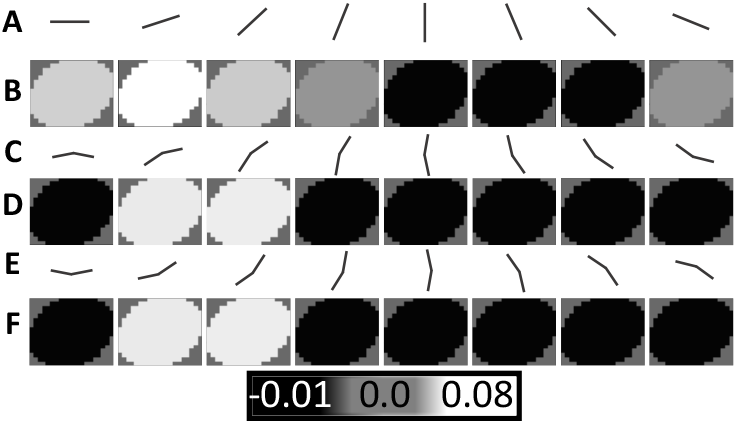
Afferent Connections for model V4 L2/3 neurons selective to low curvatures throughout the receptive field. **A**: Pictorial representation of the preferred stimuli of V2 afferent inputs to V4 having selectivity to “ultra-long” straight lines at a range of orientations. **C & E**: Pictorial representation of the preferred stimuli of V2 afferents inputs to V4 having a 157.5^°^ interior angle at a complete range of rotations every 22.5^°^. Details of V2 neuron selectivity are described in the Methods 4.4.3. **B, D, F**: The synaptic connection strength versus position in topographic space relative to the soma at the center for a V4 neuron selective to low curvature throughout the receptive field. Each square is a single 15 × 20 convolution filter implementing the weights for the V2 afferent inputs indicated by the pictogram above. White=excitatory, Black=Inhibitory, Gray=No connection. All of the weights in B-F together are the connections to the topographically arranged inputs for one V4 neuron. The strongest excitation comes from afferents selective to ultralong configurations at an angle 22.5^°^ above horizontal, and weaker excitation from afferents selective to ultralong inputs at nearby angles. There are also nearly as strong weights on the afferents selective to 157.5^°^ interior angle corners at an angle of 22.5^°^ above horizontal.

In comparison to the model neurons for V1 and V2, the V4 model neuron has a relatively large receptive field: 15x20 in the reduced frame of V4, equivalent to 60x80 in the original 112 × 112 pixel visual field model. (In comparison, the V1 neurons have receptive fields of 6 × 9 and the V2 neurons have fields of 18 × 18 in the frame of V1.) This large receptive field is required to match the recordings, as shown in Figure 11 and the model does not match the weak secondary stimulation.

Due to the large receptive field, the total excitation in the V4 L2/3 units is much larger than in the V1 and V2 connection patterns. The total excitation for the low curvature selective V4 L2/3 neuron model is approximately 164, in units of the neurons threshold, while the total inhibition is 52. The high amount of excitation is required to match the recordings: As can be seen from Figure 11, a small stimulus in a fraction of the neurons receptive field is sufficient to cause a spike. Because the response is relatively uniform in the receptive field, it follows that there must be many times more excitative inputs available than what is required to respond to a small stimulus. The preferred stimuli were illustrated in Figure 1I: it is a classic parallel line grating, but with sone caveats: it allows imperfectly straight lines, and it requires a matching stimuli in only a small portion of the receptive field to trigger a spike. For more details of the how the connection pattern was construction see the Methods Section 4.4.4.

### 2.4 PSP Simulation Dynamics

Figure 13 illustrates the dynamics of PSP summation at the soma for the model in several illustrative neurons. Figure 13A shows examples of Pyramidal model neurons in V1. The time constants for the model are based on the measurements of [Holmgren et al., 2003] and shown in Table 1. [Holmgren et al., 2003] found that PSP’s in Pyramidal neurons have duration that is relatively long compared to the time scale of spike synchronization : The 10-90% rise time averages 4.5 ms and the width at half amplitude is 30 ms. In comparison, spikes arrive in V1 from the LGN in synchronized bursts on the order of a few ms [Singer, 1999]: As described in detail in the Methods, these arrival times are modeled with a gamma distribttion with a standard deviation of 2 ms. As a result, the PSP summation trajectories illustrated in 13A (from V1 model pyramidal neurons) appear fairly contiguous as summation and threshold crossings occur before any substantial decay of the individual PSP’s. Note that according to [Holmgren et al., 2003] the PSP time constants of inhibitory to pyramidal connections are substantially similar to those of pyramidal-pyramidal connections; for the model of this paper these time constants were made the same.

**Table 1.** PSP Time Constants (ms) and properties for Figure 13.

| Sub-Figure | $\tau_{rise}$ | $\tau_{fall}$ | 10-90% rise time | Width at half-amplitude | Source |
| --- | --- | --- | --- | --- | --- |
| A | 4.5 | 30 | 5.3 | 36 | [Holmgren et al., 2003] Pyr-Pyr |
| B | 1.9 | 13 | 2.2 | 14.6 | [Holmgren et al., 2003] Pyr-FS |
| C | 0.25 | 1 | 0.2 | 2.6 | NA |

**Fig. 13.**
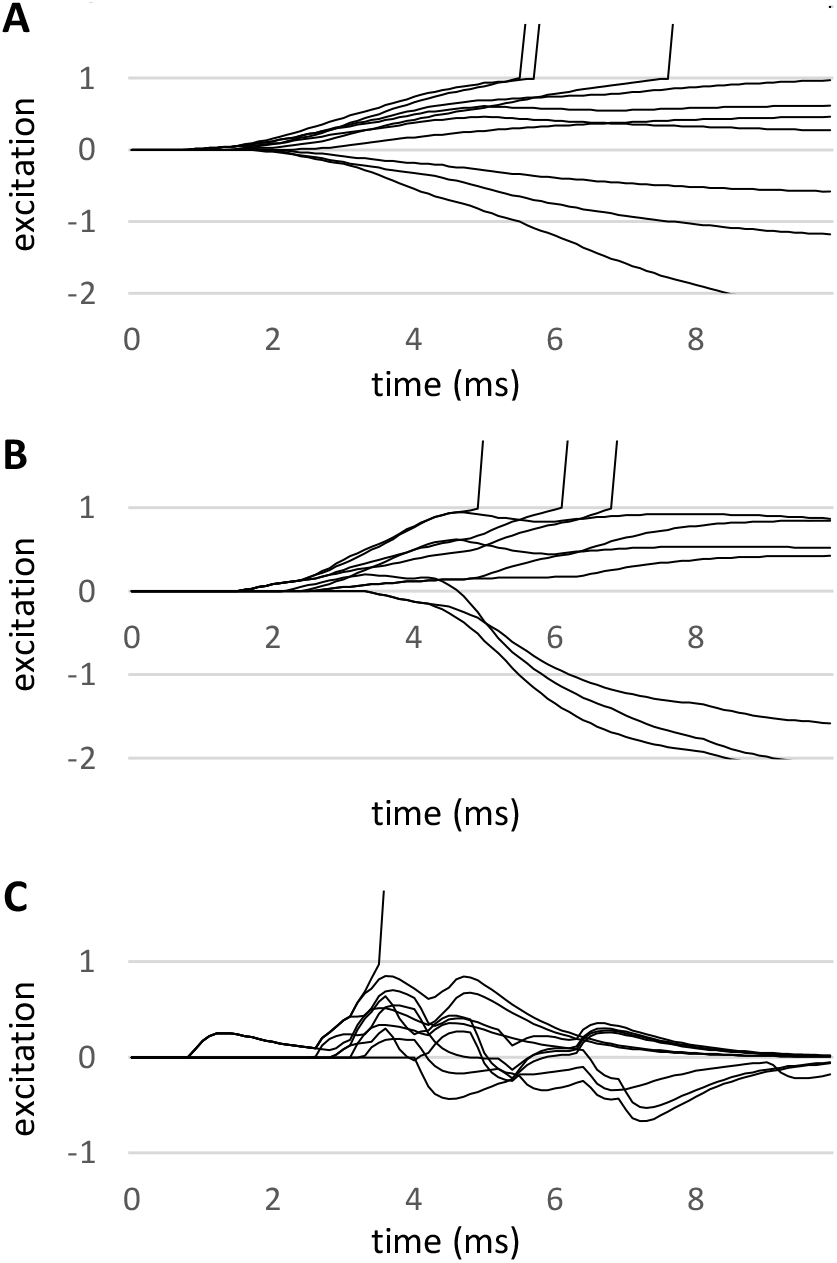
Depolarization Dynamics **A** Parameters of the model L2/3 Pyramidal and L4 neurons **B** Parameters of the model inhibitory neurons, which are modeled as “Fast Spiking”. **C** Test of faster dynamics.

Figure 13B illustrates the PSP’s for the time constants of the PSP’s in a model inhibitory neuron, received from a pyramidal neuron, in accordance with the result in [Holmgren et al., 2003]. These are “fast spiking” neurons and have significantly shorter PSP time constants: the 10-90% rise time is just 2.2 ms and the width at half amplitude is 14 ms. For the inhibitory neuron time constants the inflections in the trajectory due to the addition of excitation or subtraction of inhibition are somewhat more apparent but the PSP time scale is still long compared to the summation. Note that for illustrative purposes these model neurons of Figure 13B include inhibitory inputs: The actual inhibitory neurons in the model have only excitatory inputs and the trajectories are less able to illustrate the time constant properties.

In order to confirm that the model could reproduce more dynamic PSP trajectories, additional simulations were performed using time constants faster than those reported in [Holmgren et al., 2003]. The results of these simulations are shown in Figure 13C; for these simulations the 10-90% rise time was decreased to 0.2 ms and the width at half amplitude reduced to 2.6 ms. Also, for illustrative purposes the synchronization of the input spikes was decreased: the arrival time gamma distribution had a 3 ms standard deviation instead of 2 ms. With these unnatural time constants it is clear that the model can reproduce much more rapid PSP dynamics. However, this is for illustrative purposes and not applicable to the cortical model described in this paper.

While there is no question of the model’s ability to compute the time course of faster PSP dynamics, [Holmgren et al., 2003] suggests that the reality is a slower dynamics relative to the precise input time synchronization. The PSP time constants are also slow relative to the presumed frequency of cascades: As summarized in [Buzsáki and Wang, 2012], cortical neurons engaged in gamma timescale activity typically have active bursts every 10-30ms. With such a slow time constant of decay, width at half amplitude 30ms for the excitatory and 14ms for the inhibitory neurons, there is not enough time for passive decays to return the neuron to a resting state. For these reasons it is hypothesized that the networks responsible for gamma oscillations include inhibition to return the cortical neurons back to their resting state more quickly. However, this level of detail is beyond the scope of this modeling study, focused on the feedforward cascade as it is.

### 2.5 Spike Propagation

The distribution of spike times in each layer of the model is illustrated in Figure 14**A**, in response to one of the stimuli composed of convex and concave boundaries from [Pasupathy and Connor, 2001] (Figure 14**B**). As described in the Methods, the spike transmission time between layers are drawn from gamma distributions designed to match typical intra- and inter-cortical propagation delays (See Methods Section 4.1.1 and Table 5.) Under these assumptions it takes around 25 ms for a spike wave to propagate from the model V1 L4 input layer to V4 L2/3. The results are in agreement with [Girard et al., 2001] which shows orthodromic latency between V1 and V2 neurons of just around 2-3 ms and [Plomp et al., 2017] which shows inter-laminar latencies between L4 and L2/3 of around 1 ms. It is also consistent with the overall picture presented in [Thorpe and Imbert, 1989] and [VanRullen et al., 2005] that each cortical area takes around 10 ms to perform its role in a rapid feedforward cascade that leads to visual recognition.

**Fig. 14.**
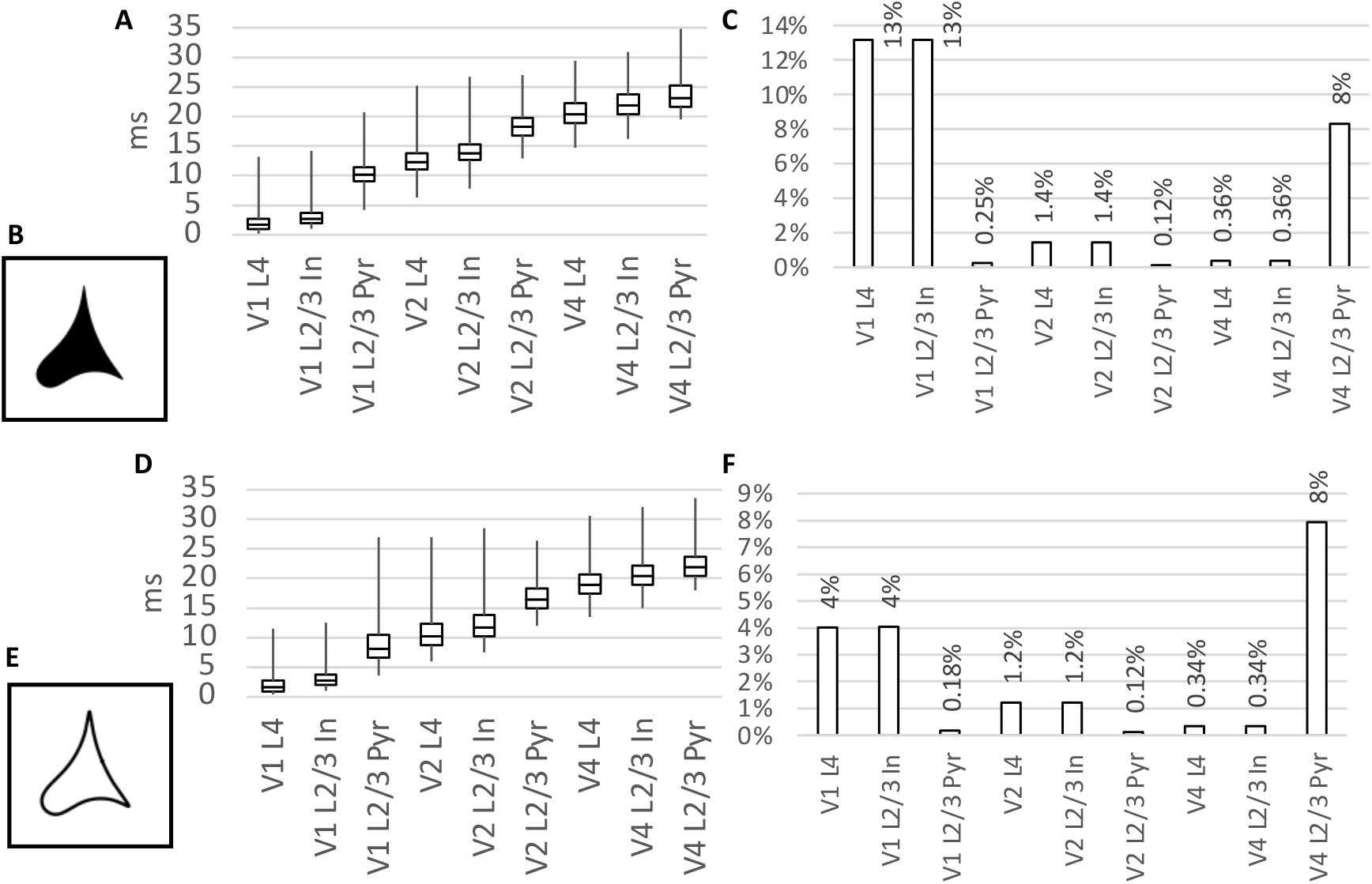
Spike Propagation **A** Spike time distribution in the model for a single stimulus from [Pasupathy and Connor, 2001] (shown as **B**) Box-plots show the median and upper/lower quantiles, and the whiskers show the 1st/99th percentiles. **C** Percentage of neurons spiking in each layer for the stimulus **B. D** Spike time distribution in the model for a single outline stimulus based on [Pasupathy and Connor, 2001] (shown as **E**) **F** Percentage of neurons spiking in each layer for the stimulus **E**.

The simulations of Figure 14 were run for 36 ms, although it was found that more than 99% of spikes that will arrive have arrived by around 27.5ms. The time limit for the simulations shown in Figures 3 though 11 was set to 30 ms to capture the vast majority of spikes in a reasonable length simulation.

Figure 14**C** shows the percentage of neurons spiking in each layer in response to the stimuli. While around 10% of the model units in the first V1 and final V4 layers spike, in the interior layers spiking is much sparser: only 1% or fewer units fire. Note that for this test the final layer includes both the model units created to match the results of [Pasupathy and Connor, 2001] (Figures 4 and 8) and [Nandy et al., 2013] (Figures 6, 10 and 12), and further includes all of these units at a complete set of rotations as described in the Methods.

Figure 14**D** through **F** shows the same tests as Figure 14**A** through **C**, but in this case one of the convex and concave boundaries stimuli from [Pasupathy and Connor, 2001] is rendered with an outline. This stimuli shows significantly less spikes in the first layer and slightly faster propagation of the signal to later layers. Otherwise the results are substantially the same. More details of tests on outlines of convex and concave boundary stimuli are shown in Section 2.6.3.

### 2.6 Simulated Responses to Novel Stimuli

With the model tuned to reproduce results from [Pasupathy and Connor, 2001] and [Nandy et al., 2013], it can be used to predict results for novel stimuli. One small example of this was already presented in Figure 14**E**: While [Pasupathy and Connor, 2001] used silhouette contours, the stimuli of [Nandy et al., 2013] are lines - Figure 14**E** is a blend of the two approaches, making a contour boundary stimuli as an outline. Model experiments like this can be used to verify the model in future neurophysiological experiments, improve future models with more specific constraints, suggest useful stimuli for future experiments, and help inform hypotheses about the function of V4.

#### 2.6.1 Swapping Units and Stimuli from Past Studies

The V4 units selected for presentation in [Pasupathy and Connor, 2001] and [Nandy et al., 2013] were different: [Pasupathy and Connor, 2001] focused on neurons selective for particular degrees of curvature in particular zones of the receptive field, while [Nandy et al., 2013] focused on neurons with selectivity for a degree of curvature throughout the receptive field. In each case, the stimuli presented are well suited to illustrate the properties of the neurons they selected. How the neurons of one study would have responded to the stimuli of the other is an interesting question.

Figure 15 illustrates what response could be expected if the units presented in [Pasupathy and Connor, 2001] were stimulated with the approach of [Nandy et al., 2013]. For the neurons selective to acute curvature the response, shown in Figure 15A and 15B, was weak and mainly to straight line stimuli and not to curves. This is because the stimuli of [Nandy et al., 2013] do not contain angles more acute than 90^°^, but as illustrated in Figure 4 the model neuron selective for acute convex curvature are optimally stimulated by stimuli at an angle of 45^°^. So the responses in Figure 15 A and B are a secondary response to nearly straight stimuli that are aligned with the excitable portion of the receptive field. In this case the stimuli of [Nandy et al., 2013] seems inadequate to capture the full variability of V4 neurons.

**Fig. 15.**
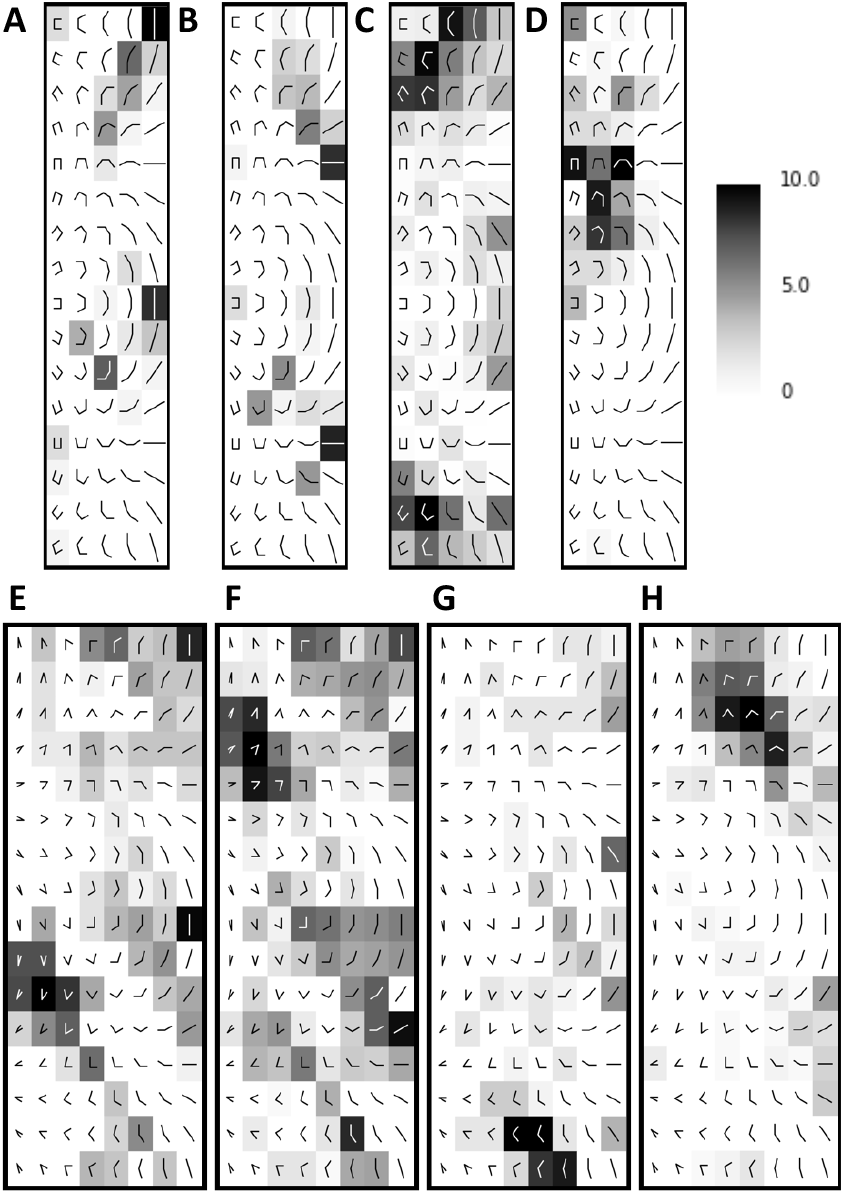
Simulated response of neurons from [Pasupathy and Connor, 2001] to stimuli of [Nandy et al., 2013], **A**-**D**, and a novel stimuli set of corner stimuli, **E**-**H**. Each pictogram represents a composite bar stimuli that varies by curvature (x-axis) and orientation (y-axis); the gray scale coded scale shows the response firing rate. For a complete description of the stimuli from [Nandy et al., 2013] see Figures 2 and the text accompanying Figure 5. The stimuli of corners are constructed in a similar fashion, albeit with two bars, as described in the Methods Section 4.6.2. **A & E** The neuron of Figures 3A and 4 selective to for acute convex curvature at the bottom left. **B & F** The neuron of Figure 3B selective for acute convex curvature at the top right. **C & G** The neuron of Figure 7A tuned for concave curvature at the right. **D & H** The neuron of Figure 7B tuned for broad convex curvature at the top.

The neurons selective to medium curvatures in [Pasupathy and Connor, 2001] respond well to the stimuli of [Nandy et al., 2013] in a manner matching the selectivity to the stimuli of [Pasupathy and Connor, 2001]. This is illustrated in Figure 15C for the V4 neuron tuned for concave curvature at the right, and Figure 15D for the V4 neuron tuned for broad convex curvature at the top.

Figure 16 illustrates what response could have been expected if the neurons presented in [Nandy et al., 2013] were stimulated with the approach of [Pasupathy and Connor, 2001]. In this case, all of the neurons responded very strongly to the stimuli. With the parameters tuned to match the results of [Nandy et al., 2013], the model neurons responded so vigorously to almost all stimuli that there was no differentiation in the shape tuning. To continue the experiment, the excitation threshold in the model the neurons was raised to the point where there was a varied range of responses across stimuli - this can be viewed as a form of adaptation by the model neurons albeit manual.

**Fig. 16.**
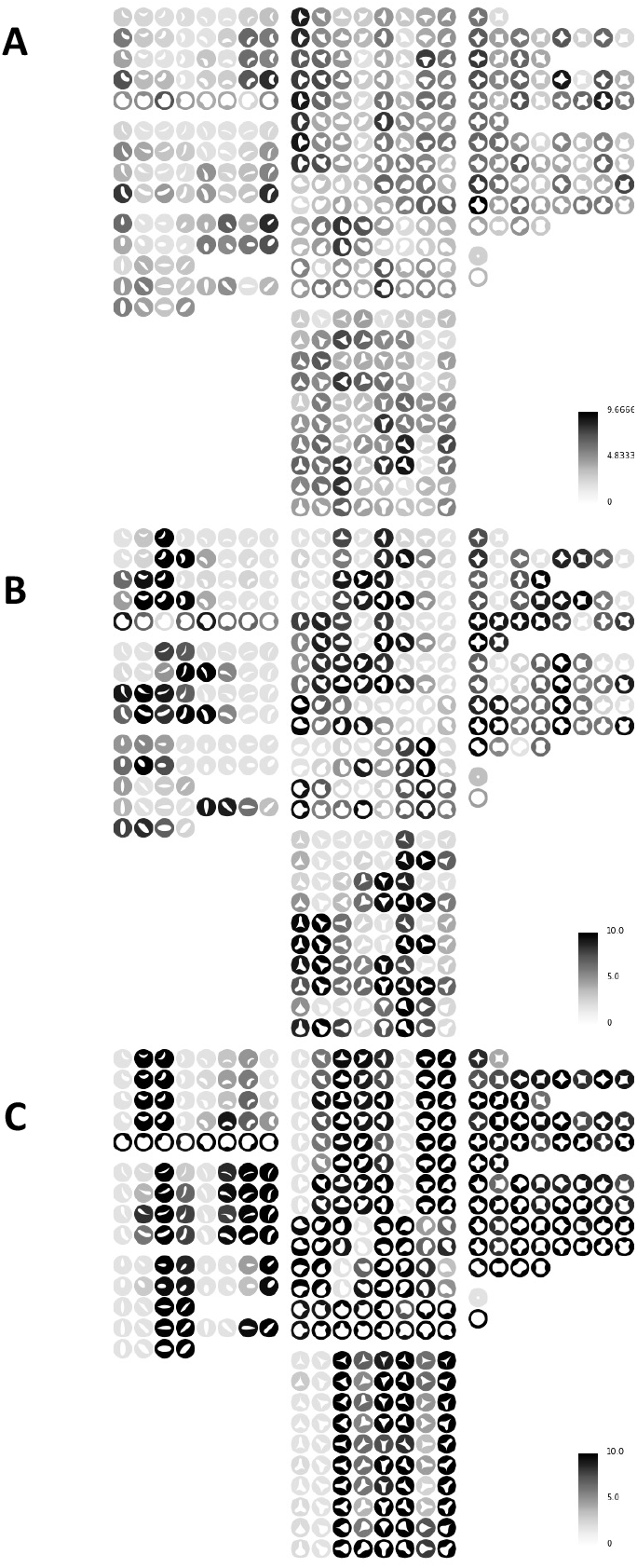
Simulated response of neurons from [Nandy et al., 2013] to stimuli of [Pasupathy and Connor, 2001]. See Figure 3 for details. **A** The neuron of Figure 5 and 6 selective for high curvature. **B** The neuron of Figure 9 and Figure 10 selective for medium curvature. **C** The neuron of Figure 11 and Figure 12 selective for low curvature.

However, even after raising the threshold of the model neurons based on [Nandy et al., 2013] to the point where the stimuli of [Pasupathy and Connor, 2001] exhibit a full range of responses, so many of the stimuli elicited strong responses that it is still difficult to discern clear patterns. The response of the high curvature selective unit to the boundary contour stimuli is illustrated in Figure 16A: It tends to respond to shapes that have at least some acute angles, but the pattern is irregular and inconsistent - all the shapes triggering strong response have some acute angle, but not all shapes with acute angles trigger a strong response. This is due to the fact that the shape of the receptive field (Figure 6) is elongated on the vertical dimension and some stimuli do not lie evenly inside the receptive field. The response of the medium curvature selective unit is shown in 16B and a similar characterization applies: It tends to favor shapes with some broad, non-acute curves in some part of the shape. But the response is uneven at different orientations. The response of the low curvature selective unit is illustrated in Figure 16C. Knowing that the neuron is selective to nearly straight lines at an angle that is approximately 22.5^°^ above horizontal (as illustrated in Figure 12) it is fairly apparent that every shape to which the neuron has at least some boundary that matches. But the selectivity to straight and nearly straight lines at a particular orientation is not as apparent in the contour boundary stimuli of [Pasupathy and Connor, 2001] as in the grid of curves stimuli from [Nandy et al., 2013].

#### 2.6.2 Response to a Grid of Corner Stimuli

The comparisons in the precceeding section suggests that both the contour boundary stimuli of [Pasupathy and Connor, 2001] and the grid of curves stimuli of [Nandy et al., 2013] have certain blind spots in characterizing the response of V4 units that the other covers well. A better stimuli set for characterizing all of these units may be one that maps the orientations and acuteness of the selectivity on a grid like in [Nandy et al., 2013] but includes more acute angled stimuli. For example, each stimuli could consist of just two bars at a full range of angles from fully straight to highly acute. If the hypothesis that V2 complex selective neurons are well described by having corners as their preferred stimuli, then a stimuli of corners will be an optimal stimuli for representing V4 selectivity as it forms a basis (in the mathematical sense) spanning the independent components that may contribute to the V4 neuron response.

Hypothetical results from a stimuli set of corners are shown in Figure 15E-H for the model neurons based on [Pasupathy and Connor, 2001]. The stimuli of corners are constructed in a similar fashion to the three bar curvature stimuli, albeit with two bars, as described in the Methods Section 4.6.2. This stimuli set triggers a robust response in the acute selective neurons from [Pasupathy and Connor, 2001] and captures the specific range and direction of curvature to which those neurons respond (Figure 15E and F). It also shows the extent to which the neurons respond to a range of corner stimuli that are close to the preferred orientation and interior angle The corner stimuli set also adequately captures the neurons responsive to concave curvature (Figure 15G) medium convex curvature (Figure 15H).

The simulated response of neurons from [Nandy et al., 2013] to stimuli of oriented corners is shown in the Appendix Figure A4. As the stimuli are substantially similar to the stimuli of [Nandy et al., 2013], the results are very similar to Figures 5, 9 and 11. The only differences is it reveals that the high curvature selective neuron is also selective to 57.5 ^°^ corners, but is hardly stimulated by 45 ^°^ corners.

#### 2.6.3 Response to Outlines of Convex and Concave Boundary Stimuli

As already illustrated in Figure 14, a novel stimuli set can be created by blending the characteristics of the stimuli in [Pasupathy and Connor, 2001] and [Nandy et al., 2013]: Contour boundary stimuli that are presented as outlines instead of silhouettes. For another example, see Figure 18A and B. Simulations were performed for the complete data set of [Pasupathy and Connor, 2001] presented as outlines, instead of as silhouettes. The results of this experiment are illustrated in Figure 17: the responses of the simulated V4 neurons to the outline are qualitatively similar to the standard silhouette. In every case the main shapes responded to are similar, but are not entirely the same. The neuron selective to concave shape at the right (Figure 17C) responds somewhat more to the outline stimuli (right) than the silhouette. Also, the neuron selective to broad concave curvature at the top (Figure 17D) responds somewhat more to the silhouette stimuli (left). For the neurons responsive to acute convex (Figure 17A and B) it appears that the responses are more equal in overall response but still not precisely the same.

**Fig. 17.**
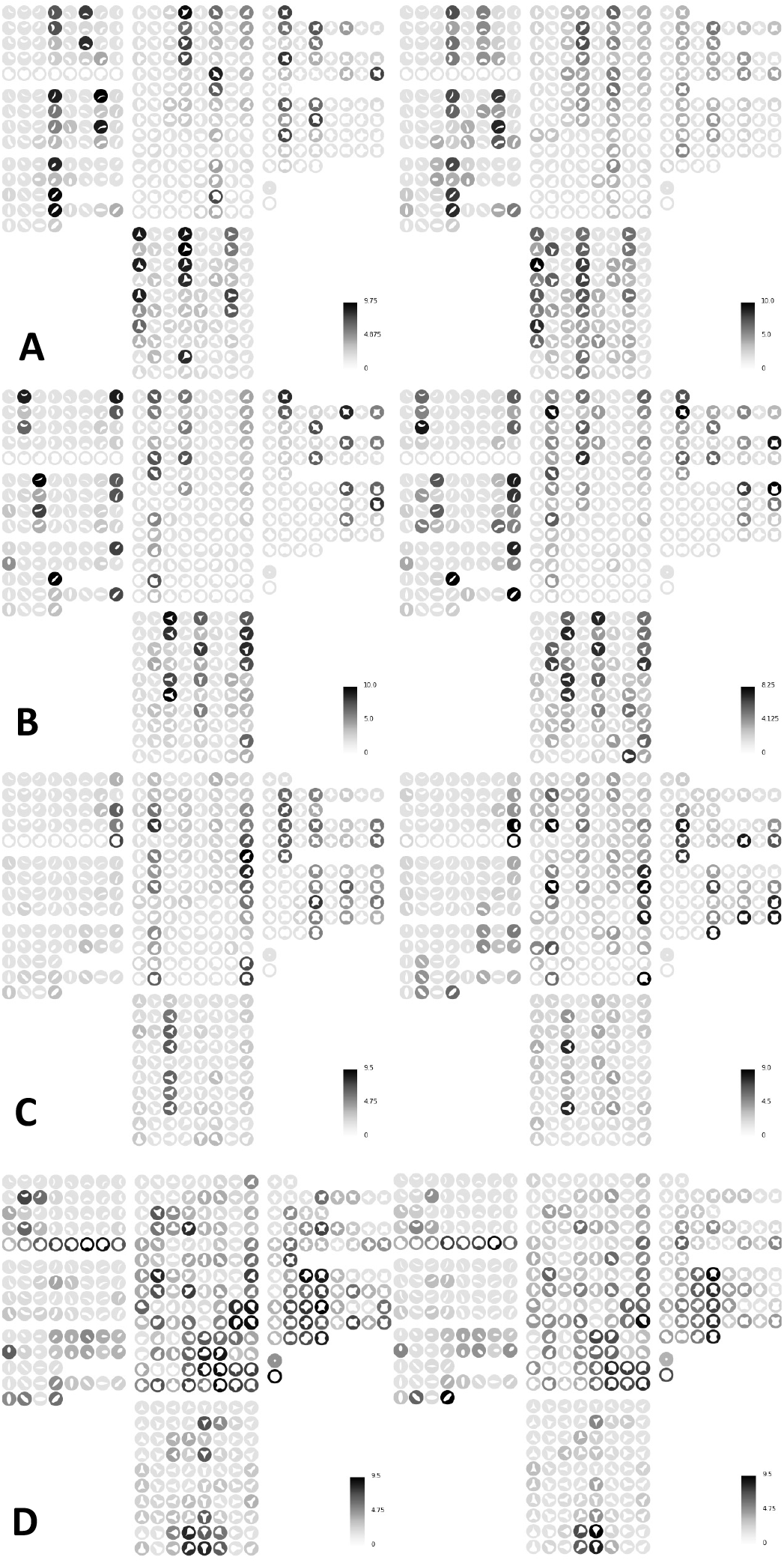
Simulated response to the stimuli of [Pasupathy and Connor, 2001] in silhouette (Left) vs. outline (Right). **A** The neuron of Figure 3A, acute convex curvature at the bottom left. **B** The neuron of Figure 3B, acute convex curvature at the top right. **C** The neuron of Figure 7B, concave curvature at the right. **D** The neuron of Figure 7B, broad convex curvature at the top.

**Fig. 18.**
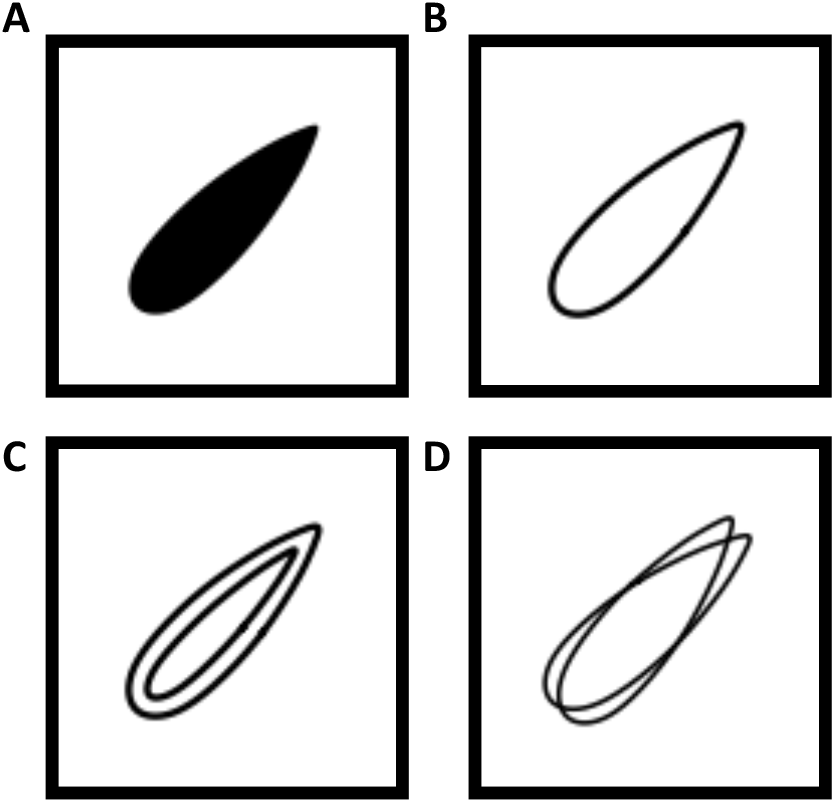
Examples of alternate stimuli based on [Pasupathy and Connor, 2001]. **A** One of the original stimuli. **B** Outline. **C** Contour pair. **D** Rotated pair.

A quantitative comparison of the response to the outline boundary stimuli as compared to the standard silhouette stimuli is shown in Table 2. The first measurements is the mean of the ratio of responses to corresponding shape outline stimuli and silhouette stimuli denoted simply |*Outline*| where the silhouette is assumed as the baseline. The second metric is the Pearson correlation coefficient between the responses, denoted *r*(*Outline*), treating the response as a vector over all the stimuli. These results confirm that the simulated concave selective neuron has a higher average response to outlines, while the simulated board convex selective neuron has a lower average response to the outline stimuli. The acute selective neurons have very nearly the same response to both outline and silhouette stimuli. At the same time, the concave selective neuron has the least correlation between the responses to outline and silhouette versions of the same stimuli.

**Table 2.** Comparison of Response to Silhouette and Outlines Boundary Stimuli. |*Outline* |: Average magnitude of responses for outline stimuli relative to the response of the standard silhouette stimuli. *r*(*Outline*) : Correlation of of outline and silhouette stimuli response vectors.

| Neuron | $\vert Outline\vert $ | $r(Outline)$ |
| --- | --- | --- |
| Acute Convex Left | 1.01 | 0.90 |
| Acute Convex Right | 1.05 | 0.87 |
| Concave Right | 1.21 | 0.66 |
| Broad Convex Top | 0.73 | 0.90 |
| Average | 1.00 | 0.83 |

The reasons for the different response to outlined versions of the medium curvature convex and concave boundary based stimuli are striking because both neurons are selective for curves of the same degree, the only difference being the convex versus concave orientation. The reason for the differences has to do with the slightly unequal response to V1 neurons selective for edges and lines. These differences will be detailed in the Method section on tuning the response properties of the model. This subtle variation between the responses of neurons selective for convex and concave medium curvature is an phenomena which could validate the model, or help to refine it.

#### 2.6.4 Response to Concentric and Rotated Outline Boundary Stimuli

Another set of experiments was performed to compare the response of the simulated V4 neurons when an outline of a boundary based stimuli is presented in two altered versions: A concentric version of the same shape as compared to a pair of outlines slightly rotated relative to each other. The concept is illustrated in Figure 18: The original stimuli is A, and an outline stimuli like the ones used in Section 2.6.3 is shown as B. Figure 18C is the concentric outline pair, and D is the rotated outline pair. In the rotated pair, two slightly rotated versions offset by +/-*π/*16 radians replace the original. As detailed in the Methods Section 4.6.1, some shapes were omitted from this data set because they were either too narrow to contain a concentric shape, or did not have meaningful rotations. The resulting set of stimuli consisted of 324 boundary configurations, out of the original 366. This experiment was chosen because it may help explain the results of [Srinath et al., 2021], which found that many V4 neurons are most responsive to 3-D objects. The concentric shape pairs (Figure 18C) were chosen because they resemble contour lines of shadows and luminescence that occur naturally on 3-D stimuli. The rotated shapes (18D) acts as a control that has approximately the same amount of total intensity in the receptive field and in almost the same configuration, but it does not have similarity to a 3-D object. See Section 4.6.1 for details on the generation of the stimuli used in these experiments.

The results in Table 3 show that overall the V4 neurons are more selective for the concentric outline pairs than the rotated outline pairs. This is particularly true for the neuron selective to medium curvature at the right, and also true to a lesser degree for the neurons selective to acute convex curvature. In contrast, the rotated pairs are less effective at stimulating these neurons; this is particularly true for the concave and medium curvature selective neurons but also to some degree for the neurons selective for acute convex curvature. The one exception is the neuron selective to concave curvature - for that neuron, the model suggests that concentric pair stimuli will have a slightly reduced response in comparison so the simple outline, while the rotated pair stimuli will have a moderately increased response. These results suggest that comparison of response for concentric shape pairs stimuli and rotated shape pairs would be a useful experiment for validating and improving the OC model and understanding selectivity of V4 neuron to 3-D objects described in [Srinath et al., 2021].

**Table 3.**
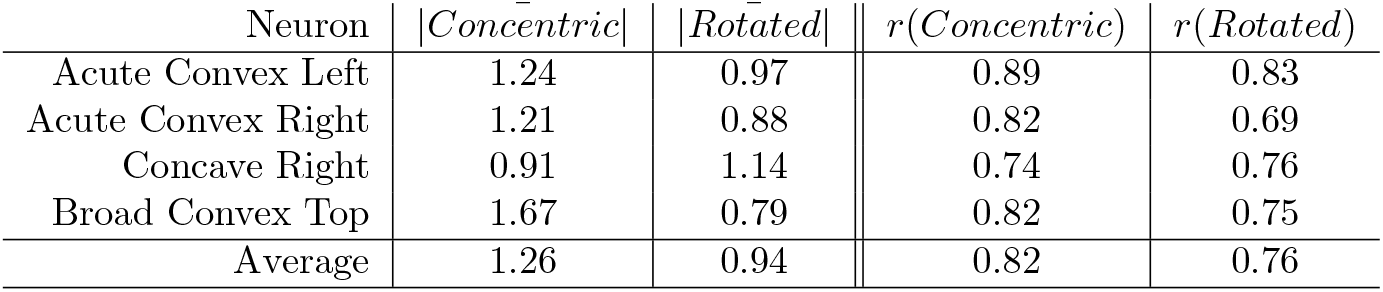
Response to Paired Concentric and Rotated Outline Boundary Stimuli: |…| : Average magnitude of response compared with the outline boundary stimuli. *r*(…) : Correlation of responses with response to outline boundary stimuli.

## 3 Discussion

### 3.1 Conclusions

The Organic Convolution (OC) model is a simplified but biophysically accurate model for the feed forward propagation of spikes in cortex-like layers. The biophysical parameters for the time constants governing the spiking model PSP equations (Equation 1 and Table 4) are within plausible ranges as are the time constants for the distributions modeling the arrival and transmission of spikes (Table 5). Therefore OC meets the speed requirement in visual recognition [Thorpe et al., 2001] and shows how processing from V1 to V4 can occur with a cascade of single spikes.

**Table 4.** PSP Time Constants (ms) and properties for Equation 1; Source: [Holmgren et al., 2003].

| Type | $\tau_{rise}$ | $\tau_{fall}$ | 10-90% rise time | Width at half-amplitude |
| --- | --- | --- | --- | --- |
| Excitatory Pyramidal / Stellate | 4.5 | 30 | 5.3 | 36 |
| Inhibitory Fast Spiking | 1.9 | 13 | 2.2 | 15 |

**Table 5.**
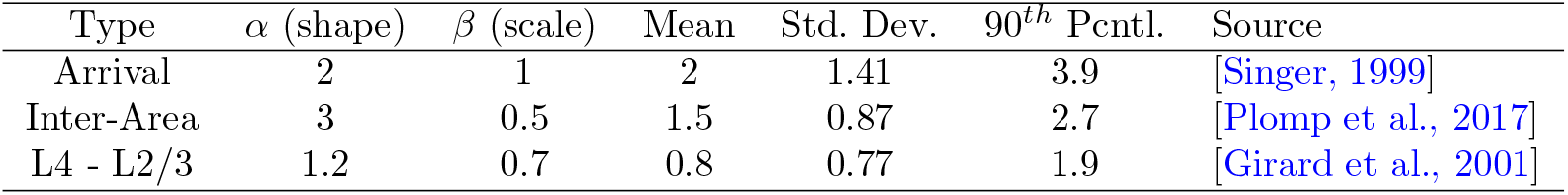
Arrival & Transmission Gamma Distribution Time Constants and properties (ms)

The debatable part of this model is whether axons and dendrites can assemble into the proposed patterns of afferent connections throughout cortex. The connections patterns seem biologically plausible in that they define local receptive fields and are quite simple. The evidence from V1 [Ko et al., 2013] and V2 [Kim et al., 2014] is that neurons do pre-wire into computationally useful patterns. The proposed functional connection patterns in this study are suggestions for the types of connections are sufficient to match V4 recording data but are not definitive.

It also is not realistic to assume that all neurons of a given type will be regularly shaped nor is it plausible that all connection patterns of similar units at different locations are exactly identical. The OC model in this study makes more sense when the connection patterns are viewed as a model for the “average” or “ideal” version of each unit type. Cortex may contains many units that follow a connection pattern in any area of the receptive field, but each individual unit must have its own irregular shape determined by its dendritic growth pattern around the other elements in the neuropil. When combined such units may functionally achieve a robust version of the average connection pattern. Investigating OC models with realistic numbers of units, more variety in unit shape and connections, and transmission failures is an attractive area for further research.

For understanding the nature of visual processing in V4, the OC model may suggest a synthesis of the points of view in different neurophysiological approaches. Studies suggesting V4 responds in a place invariant manner to shapes defined by closed paths [Pasupathy and Connor, 2001, Kim et al., 2019] may be reconciled with the fine shape and orientation tuning shown in [Nandy et al., 2013]: The same underlying curvature selective connection patterns are present in both types of units. Selectivity to a curvature at a specific position in the receptive field is a partial implementation of a rotational connection pattern that may also wrap around the entire receptive field.

The recent results in [Hu et al., 2020] seem in agreement with [Pasupathy and Connor, 2001] and [Nandy et al., 2013] and the OC model in suggesting that an important function of V4 may be to represent local variations in orientation and curvature. The best way to resolve questions of which stimuli best characterize V4 neurons would be to use multiple types of stimuli in a single study, and see if a single model can reproduce results collected under uniform conditions. To reconcile earlier studies with the recent findings of [Srinath et al., 2021] that V4 neurons may prefer 3-D shapes, these should be compared to the response to similar shaded, outlined, and contoured versions of the same shape. This study also suggested that a stimuli of corners like the one in [Nandy et al., 2013] may be optimal for probing V4 receptive fields, because stimuli may be a good description of the preferred stimuli of V2 afferent inputs.

At the same time, with the greatest respect for the painstaking work of neurophysiological recording in awake subjects, and in particular the difficulty of recording from visual area V4, an outside observer is left with the impression that neurophysiologists studying V4 resemble the parable of the blind men and an elephant: Each study shows a different kind of selectivity depending on the nature of the stimuli presented. This modeling study attempted to point to a common underlying mechanisms with which all of these behaviors can be explained.

This study does not include a model of an unsupervised connection learning algorithm (e.g. [Miller, 1994]) that can generate the connection patterns. This may seem like a weakness, but to defuse this criticism consider an analogy: Biologists often observe stripes of pigment on the bodies of animals or plants and reach conclusions about the function of those stripes for camouflage, signals, etc. But no one expects the biological mechanisms by which pigmented stripes are created to involve an unsupervised learning algorithm. In the same way, neuroscientists can reason about the functions of simple strips of a certain types of preferred connections. The function of such simple arrangements may be understood before we understand the mechanism of their generation. For example the connection arrangements for motion detection [Ko et al., 2013] are understood functionally but the precise mechanism of their generation is still a matter under investigation.

For practical reasons, AI studies of neural networks have focused on algorithms to learn the large number of neural network parameters from labeled data with Deep Learning algorithms. If the ideas behind the OC model are correct, then evolution has found developmental mechanisms to create simple patterns that perform some of the same function without learning from experience. The approach in this study is to use models with as few parameters as possible, so that a non-gradient based optimization and a small amount of deliberate manual tuning sufficed to find adequate parameterization to match the data. In contrast, a recent proposal is that “goal driven” computational neuroscience studies should use Deep Learning [Yamins et al., 2014, Kietzmann et al., 2019]. These studies can be seen as implying that models with designed parameters are retrograde for neuroscience, just as it is for artificial intelligence. But this study shows that biophysicists seeking mechanistic explanations for cognitive processes should favor parsimony in the underlying model over convenience in finding useful parameterizations. The model in this study uses orders of magnitude fewer parameters than DL networks and still reproduces neurophysiological results very well. Whether or not this particular embodiment of Organic Convolution is exactly right, this is evidence for the “critique of pure learning” of [Zador, 2019], that much of brain function must come from simple structures encoded in the genome.

This study demonstrates a novel method for simulating waves of individual spikes in neural networks with connection patterns that can be modeled by convolution using GPU hardware . While most of the issues raised in this discussion are not resolved definitively, it is clear that convolutional neural networks and tensor technologies can be used to experiment with models of afferent connection patterns in spiking neural networks at scale. Such models may be quite simple and still help explain neurophysiological data.

### 3.2 Ideas and Speculation

Proof of the Organic Convolution model could come from direct observation using connectomics techniques, for review see e.g. [Behrens and Sporns, 2012]. With the connectomics technologies that have become available in the past decade, the details of the V1, V2 and V4 feedforward connections could be mapped at different stages of development to provide direct evidence for the number and kind of connections that exist. The results in this study suggest it may be especially interesting to analyze the geometric arrangements of connections in topographically organized cortical afferents. Also, additional confirmation of Organic Convolution would require understanding of the mechanisms that guide such pattern formation, as discussed in [Shirasaki and Pfaff, 2002]).

The coding used in the convolution model is a population place code: each unit responds to the presence of a particular pattern of inputs at a particular location in the visual field, albeit with broad tuning, and there are functionally identical units tiling the receptive field. Given the place code in the OC model, the functioning of the model is essentially the same whether considered as a binary network or a spiking network, due to the long time scale of integration and tight synchronization of the input. If the OC model is an accurate representation of brain activity, it suggests that the venerable McCullough-Pitts neuron model [McCulloch and Pitts, 1943] may go a long way in explaining important classes of brain activity. Also note that the OC model is compatible with earliest arrival spike time coding proposed in [Thorpe et al., 2001]: If the inputs to area V1 in the OC model are ordered so that the earliest arrivals are from the most intense stimuli, it may enhance the place code defined by the connection patterns. Models combining OC with spike time coding is an attractive future area of research.

The experiments in this study model a single cascade of spikes as the basic unit of computation, and such a system will manifest rate coding as well as place coding: If there are repeated waves of activity in response to a single stimuli, the place coded neurons best matching a stimuli would fire with the fewest failures and at a higher overall rate. But there is an important distinction between the meaning of “coding” in these two usages: The topographic place code is a mechanistic element of the computation because the patterns of connections rely on locality - without the place coding, the computation would not work. The term rate coding used here means that information can be read out from the network by the rate. But the rate is not a part of the computation: the biomechanics of the computation relies on the synchronized cascade. If each neuron spikes at most once, there is no rate code at all in the elementary computation. This raises an interesting new challenge to the biological plausibility of a common DCNN methodology: the RelU activation function. The Relu function is justified as biologically plausible because it is taken to represent a neural unit which has rates as inputs and a rate as the output [Glorot et al., 2011] - the RelU hinge is taken as increasing output rate once the stimuli match to the convolution filters is above the threshold. But if it is true that the biological mechanism of neuronal computation does not take rates as inputs and produces a rate as an output, a RelU activation neural network is not a biologically plausible neural network.

An important related question is precisely how Organic Convolution is used to achieve object recognition in IT. The convolution filters used in the OC model in this study are not exactly like those discovered in trained machine learning models, but they may still serve a similar purpose. Initial experiments suggest a network like the OC model may be used to perform machine learning tasks by feeding the final convolution output into a fully connected network that is trained with gradient descent [LeCun et al., 2015]. If OC is found to be a good representation of biology, and it can also function in object recognition applications, it may serve as a bridge for understanding the relationship between the current generation of artificial neural networks and the brain.

## 4 Materials and Methods

### 4.1 Spiking Neuron Model

The spiking neuron model used for the Spike Time Convolution (STC) model is the Spike Response Model (SRM) described in [Gerstner and Kistler, 2002]. The SRM model is an abstraction of the integrate and fire neuronal biophysics but it captures the key properties of spike integration and timing for single action potentials. Each model neuron has a potential level given by a the sum of the excitatory and inhibitory post synaptic potentials (PSP). The time course of the PSP’s are described by a difference of decaying exponentials. For any given synapse (indexed by *i*) and given an input spike arrival time *s* the influence of the PSP is given by the kernel:

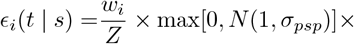

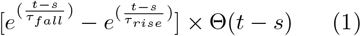

In equation 1 *w*_*i*_ represents the synaptic input sign and weight. The term max[0, *N* (1, *σ*_*psp*_)] provides a random scaling to every PSP. For the experiments presented in this study *σ*_*psp*_ = 0.25. With this choice, 68% of PSP’s will be within 0.75-1.25 of their default weight, 95% will be between 0.5 and 1.5, and the max operation ensures that tail events cannot reverse the sign of a PSP. Weights are positive for excitatory and negative for inhibitory, depending on the type of the afferent input and the weight is determined by the connection pattern to topographically organized afferent inputs, as described below; *Z* is a normalization constant chosen so that the maximum value of the PSP without random noise scaling is *w*_*i*_; Θ(*x*) denotes the Heaviside step function. The simulations of each wave of cortical activity lasted 30 ms, with a discretization step of 0.3 ms in the simulated PSP dynamics.

Given the PSP kernel function in equation 1 and a set of input spike times on a set *S* of afferent inputs indexed by *i* the neuron spikes at the time

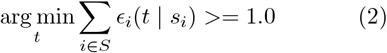

where the the threshold in equation 2 is set to 1.0 for model interpretability: with this definition, all synaptic weights *w* in the convolution kernels (Figures 20, 21, 12, 10 and 6) are expressed in terms of the positive excitatory threshold. The constant *Z* in equation 1 is determined by first calculating the time that PSP achieves its maximum, as described in the Appendix.

**Fig. 19.**
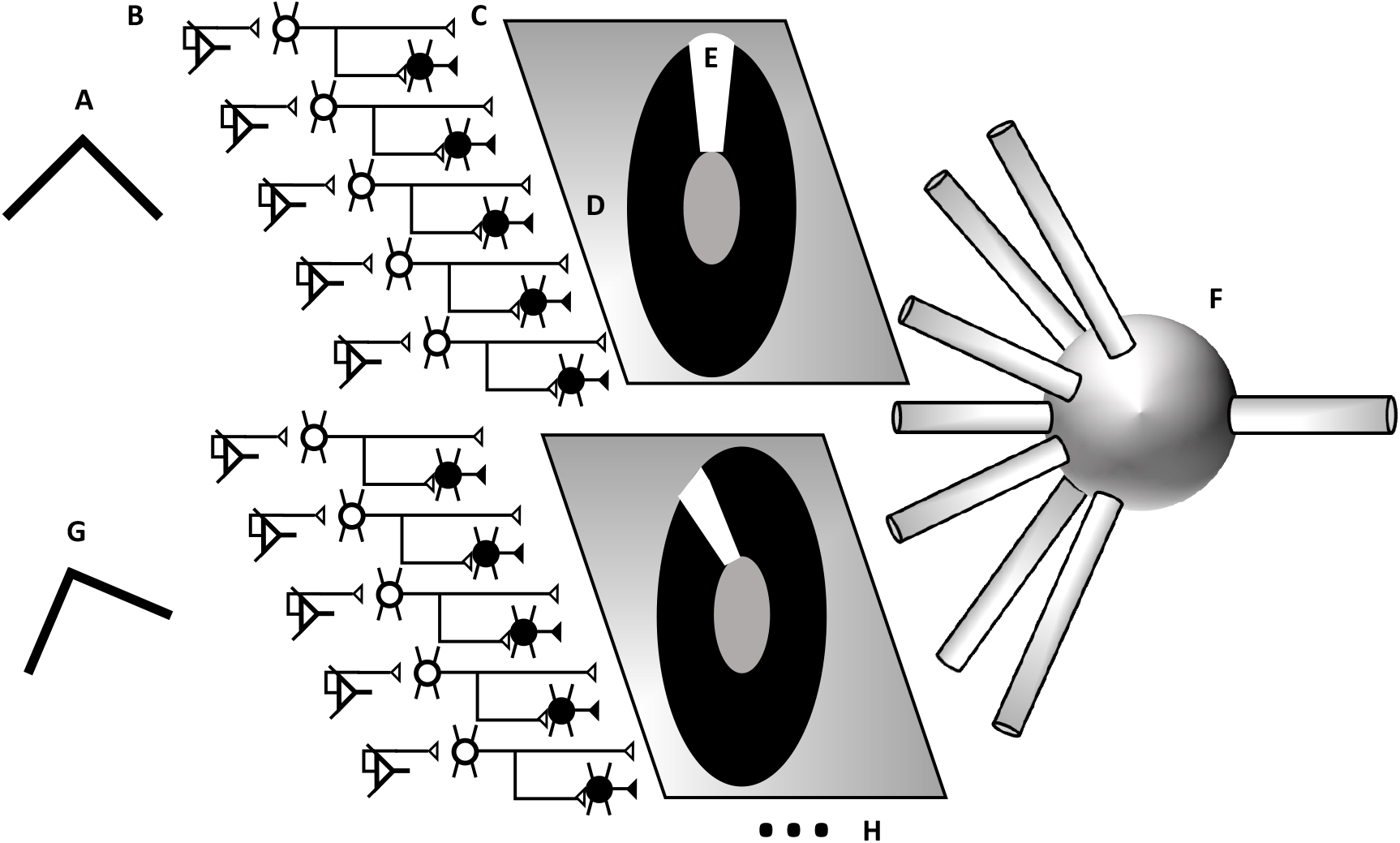
Afferent Connection pattern illustration for a V4 curvature selective neuron. **A** V2 contains neurons selective to corners with a variety of interior angles and orientations in visual space, e.g. 90 ° corners oriented up. **B** Each type of selectivity is repeated in units covering the visual space in a topographically organized manner. Such relations are maintained as the signal is passed to V4 L4 neurons, and V4 fast spiking interneurons. **C** The topographically organized excitatory and inhibitory afferents are available to V4 L2/3 neurons. Afferents are selected to form patterns of inhibition D and excitation E by the basal dendrites and proximal apical dendrites which extend radially in the topographic plain from the V4 L2/3 pyramidal neuron soma. G The same systematic arrangements repeat for the other interior angles and rotations of selectivity available from V2. The same framework also applies to afferent inputs from the LGN in V1 and a_erents from V1 in V2.

**Fig. 20.**
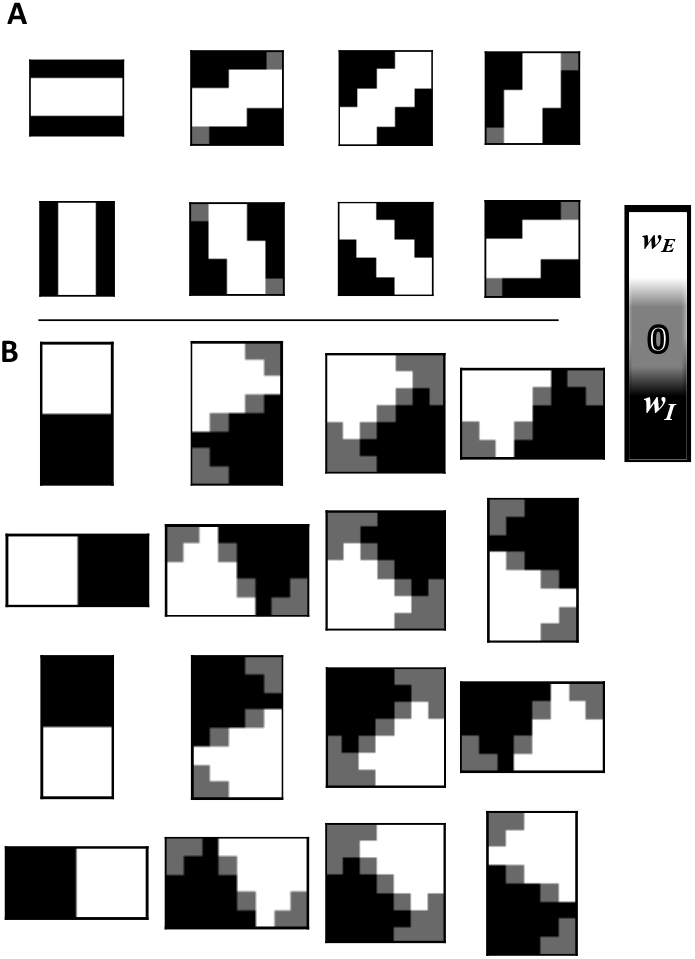
Afferent Connection Patterns for V1 model neurons. The convolution pattern represents the afferent connections to excitatory and inhibitory inputs as illustrated in Figure 19. **A** The synaptic connection strength versus position in topographic space relative to the soma at the center for V1 neurons selective to lines/bands of intensity. There are 8 different orientations, oriented 22.5^°^ apart, resulting in 3 unique filter variants. The filters are 4×5 (horizontal/vertical) pixels or 5×5 pixels (others). **B** The synaptic connection strength versus position in topographic space relative to the soma at the center for V1 neurons selective to edges in the intensity. There are 16 different orientations oriented 22.5^°^ apart resulting in 3 unique filter variants. The filters are 4x8 (horizontal/verical), 7×7 pixels (45^°^ rotations) and 5x8 (others). The excitatory and inhibitory weight values for the filters are shown in Table 8

**Fig. 21.**
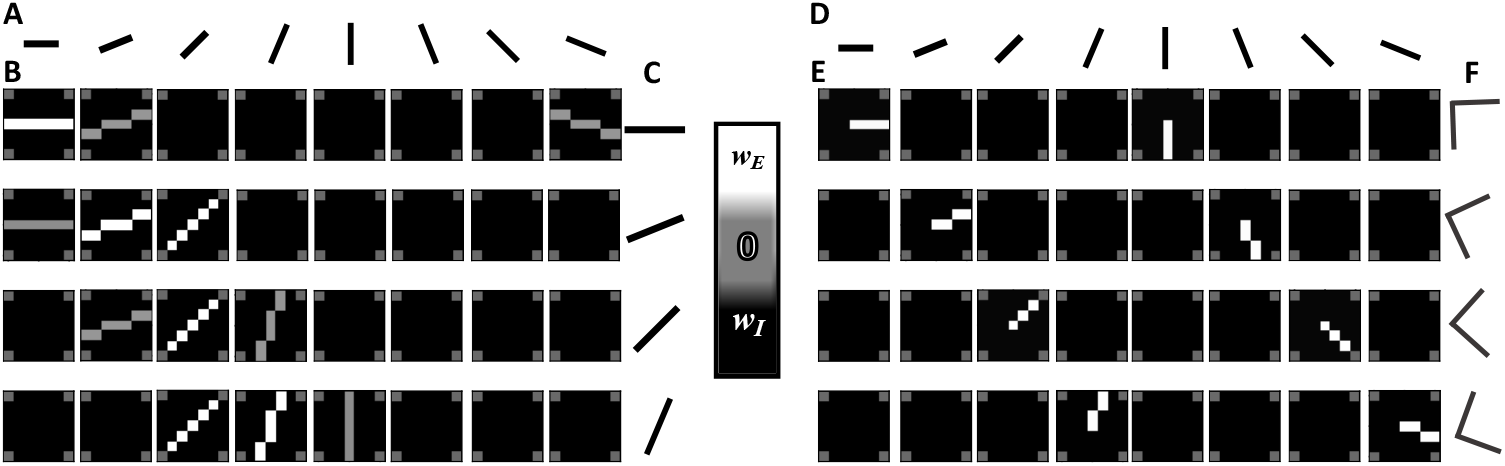
Afferent Connection Patterns for model V2 L2/3 units. The convolution pattern represents the afferent connections to excitatory and inhibitory inputs as illustrated in Figure 19. **A**-**C** Illustration of a connection pattern for an ultra-long type V2 unit. **D**-**F** Illustration of a connection pattern for a complex type V2 unit having an interior angle of 90^°^. **A & D** Pictorial representation of the preferred stimuli orientation of V1 afferents inputs to V2. **B** The synaptic connection strength versus position in topographic space relative to the soma at the center for four V2 neurons selective to long, straight edges or lines; each row are the 7x7 convolution filters for one type of V2 unit. Four orientations are illustrated, ranging from horizontal to an upward angle of 67.5^°^ in steps of 22.5^°^, and the model has a total of 8 orientations. The strongest excitation are for inputs aligned to preferred orientation of the unit in a line down the center of the receptive field. There are weaker excitatory inputs having orientations 22.5^°^ from the preferred orientation. **C** Pictorial representation of the preferred selectivity for the unit having the convolution filters in the same row. **E** The synaptic connection strength versus position in topographic space relative to the soma at the center for four V2 neurons selective to corners having a 90^°^ interior angle, and four different orientations 22.5^°^ apart. The model contains similar units having interior angles ranging from 45^°^ to 157.5^°^ in steps of 22.5^°^. Each such unit has 16 different preferred orientations, for a total of 96 different types of units. The pattern is formed by having two excitatory strips running from the center to the edge of the receptive field, each coming from an afferent with an appropriate orientation for one arm of a corner. The positions of the excitatory strip rotate with the preferred orientation of the unit. **F** Pictorial representation of the preferred selectivity for the unit having the convolution filters in the same row.

Time constants for the PSP’s in the model are based on [Holmgren et al., 2003] and shown in Table 4. [Holmgren et al., 2003] showed that the time constant of both EPSP and IPSP in pyramidal neurons are approximately the same in rise time and differ primarily in decay time (these findings had large error bars around the mean). The decay times are so long that they do not really affect the spike time in a feedforward cascade like those modeled in this study, so for simplicity this model used the same time constants shown labeled “Excitatory” for EPSP and IPSP in all of the excitatory neurons in the model (“Excitatory” in 4 refers to the neuron, not the synaptic input.) [Holmgren et al., 2003] found that the Excitatory connections onto FS spiking interneurons had a much faster rise time and decay and they are approximated by the parameters labeled “Inhibitory” in table 4 (“Inhibitory” refers to the excitatory synapse in the FS inhibitory cells, not the inhibitory synapse on the pyramidal cell.)

**Table 6.**
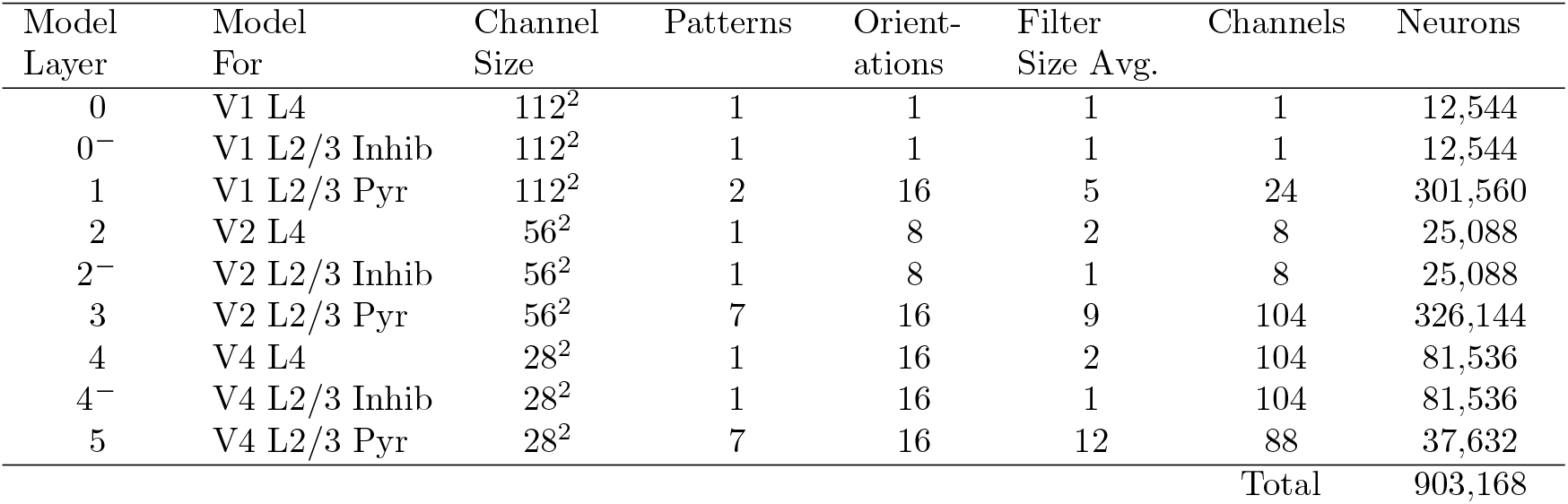
Network Summary.

**Table 7.** Connection Pattern Summary.

| Type | Orientations<br>(Channels) | Variants | Weights | Field | Radii | Non-Wt<br>Params | Total<br>Params |
| --- | --- | --- | --- | --- | --- | --- | --- |
| V1 L4 Input | 1 | 1 | NA | $1 \times 1$ | NA | 0 | 0 |
| V1 L2/3 Inhibit | 1 | 1 | 1 | $1 \times 1$ | NA | 0 | 1 |
| V1 Strip | 8 | 3 | 6 | $4 \times 5$ | 24 | 3 | 9 |
| V1 Edge | 16 | 3 | 6 | $4 \times 8$ | 32 | 2 | 8 |
| V2 L4 Pool | 8 | 1 | 1 | $2 \times 2$ | 4 | 0 | 1 |
| V2 L2/3 Inhibit | 8 | 1 | 1 | $1 \times 1$ | NA | 2 | 3 |
| V2 L2/3 $45^\circ$ | 16 | 2 | 5 | $9 \times 9$ | 32 | 2 | 7 |
| V2 L2/3 $67.5^\circ$ | 16 | 2 | 6 | $9 \times 9$ | 32 | 2 | 8 |
| V2 L2/3 $90^\circ$ | 16 | 3 | 6 | $9 \times 9$ | 32 | 2 | 8 |
| V2 L2/3 $112.5^\circ$ | 16 | 2 | 6 | $9 \times 9$ | 32 | 2 | 8 |
| V2 L2/3 $135^\circ$ | 16 | 2 | 5 | $9 \times 9$ | 32 | 2 | 7 |
| V2 L2/3 $157.5^\circ$ | 16 | 2 | 6 | $9 \times 9$ | 32 | 2 | 8 |
| V2 L2/3 $180^\circ$ | 8 | 3 | 10 | $7 \times 7$ | 32 | 2 | 12 |
| V4 L4 Pool | 16 | 1 | 1 | $2 \times 2$ | 4 | 0 | 1 |
| V4 L2/3 Inhibit | 8 | 1 | 1 | $1 \times 1$ | NA | 0 | 1 |
| V4 Acute Convex | 16 | 1 | 2 | $24 \times 24$ | 128 | 7 | 9 |
| V4 Concave | 16 | 1 | 2 | $24 \times 24$ | 128 | 8 | 10 |
| V4 Medium Convex | 16 | 1 | 2 | $24 \times 24$ | 128 | 8 | 10 |
| V4 Low Curve | 8 | 1 | 2 | $15 \times 20$ | 96 | 8 | 10 |
| V4 Medium Curve | 8 | 1 | 2 | $16 \times 22$ | 96 | 8 | 10 |
| V4 High Curve | 8 | 1 | 2 | $16 \times 32$ | 96 | 8 | 10 |
| Total (21) | 242 | 34 | 73 |  |  |  | 141 |

**Table 8.** V1 Pattern Weights by Pattern and Rotational Variant. *w*_*E*_ : Excitatory input strength per afferent input. *w*_*I*_ : Inhibitory strength per afferent input. ∑*w*_*E*_ : Total excitatory weight. ∑*w*_*I*_ : Total inhibitory weight. Units: 1=Threshold.

| Pattern | Variant | $w_E$ | $w_I$ | $\sum w_E$ | $\sum w_I$ |
| --- | --- | --- | --- | --- | --- |
| Stripe | 0° | 0.19 | -0.29 | 1.9 | -2.9 |
| Stripe | 22.5° | 0.17 | -0.24 | 1.9 | -2.8 |
| Stripe | 45° | 0.14 | -0.24 | 1.8 | -2.8 |
| Edge | 0° | 0.07 | -0.11 | 1.1 | -1.7 |
| Edge | 22.5° | 0.08 | -0.12 | 1.2 | -1.8 |
| Edge | 45° | 0.07 | -0.1 | 1.1 | -1.7 |

Variability in the amplitude of individual cortical pyramidal cells is high: [Seeman et al., 2018] found the coefficient of variation (CV) to be approximately 0.6 in individual cells. However the units in this study represent idealized “average” units in which one model unit represents an average of a number of real neurons in mammalian visual cortex. For that reason the OC model uses a more conservative value of 0.25 CV on all PSP, which is embodied by multiplying the PSP value by a random value drawn from *N* (1, 0.25). See Section 4.3 for details of the PSP implementation as a tensor calculation.

#### 4.1.1 Input Model and Spike Transmission

The first layer of the network represents V1 L4 receiving input from LGN but these neurons are not modeled explicitly: V1 L4 units spike with a probability equal to the image intensity. The full PSP model of equation 2 begins with the models for V1 inhibitory and L2/3 pyramidal neurons. Because the stimuli (Figure 2) are black and white, in practice this means most V1 L4 neurons in the model spike deterministically with respect to a single image - only units representing an anti-aliased boundary in the stimuli have mid range intensity that lead to significant trial to trial variability in the response to the same image.

Although the occurrence of spikes in V1 L4 of the model is largely deterministic to the study stimuli, the timing of the V1 L4 spikes are random in every trial. The V1 L4 unit spike times are drawn from a gamma distribution with parameters in table 5. The arrival times are based on findings from [Singer, 1999] showing that inputs from LGN to V1 L4 arrive with a synchronization window of around 5ms. Inhibition in the V1 model (only) follows a simplified model in which the arrival time for the IPSP in the L2/3 pyramidal cell is simply delayed by a constant 1 ms [Hoffmann et al., 2015].

The delays for transmission time between layers in the same cortical area and between cortical areas are also modeled by gamma distributions with time constants in table 5. The L4 to L2/3 timing is based on [Girard et al., 2001] which show a mean of less than 1 ms transmission delay on average. The L4 - L2/3 delay distribution is used for V4 to Inhibitory, V4 to L2/3 Excitatory and V4 L2/3 Inhibitory to Excitatory delays. The transmission time between areas of cortex was studied in [Plomp et al., 2017] which found around 1.5 ms transmission delay. A distribution based on [Plomp et al., 2017] is used for transmission between L2/3 excitatory neurons and L4 neurons in the next cortical area model.

### 4.2 Network Model

The model network consisted of nine layers, representing L4 and L2/3 in cortical areas V1, V2, and V4; inhibitory and excitatory L2/3 neurons are considered separate layers. As shown in Table 6, each layer represents a number of model neurons that is the product of the layer area and the number of different types of units at each position. The channel size starts as the dimension of the input image in pixels (112 × 112) in the input layer, and at deeper layers the dimension shrinks due to pooling operations. The number of units at each position in the model is given by the number of connection patterns and the number of orientations at which the pattern is instantiated. Each unit type is modeled as a channel in the tensor simulation as described below. The total number of equivalent neurons represented by the model is approximately 900,000.

The numbered model layers in Table 6 refer to the convolutional or pooling network layers in the DCNN. In the model pooling layers are implemented with convolution operations as described in section 4.4.5. The input layer is listed as 0, and performs the function of transferring the input to real valued tensors as described in Section 4.1.1. Inhibitory layers are not listed as independent layers, because they do not perform either convolution or pooling but are tensor post-processing calculations attached to the preceding L4 pooling layer as described in Section 4.3. Therefore the V4 L2/3 pyramidal neurons are the 5^*th*^ convolutional layer in the model as shown in Table 6.

As explained in the Introduction, the model focuses on a simple feedforward wave of spikes and does not include other details of the cortical circuitry, such as lateral connections within cortical layers [Kisvárday et al., 1997] or cortical micro- circuits to L5/6 [Douglas and Martin, 2004]. The model also does not include inter-cortical circuits giving feedback from V4 to V2 and V1 and from V2 to V1, nor the interaction between feedback and lateral circuits [Liang et al., 2017].

Conceptually, the model assumes that the feedforward dynamics in the ventral pathway is driven by episodic arrival of inputs in V1 from the lateral geniculate nucleus (LGN), synchronized with precision on the order of a few milliseconds, which has been observed in multiple studies, as reviewed in [Singer, 1999]. The model as implemented is memoryless in that it only models single episodes of activity.

Table 6 also shows the average convolution filter size, which is the units receptive field in relation to its channel size. The average is taken over all types of units in the pyramidal layers. It shows that the filter sizes increase in the deeper layers. This increase is even steeper taking into account the pooling effect: The average filter diameter in the V2 L2/3 Pyramidal model is 8.7 in its own channel, which is equivalent to 17.4 in the input layer due to the 2× pooling. For the V4 L2/3 Pyramidal model the average filter diameter is equivalent to 50 pixels in the input layer.

The increase in receptive field size in the model is roughly in line with observations from monkey visual cortex: In V1, receptive fields are on the order of 1^°^ [Price and Born, 2009, Van den Bergh et al., 2010]. Studies have shown that V2 units have 2-4 times larger receptive field than V1 [Price and Born, 2009]. In the model, the ratio of V2 to V1 filter sizes is 3.3× after translating the V2 receptive field into V1 layer units. For complete details on the convolution filters in the V1 and V2 models see the Methods Sections 4.4.2 and 4.4.3 respectively.

Past measurement of V4 receptive field sizes range from 2-10^°^ [Roe et al., 2012], or 2-10× the average V1 receptive field size. The model is at the high end of the plausible range: The model ratio of V4 to V1 receptive field is 9.4× after correcting for pooling, and assuming the average model pixel filter size 5.3 in V1 represents 1^°^ of visual field. This high ratio found in this model study is likely due to selection bias in the neurophysiological used for tuning the model units: The results from [Nandy et al., 2013] focus on neurons with particularly large and interesting receptive fields. Overall the ratio of receptive field sizes in the model are plausibly in line with past neurophysiological studies. For further details on the sizes of each type of channel in the pyramidal layers see the Supplemental Methods Section 4.4.4.

### 4.3 Tensor Simulations

#### 4.3.1 Convolution With a Binary Neuron Model

As an introduction to the Spike Time Convolution (STC) model, listing 1 is the Python code for a convolutional neural network layer in which the model neuron is a simple binary threshold unit with no time dependence [McCulloch and Pitts, 1943]. Listing 1 uses the `tensorflow` and `keras` packages: The class `BinaryLayer` descends from `tf.keras.layers.Layer` . `BinaryLayer` has convolution filter weights (and a stride) defined at initialization. The `call` function in Listing 1 defines the transformation from input tensors to output tensors for the layer. The convolution itself is performed with the `tensorflow` model function `nn.conv2d` . To perform a simulation, multiple `BinaryLayer` ’s are added to a `tf.keras.Sequential` model object, as shown on lines 20-26 of Listing 1. After adding the layers, the model is compiled and outputs can be generated from a `tf.data.Dataset` holding the stimuli.

The code shown in Listings 1 is a simplification of that used for the simulations shown in the Results section: For one thing, it assumes the convolution filters Listing 1 perform the convolution using a single convolution filter size (lines 11-14). It is conventional to use a single filter size in DCNN applications for machine learning. For the Organic Convolution experiments described here, the filter sizes are determined by the receptive fields observed in recordings and they vary depending on the unit, particularly in the V1 and V4 models.

**Listing 1** Binary Convol

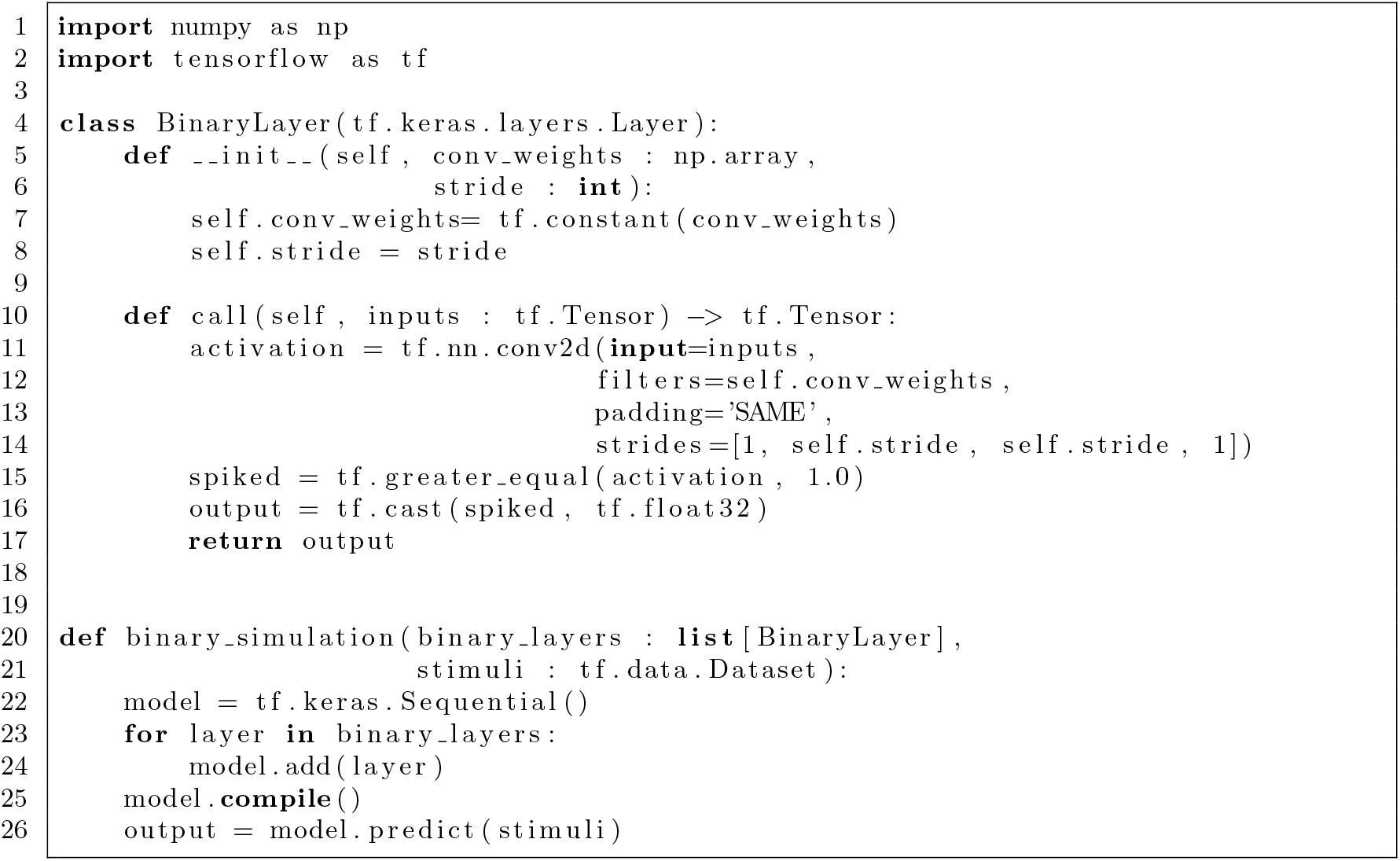

Note that the `BinaryLayer` shown in listing 1 is not just an illustrative abstraction: During research and development, connection patterns were first tested with a network of binary neurons just like in listing 1. The behavior of the binary model is a non-random version of the behavior of the spiking model, and the speed of the binary simulations makes it useful for prototyping. After prototyping with a binary model, the weights were increased and retuned so the units would spike appropriately under the noisy conditions of the spiking simulation.

#### 4.3.2 Spike Time Convolution With the Spike Response Model

Python code to model one layer of a Spike Time Convolutional neural network implementing the Spike Response Model is shown in Listing 2 and Listing 3, and 4. Listing 2 shows the `build` function, which is the standard tensorflow function to setup a customer layer. Listing 3 shows the `call` function, which transforms input to outputs. Listing 4 is a helper function to do the inhibitory interneuron calculations.

The layer class `SpikingLayer` inherits from the `BinaryLayer` in Listing 1 and shares the same code for the convolution operation. The `SpikingLayer` has the biophysical parameters as arguments:

A boolean `is l23pyr` indicates whether the layer is an L2/3 pyramidal convolution layer, or if not then an L4 pooling layer.

- The PSP time constants (*τ_rise* and *τ_fall* in Equation 1) are the required arguments `epsp_trise` and `epsp_tfall` for excitable, and optional arguments `ipsp_trise` and `ipsp_tfall` for inhibitory.
- The standard deviation of PSP variability is `psp_stdev` .
- The gamma distribution of delays from the *prior* layer to the current layer is `delay_alpha` and `delay_beta`, which is set to the appropriate values from Table 5 upon construction.
- A connection strength for the inhibitory interneuron’s interior PSP which does not use convolution, `ipsp_wt`

**Listing 2** Spike Time Convolution Layer Setup

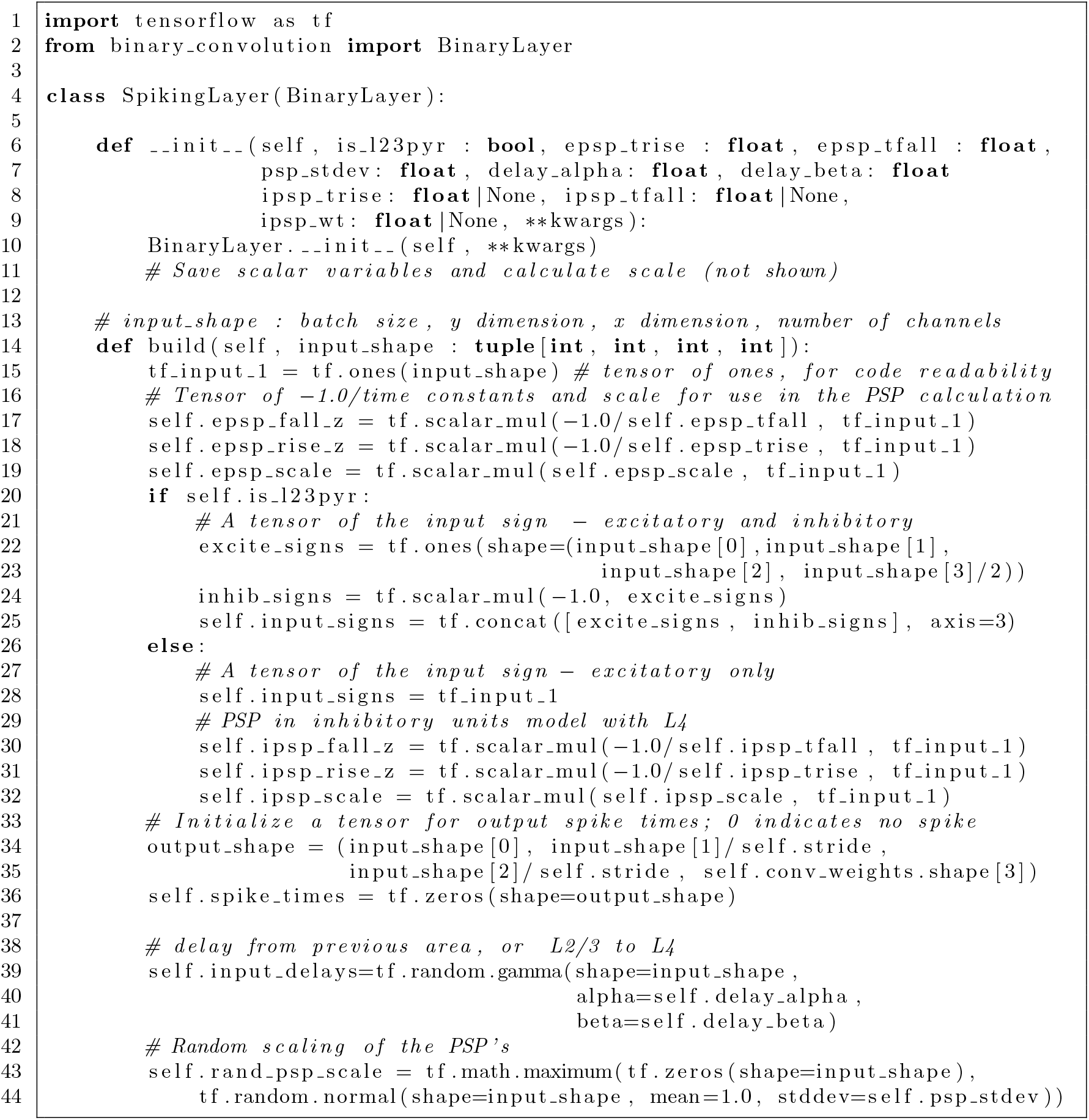

The full details of the `SpikingLayer` initialization are omitted from Listing 2 for brevity. The constructor simply copyies the parameters to member variables - the more important setup is performed in the `build` function.

Neural network layers in the `keras` framework have the option to define a `build` function that performs additional setup after initialization, given the input shape. (The input shape is unknown at initialization.)

**Listing 3** Spike Time Convolution Call Function

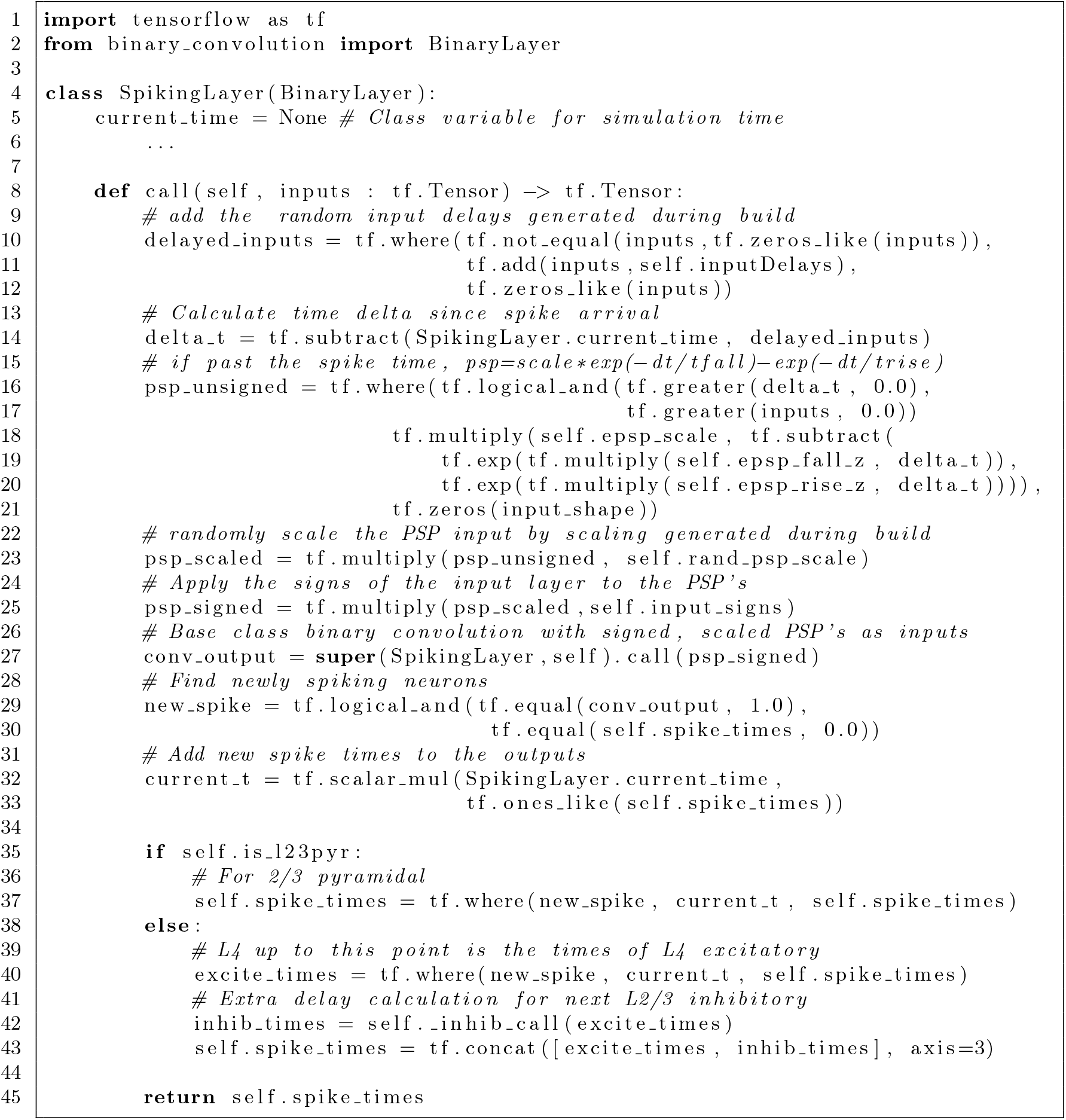

The `SpikingLayer` defines a `build` function, shown in Listing 2, to perform the following setup tasks:

- Define tensor versions of the PSP time constants having the same shape as the input (lines 17- 18, for excitatory neuron PSPs and 30-32 for inhibitory neuron PSP’s)
- For layers with both excitatory and inhibitory input, create a tensor of signs to apply to the PSP summation so that the inhibitory units reduce activation (lines 19-26).
- Create a tensor variable to hold the spike times for every unit in the layer (lines 34-36).
- Define the tensor of random delay times that will be applied to input spike times (lines 39-41)
- Define the tensor of PSP random size variation that will be applied to input spikes (lines 43-44.)

**Listing 4** Spike Time Convolution Inhibition Delay Helper Function

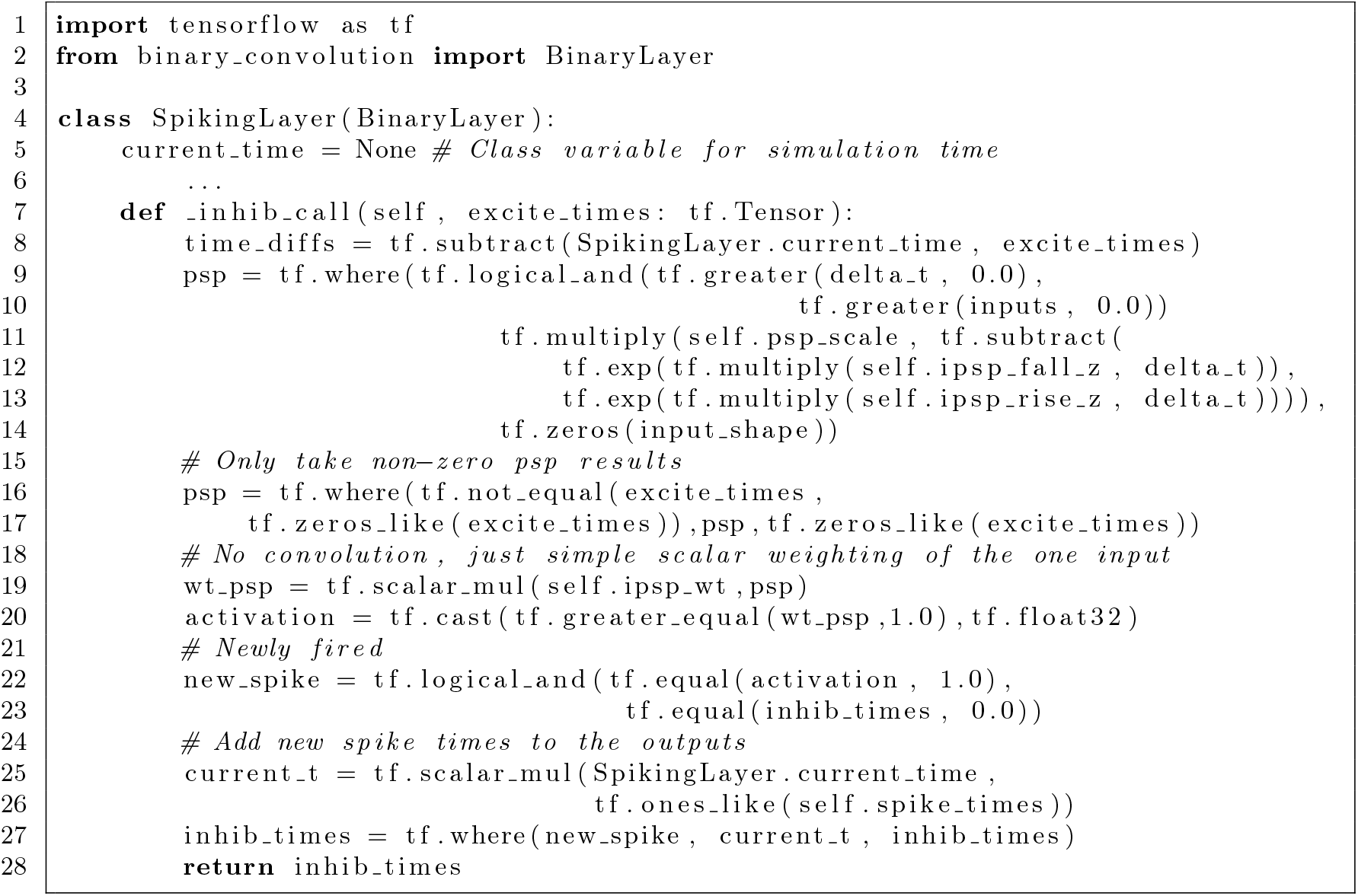

Note that as described in Section 4.2, L2/3 inhibitory interneurons are modeled with a post-processing calculation as part of the preceding L4 layer. For this reason, a non-L2/3 pyramidal network layer (variable `self.is_l23pyr`) includes both time constants for excitatory and inhibitory neuron PSP’s.

The `call` function of the `SpikingLayer` takes an input tensor that represents the spike times of the units in the previous layer. The spike time tensor stores zero to indicate no spike. Note that the `SpikingLayer` also has a *class* variable `current_time` that is used to hold a reference to the current simulation time (Listing 3 line 5).As will be explained below, the simulation starts with one discretization step after 0, and the class variable `current_time`, is set before the `call` method is invoked (Listing 5).

In `call` for the `SpikingLayer`, shown in Listing 3, the main steps are as follows:

1. Add the delay from the previous layers spike times to any inputs that spike (line 10)
2. Calculate the time since the spike of each unit that has fired (line 14); negative values result for inputs that have not spiked, but these are screened out at the next step.
3. Calculate the current PSP activations for excitatory neurons (L4 or L2/3 pyramidal) with the SRM model (line 16). This includes logic equivalent to the step function in equation 1.
4. Apply the previously defined random scaling constants to the PSPs (line 23.)
5. For those inputs that are inhibitory, apply the required sign to the PSP activation (line 25)
6. Invoke the base class `call` function which performs the convolution with the current PSP values as the inputs (line 27). This step converts the input dimension to the output dimension, and emits a 1 for every neuron that has fired.
7. Find what inputs have newly fired by comparing the current convolution outputs to the previous spike times and store it in another tensor (line 29-32).
8. For L2/3 pyramidal neuron layers, this is the end - the current time is stored in the member variable `spike_times` for all of the units that have newly spiked (lines 3
9. For L4 + L2/3 interneurons layers, the spikes at this stage represent the L4 neuron excitatory output spikes. The timing of resulting inhibitory interneuron spikes in L2/3 are calculated in a additional helper function `inhib call` on line 42, and combined with the excitatory spike_times into a combined output vector for the next L2/3 pyramidal layer (line 43).
10. Return the newly updated member variable of spike_times as the new output vector (line 45)

**Listing 5** Spike Time Convolution Simulation

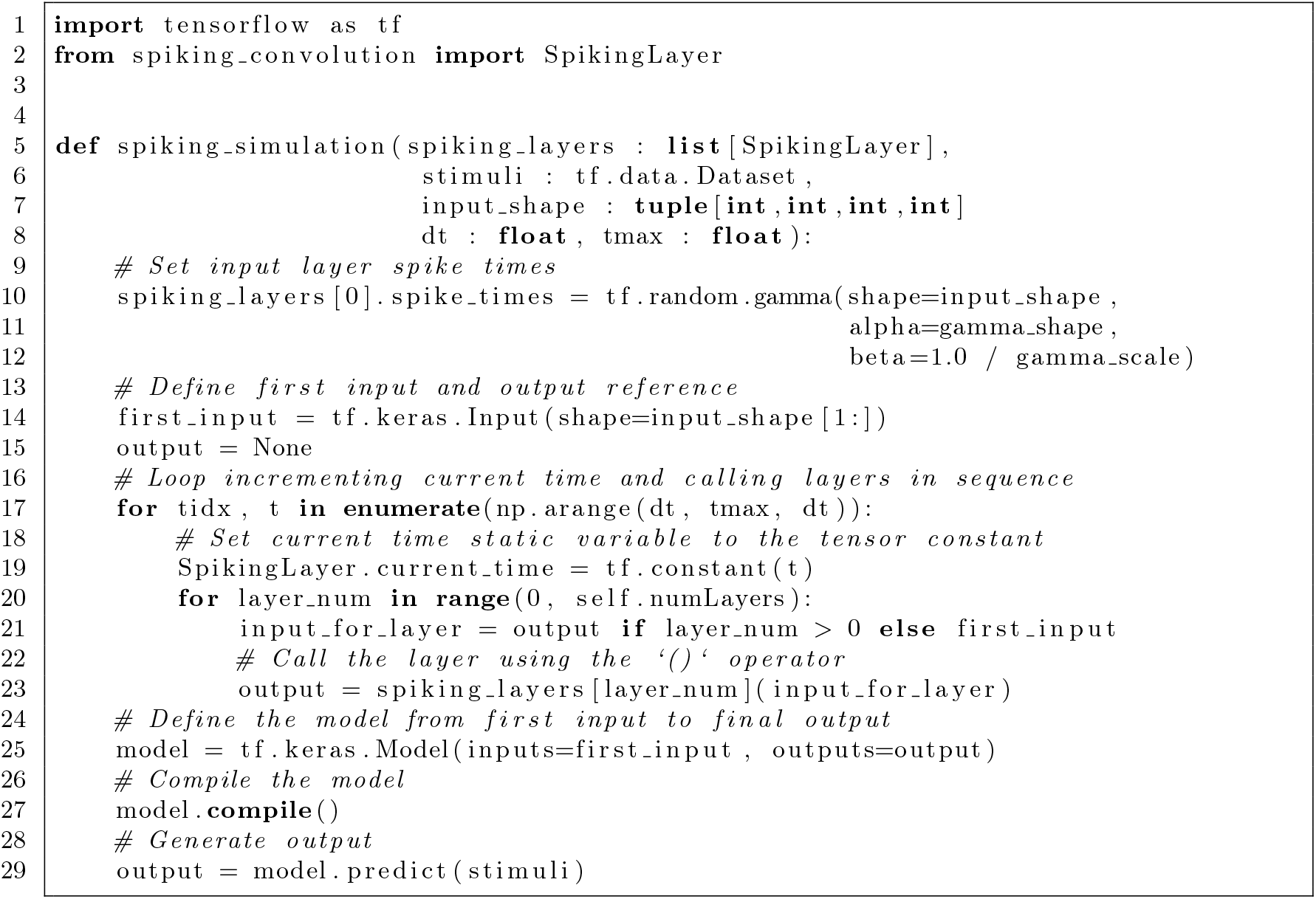

The inhibitory interneuron logic, shown in Listing 4 is a simplified version the excitatory PSP calculation. Note that throughout Listing 4 “IPSP” refers to the constants defined for the inhibitory interneuron itself, not its PSP in the excitatory neuron (which actually follows the same dynamics as the EPSP in the L2/3 pyramidal neuron, as described in Section 4.1.) Transmission for from L4 to L2/3 fast spiking inhibitory interneurons to L2/3 pyramidal neurons was studied in [Hoffmann et al., 2015] which shows that interneurons reliably convert efferent spikes into IPSP’s in the afferent L2/3 pyramidal neurons with an AP to IPSP latency of 1.1 ± 0.5 ms. The model here is a simple PSP dynamics in the interneuron using the time constants of [Holmgren et al., 2003] and ignoring additional delays in the transmission from the inhibitory neuron to the L2/3 pyramidal neurons. As there is only one input to each model inhibitory unit, the only additional parameter is a fixed synaptic weight in the interneuron (applied on line 19, Listing 4). The strength of the input in the interneuron was set to 1.5: A PSP with the time constants of [Holmgren et al., 2003] reaches its maximum at around 3.5 ms, and with the 1.5 *×* scaling is crossed threshold (defined as 1) after 1.35 ms. Note that randomness in the scale if the IPSP in the L2/3 pyramidal neuron is applied in the pyramidal layer. While simple, this model is consistent with the findings of [Hoffmann et al., 2015] and shows how this approach could be used for more detailed studies of interneuron properties in the future. For the present work it is enough to show that realistically delayed but reliable inhibition arriving at the L2/3 pyramidal neurons is consistent with Organic Convolution theory of cortical processing.

After defining the layers, the tensor graph for a complete spiking simulation must be created before passing in inputs and executing the graph, which is shown in Listing 5. The first step in the simulation graph is to set the input times in the first `SpikingLayer` (the input layer) to random times drawn from a gamma distribution. The main tensor graph is created in a double loop over time, and layer (lines 17-20.) The first step in the simulation loop for a point in time is to set the `current_time` class variable to a tensor constant for use in all the layer calculations (Listing 5 line 19.) Then the layers are called in succession with the output of each layer as the input to the next (lines 20-23); note that for `keras.Layer` the `call` function is overloaded with the () operator and does not need to be named explicitly (line 23).

To perform a simulation, a base class `keras.Model` is created from the first input and the final output; the standard `tf.keras.Sequential` model cannot be used for this specialized use case. After creation, the `Model` can be used to generate outputs as usual by first compiling the model, and then generating outputs with the `Model.predict` function.

### 4.4 Convolution Model for Patterns of Afferent Connections

Topographically organized connection patterns are used to determine the convolution filter weights in the model. The patterns are implemented as rules giving the input preference for dendritic locations as a function of the position relative to the soma, using trigonometric functions. A conceptual illustration of such patterns is shown in Figure 19, using a V4 neuron selective to medium convex curvature as an example: V2 contains a topographically organized set of neurons with selectivity to corners including an exhaustive set of corner angles and rotations (Figure 19A and G.) These V2 pyramidal cells are the excitatory afferent inputs of the V4 L2/3 pyramidal neuron via the V4 L4 neurons, which also excite the inhibitory afferent inputs to the V4 L2/3 pyramidal neuron, the L2/3 Fast Spiking inhibitory neurons (Figure 19B). These excitatory and inhibitory afferent inputs to the V4 L2/3 pyramidal neuron arrive in the neuropil around V4 L2/3 pyramidal neuron in a topographically organized manner (Figure 19C.)

The basal dendrites of L2/3 pyramidal neuron extend outward from the direction of the apical dendrite in the plane of the cortical layer, which is the plane of topographical organization (Figure 19F.) With this organization, the dendrites of the L2/3 pyramidal neuron may select stripes of excitation (Figure 19E) against a background of inhibition (Figure 19D.) Similar simple structures can explain the feedforward afferent connections formed by V1, V2 and V4 pyramidal neurons.

For all connection patterns discussed in this study the location of the inputs in the dimension perpendicular to the cortical layers (in the direction of lower and higher cortical layers) is not relevant to the operation of the pattern. Only the layout of afferent inputs in the plane of the topographic map, or parallel to the layers of cortex, is relevant to the functioning described and the model. For this reason the model only represents input layout in a two dimensional plane.

#### 4.4.1 Oriented Pattern Deffnition Overview

In every layer of pyramidal neurons, all of the connection patterns are generated at multiple orientations, and every orientation defines a separate channel in the convolution model. A common pattern generation logic was used to facilitate creation of a large number of filters that are rotationally equivalent. Each channel in the convolution model, representing a pattern at a specific rotation, has a baseline angular orientation in the plane of the topography. The pattern itself is defined by a rule or set of rules determining which afierent input types are excitatory and which are inhibitory based on the radius of a point from the center of the receptive field and angle relative to the neuron baseline. Specific rules used to make the filters in each cortical area will be described in the following sections Section 4.4.2, Section4.4.3 Section 4.4.4 for V1, V2 and V4 respectively. Various measurements of the patterns and the resulting filters are shown in Table 7.

Given a rule defining the pattern, the following process was used to creates the convolution filters for the model: 1.

1. Pick an ellipse defining the receptive field at the baseline orientation; points within the rectangular convolution filter dimensions but outside of the ellipse boundary are set to zero. The semi-major and semi-minor axes of these ellipses are shown as the \Field” in Table 7 and these are also the rectangular size of the convolution filter for the unrotated (rotation 0°_) version of the units.
2. Radii are traced traveling from the center of the receptive field to the edge of the ellipse. These paths can be conceptualized as dendrites, although they are not realistically formed: They are perfectly straight and numerous enough to cover every filter position within the bounding ellipse. The number of dendritic radii paths used to define each pattern’s filters are shown as “Radii” in Table 7
3. The pattern rule is applied at sample points along the dendritic radii to determine the weights for every point in every available input channel. The resulting excitatory and inhibitory connections are converted to locations in the convolution filter.
4. For each rotated version of the pattern (having a different orientation selectivity) the ellipse and all of the radii sample points are rotated to the new orientation. The smallest rectangle bounding the rotated ellipse defines the boundary rectangle for the convolution filters for that rotated version of the pattern. Rotated versions of the patterns were generated every 22.5^°^, resulting in 8 rotations for the patterns with point symmetry and 16 for the patterns that are asymetric.

The size and rules for the excitatory and inhibitory connections in the patterns for each cortical area were designed to match the theorized function of each area of cortex on the scale of the model stimuli. As will be described in detail below, rotation of the pattern rules for each type of connection pattern in V1 and V2 resulted in a small number of unique variants of the filter layouts that repeated at 90^°^ rotations. Although the V1 and V2 units represent from 8-16 orientation selectivities through rotation of the pattern, there were only 2-3 unique filter layouts that appear 2-4 rotations each to make up the entire 8-16 filters. The size, number of rotations, number of unique variants, and the number of weight parameters for the connection patterns are summarized in Table 7.

For simplicity, in every pattern the weights on any given afferent channel are allowed only a single excitatory value and a single inhibitory value for each variant of a pattern. So in any single convolution filter there are only 3 possible weights: *w*_*excite*_, *w*_*inhib*_ and 0. Different variants of each pattern can have *w*_*excite*_ and *w*_*inhib*_ set differently as part of the weight tuning process described in Section 4.5, but every variant must have the same **A** weight parameters (identical filters) for all of it’s rotations. As shown in table 7, the total number of tunable weight parameters in the model is fairly small: a total of 73 tunable weight parameters serves to define the weights for a total of 242 convolution channels, representing 21 patterns of connection that are rotated using 34 unique variants.

Table 7 also summarizes the total number of other parameters that define each pattern in the column *Non-Wt Params*: These include 2 parameters for the filter size (except for the inhibitory and pooling units whose size is fixed by the model structure). and some other parameters such as the bandwidth of the stripe selective units in V1 and the integration of interior angles to form curvature selectivity in V4. Table 7 also shows the total number of tunable parameters for the convolution filters, which 141. For comparison, an early successful convolution model for black and white handwriting is LeNet-5 [LeCun et al., 1998]. This network also has 3 convolution layers interspersed with pooling layers. In contrast, LeNet-5 has 154 tunable parameters in its first convolution layer alone, followed by 1,516 in the second convolutional layer and 48,120 parameters in the third convolution layer for a total of 49,792, around350*×* the tunable parameters in the organic convolution model in this study. For the OC model, the total number of filter elements in every channel of every convolution filter is 45,912 - very close to LeNet-5 (the product of the filter sizes and the number of channels in Table 7). Thus a theoretical function based approach to determining the form of convolution filters provides around 350 *×* compression in the free parameters necessary to implement the filters.

#### 4.4.2 V1 Model Connection Patterns

The V1 orientation selective model units are based on decades of research beginning with [Hubel and Wiesel, 1962] that found that V1 contains units selective to orientation in a variety of forms. This model instantiates two forms, selective to oriented bands or stripes and oriented edges. The afferent connection patterns for V1 L2/3 units are extremely simple and still reproduce the complex shape tuning of V4 recordings. The oriented stripe selective filters are shown in Figure 20A : The 8 rotations of this filter form a total of 3 unique variants of the filter layout. In the horizontal and vertical variant (1st column of Figure20A), the filter consists of a 2 *×* 5 excitatory strip flanked by 1 *×* 5 inhibitory strips; the entire filter is 5 *×* 4. When the V1 band selective pattern is rotated by 22.5^°^, it forms two more variants as 5×5 filters: One variant for the 22.5^°^ rotation from the horizontal, and 3 more rotations of the same pattern (2nd and 4th columns of Figure 20A); and one variant for the 45^°^ and 135^°^ rotations (3rd column of Figure 20A).

The oriented stripe selective filters in the V1 model are created by applying the following rules: Given that every unit has an orientation baseline, the oriented stripe selective units also have a width parameter defining the central region of afferent connections. During connection formation based on position angle and radius in the receptive field (as described in Section 4.4.1), an afferent input is excitatory whenever the absolute value of the distance multiple by the sine of the angle with the neuron’s baseline is less than the width parameter. That is, a point labeled *i* is excitatory whenever |*r*_*i*_ sin(*θ*_*i*_)| *< b* where *r*_*i*_ are the radius to the point, *θ*_*i*_ is the angle to the point relative to the neuron’s baseline angle, and *b* is the width parameter. For the horizontal and 45^°^ variants a band width of *b* = 1 was used, while for the 22.5^°^ variant a band width of *b* = 0.8 was used. The rules resulted in the excitatory and inhibitory layouts of Figure 20A.

The filters for the edge selective V1 units are shown in Figure 20B : The 16 rotations of this filter also come from 3 unique variants of the filter layout: The horizontal/vertical variant is 4×8 (1st column of Figure20B); the 22.5^°^ rotation variants are 5 *×* 8 (2nd and 4th columns of Figure20B); the 45° rotations are 7 *×* 7. Like the oriented stripe selective units, each variant has a single parameter for the excitatory weight and a single parameter for the inhibitory weights and there are a total of 6 weight parameters for the V1 edge selective units as shown in Table 8. Further explanation of how these excitatory and inhibitory weights is discussed below in the Section 4.5.

For the edge selective filters, the connection formation rule for the pattern is simply that an input afferent is excitatory whenever the the point, labeled *i*, meets the condition that *sin*(*θ*_*i*_) *>* 0 where *θ*_*i*_ is the angle to the position relative to the neuron’s baseline. As shown in Figure 20B, for points lying on the *sin*(*θ*_*i*_) = 0 neither excitatory nor inhibitory connections are made.

As described above, each variant has a single parameter for the excitatory weight and a single parameter for the inhibitory weights and there are a total of 6 weight parameters for the V1 line selective units as shown in Table 8. The three variants of each type of unit are labeled by the first orientation at which they appear : 0^°^, 22.5^°^ and 45^°^.

For the V1 stripe and edge selective units the inhibitory weight is stronger than the excitatory weight, as can be seen in table 8. For all of the V1 units the inhibitory weight is set so that 40% of the total input weight is excitatory, and the remaining 60% is inhibitory. This ratio of excitatory weight to the total weight will be referred to as *ρ*_*E*_ and is used in every pattern to set the inhibitory/excitatory balance in an intuitive manner. Setting *ρ*_*E*_ *<* 0.5 prevents spurious firing of the units when the background is solid. Further explanation of how these excitatory and inhibitory weights were set is discussed below in the Section 4.5.

Note that having 8 rotations of the V1 stripe selective pattern 22.5^°^ apart is intended as a discretization of the continuous range of orientations to which V1 L2/3 neurons respond, and not to necessarily imply that there are 8 distinct sub-types of V1 L2/3 oriented stripe receptive field neurons. The same interpretation applies to oriented versions of the various connection patterns throughout this study. Similarly, the perfect symmetry of the connection pattern is intended as a typical or average unit (see the Discussion section for additional considerations.)

#### 4.4.3 V2 Model Connection Patterns

The V2 model is based on the findings in [Liu et al., 2016] and the V2 L2/3 units in the model implements both ultralong and complex-shaped selectivities described therein. The conceptual model for how such patterns may form was illustrated in Figure 19 and described in the accompanying text in Section 4.4.

For V2, one underlying connection pattern pattern can explain selectivity to both ultralong and complex shaped corner selectivity: It is theorized that these V2 units have a small number of basal dendrites receiving feedforward excitation, and those dendrite extend radially outward in the topographically organized cortical plane, as described in Section 4.4. These dendrites with excitatory connections may prefer V1 afferents that have the same preferred stimulus orientation as the physical direction of the dendrite within the topographic arrangement. That is, a dendrite which extends in the horizontal direction in the topographic map of afferent inputs will prefer connections to horizontally oriented V1 afferents; a dendrite with excitatory connections that is positioned in the plane of the topography at 45^°^ from the horizon will prefer V1 afferents with orientation selectivity 45^°^ from horizontal, etc.

If a V2 unit has two such dendrites or clusters of dendrites that extend in a direction from the soma that is substantially opposite to each other, they will form an ultra-long receptive field. The model includes 8 different orientation preferences for ultralong units, oriented 22.5^°^ apart. As in the V1 model, the different angle and orientation versions are intended to represent a discretization of a the range of neurons that would be found in vivo and do not necessarily imply distinct neuron sub-types in nature. The V2 model ultralong units are modeled with 7 *×* 7 convolution filters which are shown in Figure 21**A**-**C**.

To match the recording results of [Nandy et al., 2013] for the low curvature selective V4 units the V2 ultralong units need broad tuning: an excitatory input aligned to the preferred orientation of the pattern and also excitatory inputs from the two neighboring afferent input orientations +/-22.5^°^ offset from the primary. Figure 21A also shows that there are three unique variants of the filter set: Horizontal/vertical oriented with weaker +/-22.5^°^ offset excitation (Figure 21B row 1); 22.5^°^ oriented with additional inputs from the horizontal and 45^°^ afferents (Figure 21B row 2); and 45^°^ oriented with additional inputs from the 22.5^°^ and 67.5^°^ oriented afferents (Figure 21B row 3). Additional rotations of these three variants form the complete set of 8 different ultralong units in the V2 model. (For example, row 4 of Figure 21B is the 67.5^°^ selective version and is a rotated version of row 2, the 22.5^°^ selective version.)

The pattern connection rule used for creating the V2 ultralong unit is as follows: As described in Section 19, after determining the neurons bounding ellipse, as set of paths are drawn from the center of the receptive field to the edge. For the V2 ultralong unit 32 paths are used, spaced equally 12.5^°^ degrees apart, beginning from each neurons baseline orientation. Excitatory connections are formed to the afferent connection with orientation selectivity angle matching the orientation of the V2 ultralong unit for all points on the path of the baseline angle, and excitatory connections are also formed on the path 180^°^ from the baseline angle. All other orientation of afferent inputs and all other positions along the connection sampling paths form inhibitory connections.

The convolution filter weights for the V2 ultra-long model units are shown in Table 9 listed as 180^°^ interior angle (straight). The three variants of the V2 ultralong unit are labeled by the first orientation at which the variant is used. Because in the V1 model there are only 3 unique variants covering all orientations of selectivity, the V2 model filters were constrained that each variant of the ultra-long unit would have a single weight for all afferents from the same V1 variant. The weights for each variant are labeled by the angle of the afferent V1 variant from as given in Table 8: 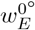 refers to the excitatory weights from the 0^°^ variants from V1 and likewise for 22.5^°^ and 45^°^. (In fact, the inputs to V2 are the pooled combination of the stripe and edge variants listed in Table 8, where the 0^°^ stripe was pooled with the 0^°^ and 180^°^ edge, etc.) This constraint serves as both a logical regularization technique: Because it is assumed that the V2 units ought to have a rotationally symmetric response to rotated stimuli, and as these afferent V1 channels will have identical response at each rotation of their variant, they should be equally weighted.

**Table 9.** V2 Complex Pattern Afferent Input Weights by Interior Angle and Angle of First Rotational Variant. 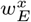: Excitatory input strength per afferent input variant *x*. Missing values indicates that no such variant is present in the model. *ρ*_*E*_ : Percentage of t otal afferent input strength that is excitatory. *w*_*I*_ : Inhibitory strength per afferent input. *w*_*E*_: Total excitatory weight. *w*_*I*_ : Total inhibitory weight. All weights are in units where 1= spike threshold.

| Interior Angle | Variant | $w_E^{0^\circ}$ | $w_E^{22.5^\circ}$ | $w_E^{45^\circ}$ | $\rho_E$ | $w_I$ | $\sum w_E$ | $\sum w_I$ |
| --- | --- | --- | --- | --- | --- | --- | --- | --- |
| 45° | 0° | 0.198 |  | 0.221 | 0.42 | -0.005 | 1.9 | -2.6 |
| 45° | 22.5° |  | 0.195 |  | 0.48 | -0.004 | 2.0 | -2.2 |
| 67.5° | 0° | 0.178 | 0.180 |  | 0.49 | -0.003 | 2.0 | -2.2 |
| 67.5° | 45° |  | 0.167 | 0.215 | 0.52 | -0.004 | 1.8 | -1.7 |
| 90° | 0° | 0.215 |  |  | 0.52 | -0.005 | 1.7 | -1.6 |
| 90° | 22.5° |  | 0.218 |  | 0.36 | -0.009 | 1.7 | -3.1 |
| 90° | 45° |  |  | 0.260 | 0.57 | -0.003 | 1.6 | -1.2 |
| 112.5° | 0° | 0.182 | 0.140 |  | 0.54 | -0.003 | 1.6 | -1.4 |
| 112.5° | 22.5° |  | 0.199 | 0.246 | 0.37 | -0.006 | 2.0 | -3.3 |
| 135° | 0° | 0.144 |  | 0.157 | 0.37 | -0.004 | 1.4 | -2.3 |
| 135° | 22.5° |  | 0.154 |  | 0.52 | -0.003 | 1.5 | -1.4 |
| 157.5° | 0° | 0.11 |  | 0.13 | 0.49 | -0.002 | 1.2 | -1.2 |
| 157.5° | 45° |  | 0.1 | 0.15 | 0.55 | -0.002 | 1.1 | -0.9 |
| 180° | 0° | 0.11 | 0.13 |  | 0.47 | -0.009 | 2.6 | -2.9 |
| 180° | 22.5° | 0.11 | 0.12 | 0.23 | 0.44 | -0.008 | 2.7 | -3.4 |
| 180° | 45° |  | 0.1 | 0.23 | 0.48 | -0.008 | 2.6 | -2.9 |

The inhibitory weights for the V2 units are constrained so that for each variant all inhibited afferent inputs have the same weight. The inhibitory weights are described by two parameters in Table 9: *ρ*_*E*_ is the total percentage of afferent input that is excitatory, and *w*_*I*_ is the resulting weight per afferent input. Note that although a large proportion of the overall weight is inhibitory, the inhibitory strength per input is small. This is due to the large number of afferent input types and high selectivity of the excitatory input by position and preferred orientation. Details of how the excitatory and inhibitory weights were tuned are described in Section 4.5. Given identical excitatory weights per input V1 variant and one parameter controlling inhibition per V2 variant, there are a total of 10 weight parameters from the 3 unique variants of the ultralong as shown in Table 9.

The V2 complex units, which are corner selective, follow the same approach as for the ultralong: It is assumed that a neuron may have two dendrites or cluster of dendrites with the connection preference that their afferent orientation selectivity corresponds to the direction of the dendrite(s) in the topographic map. In the case where those two dendrites have some angle to each other in the plane of cortex, which is the plane of the topographic map, the result will be a complex-shaped receptive field approximating a corner. The corner selectivity results from the radial arrangement of the basal dendrites: The dendrites or clusters of dendrites with these preferences for V1 afferents must meet at the soma which becomes the location of the vertex of the corner. These units prefer stimuli that resembles two lines meeting in a corner, and the angle between the two excitatory dendrites sets the angle preference for the unit. The model includes a range of angles defining the separation between the sub-units of the complex-shaped units, from 45^°^ to 157.5^°^ in 22.5^°^ increments for a total of 6 complex-shaped subunit angles. Because the complex-shaped units are not point symmetric (to 180^°^ rotations), a total of 16 different orientations (22.5^°^ apart) of each sub-unit separating angle are defined to respond to the full range of possible stimuli.

The 9*×* 9 convolution filters resulting from the 90^°^ interior angle complex receptive field V2 L2/3 unit connection pattern is illustrated in Figures 21**D**-**F**: Excitatory connections are concentrated on a few orientations in straight line arrangements. Inhibition is diffuse and spread over all other channels throughout the receptive field. Figure 21**E** shows that for the 90^°^ interior angle selective unit there are three unique variants of the filters: The filter for 0^°^ orientation (Figure 21E row 1) repeats in rotated form for 90^°^, 180^°^ and 270^°^ orientations; the filter for the 45^°^ orientation (Figure 21E row 3) repeats in rotated form for 135^°^, 225^°^ and 315^°^; and the filter for the 22.5^°^ orientation (Figure 21E row 2) repeats 8 times for the remaining orientations of the unit. As in the model of the ultralong unit, independent weights are only used on afferent channels from distinct V1 variants and as a result each 90^°^ interior angle subunit has just one excitatory weight and one inhibitory weight. For example, the 0^°^ variant (Figure 21E row 1) has inputs from the 0^°^ V2 L4 unit (column 1) and the 90^°^ V4 L4 unit (column 5); these correspond to the same V1 variant (Figure 20 column 1) and hence they all will receive the same excitatory weight 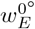 in the V2 model. This approach results in a total of 6 independent weights for the V2 L2/3 pyramidal model with 90^°^ interior angle, as shown in Table The specific weight values are listed in Table 9. Details of how the weight values were determined are explained in Section 4.5.

The convolution filters for the other interior angles of the V2 complex units ares summarized in Figure 22. Just as in the 90^°^ interior angle unit, excitatory connections are concentrated on two orientations in straight line arrangements, and there is diffuse inhibition to all other input afferent types not shown in the Figure. Each type of V2 complex unit shown in Figure 22 has two variants. In every case these two variants each occur at 8 rotated orientations to make up the full set of oriented units. Each interior angle unit shares weight for each of the three variants of input afferent from V1, so each V2 complex unit has a total of 5 or 6 weight parameters as summarized in Table 7. The specific weight values are listed in Table 9 and the details of how these values were determined are explained in Section 4.5.

**Fig. 22.**
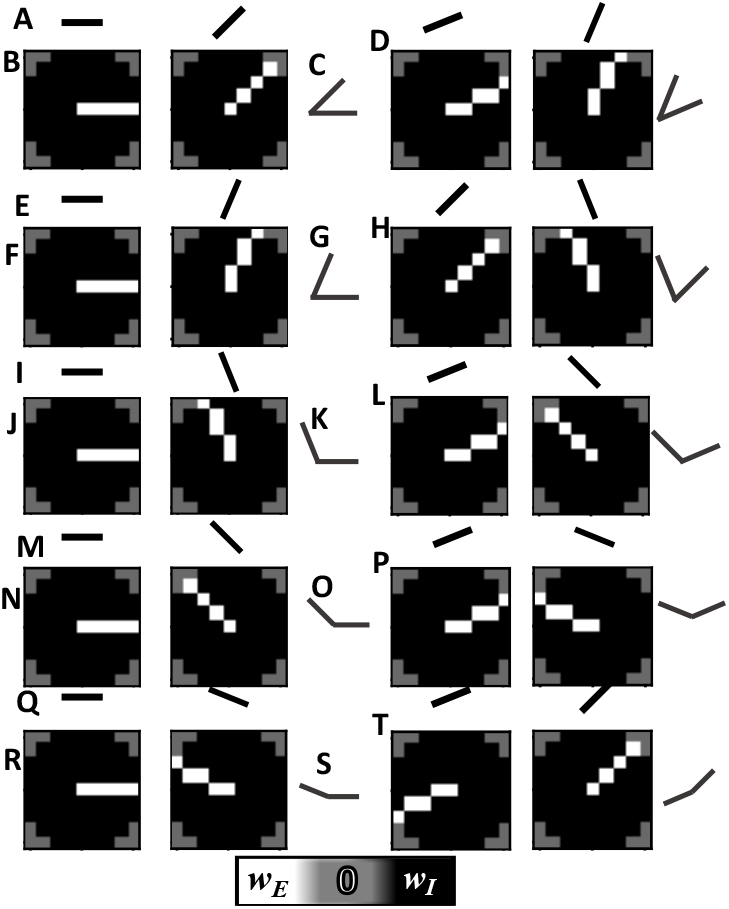
Afferent Connection Patterns for model V2 L2/3 corner selective unit variants. The convolution pattern represents the afferent connections to excitatory and inhibitory inputs as illustrated in Figure 19. For each interior angle, the two variants are shown; the 8 rotated versions of these form the 16 sets of filters representing each unit type. **A**-**D** Corner Selective Unit with 45^°^ interior angle. **E**-**H** Corner Selective Unit with 67.5^°^ interior angle. **I**-**L** Corner Selective Unit with 112.5^°^ interior angle. **M**-**P** Corner Selective Unit with 135^°^ interior angle. **Q**-**T** Corner Selective Unit with 157.5^°^ interior angle. The single bar pictograms **A**,**E**,**I**,**M**,**Q** represent the orientation selectivity of V1 afferent afferents for the first variant. Two tone heatmaps **B**,**F**,**J**,**N**,**R** show the pattern of excitatory (white), inhibitory (black) and zero (gray) weights in the convolution filters. Afferent input orientations not shown are inhibited as in Figure 21. The corner bar pictograms **C**,**G**,**K**,**O**,**S** show the resulting preferred stimuli of the V2 unit. The second rotational variant is shown for each interior angle in **D**,**H**,**L**,**P** and **T**. The specific values used for the excitatory and inhibitory weights for each type of unit are shown in Table 9.

The pattern connection rule for creating the complex V2 units is similar to that as for the ultralong unit, albeit taking into account the two different orientation selectivities for each arm of the corner: A total of 32 paths space 12.5^°^ apart are drawn in the bounding ellipse. Excitatory connections are made to afferents with orientation selectivity matching the neuron baseline orientation along the path that has the baseline orientation. A second path separated from the baseline by the interior angle of the corner selectivity forms excitatory connections to afferents that have selectivity for orientations that are also the interior angle away from the neuron baseline angle. All other orientation of afferent inputs and all other positions along the connection sampling paths form inhibitory connections.

#### 4.4.4 V4 Pattern Details

The model V4 connection patterns to their afferent inputs from V2 was introduced in the Results Sections 2.1, 2.2 and 2.3. This section provides additional details. As described in Section 4.4.1, the convolution filters are determined by calculating a set of dendritic radii paths in an ellipse bounding the receptive field and applying a connection rule at each point along the paths. For the V4 units selective for curvature, the rule is to make excitatory connections to afferent inputs that meet three criteria:

1. The interior angle of the afferent V2 unit must be within a specified range for each V4 unit type. The range is shown in Table 10, where *α*_*min*_ is the minimum interior angle and *α*_*max*_ is the maximum interior angle accepted as afferents with excitatory connections. This range defines whether the V4 unit is selective to acute, medium or straight/nearly straight curvature.
2. There may be only a specified percentage of the central angle within which the receptive field forms excitatory connections, *γ*_*E*_ in Table 10. This differentiates the units selective for curvature in a position in the receptive field reported in [Pasupathy and Connor, 2001] from those selective for curvature throughout the receptive field in [Nandy et al., 2013]. For the V4 units selective to curvature in a position *γ*_*E*_ ranged from 15% to 25% and for the V4 units with curvature selectivity throughout the receptive field *γ*_*E*_=100%.
3. The orientation selectivity of afferent input with excitatory connections equals the radius path orientation plus a constant offset for each type of V4 unit and afferent input interior angle. Therefore the orientation of the afferent input with excitatory connections rotates with position around the receptive fields. These offsets are shown in Table 11.

**Table 10.** V4 Pattern Parameters and Weights. *α*_min_ : Minimum interior angle of V2 afferent to form excitatory connections. *α*_max_ : Maximum interior angle of V2 afferent to form excitatory connections. *γ*_*E*_ : Percentage of the central angle within which the receptive field forms excitatory connections. *w*_*E*_ : Excitatory weight per afferent input . *ρ*_*E*_ : Percentage of the total afferent input weight which is excitatory. *w*_*I*_ : Inhibitory weight per afferent input. ∑*w*_*E*_ : Total excitatory weight. *∑w*_*I*_ : Total inhibitory weight. All weights (*w*) are in units where 1= spike threshold.

| Pattern | $\alpha_{\min}$ | $\alpha_{\max}$ | $\gamma_E$ | $w_E$ | $\rho_E$ | $w_I$ | $\sum w_E$ | $\sum w_I$ |
| --- | --- | --- | --- | --- | --- | --- | --- | --- |
| Acute Convex Position | 45° | 67.5° | 15% | 0.29 | 0.7 | -0.002 | 56.6 | -24.3 |
| High Curve Full Field | 67.5° | 112.5° | 100% | 0.33 | 0.6 | -0.235 | 3120 | -2073 |
| Medium Convex Position | 90° | 135° | 25% | 0.25 | 0.6 | -0.004 | 115 | -77 |
| Medium Concave Position | 90° | 135° | 15% | 0.67 | 0.7 | -0.004 | 196 | -83 |
| Medium Curve Full Field | 90° | 135° | 100% | 0.33 | 0.7 | -0.04 | 986 | -423 |
| V4 Low Curve | 157.5° | 180° | 100% | 0.11* | 0.8 | -0.015 | 227 | -57 |

**Table 11.** V4 Pattern Afferent Angle Selectivity Rotation from V2 Orientations.

| Pattern | $\beta^{45^\circ}$ | $\beta^{67.5^\circ}$ | $\beta^{90^\circ}$ | $\beta^{112.5^\circ}$ | $\beta^{135^\circ}$ | $\beta^{157.5^\circ}$ | $\beta^{180^\circ}$ |
| --- | --- | --- | --- | --- | --- | --- | --- |
| V4 Acute Convex | -157.5° | -146.25° |  |  |  |  |  |
| V4 Medium Convex |  |  | -135° | -123.75° | -112.5° |  |  |
| V4 Medium Curve |  |  | -135° | -123.75° | -112.5° |  |  |
| V4 Concave |  |  | 45° | 56.25° | 67.5° |  |  |
| V4 High Curve |  | 168.75° | 180° | -168.75° |  |  |  |
| V4 Low Curve |  |  |  |  |  | -11.25° | 0° |

All excitatory inputs for a single V4 unit type have a single weight parameter, *w*_*E*_ in Table 10. Further,

- Afferents inputs from V2 having interior angle within the range [*α*_*min*_, *α*_*max*_] that are not excitatory according to the above rules are inhibitory at a constant weight, *w*_*I*_ in Table 10. Like the V1 and V2 models, *w*_*I*_ is controlled through a more intuitive parameter *ρ*_*E*_ which sets the total percentage of afferent weight that is excitatory.
- Afferents from inputs where the interior angle is outside of the range [*α*_*min*_, *α*_*max*_] have zero weight - they are neither excitatory nor inhibitory.

The offsets from V2 unit orientations to V4 afferent inputs, shown in Table 11 translate from the orientation convention of the V2 units to the orientation of V4 dendritic radii required for the V4 pattern selectivity: In the corner selective units in V2, the convention is that a unit’s orientation is defined by the first excitatory radius path, and the other excitatory radius path is rotated by the interior angle counter-clockwise - these are the units shown in Figure 22 **B**,**F**,**J, N** and **R**. The offsets from V2 unit rotation to V4 radius paths are denoted as *β*^*δ*^ in Table 11 where *δ* is the interior angle of V2 corner selective unit

The V4 units selective for acute and medium convex curvature in a position and medium convex curvature throughout the receptive field apply a simple translation so that the excitatory afferent connections are formed with inputs whose selectivity is for a corner that is perfectly convex with respect to the V4 receptive field. For example,

Table 11 row 1 shows that the rotations for the V4 acute convex unit begin at *β* = − 157.5^°^ for the 45^°^ interior V2 unit; this which means that the 45^°^ interior angle unit having its first excitatory arm at 202.5^°^ is perfectly convex to a horizontal radius path. For the V4 acute and medium convex units *β* decreases by 12.25^°^ for every 22.5^°^ increase in the interior angle *δ* of the V2 corner selectivity. This arrangement keeps the excitatory inputs convex to the center of the receptive field as the position rotates around the V4 unit receptive field.

In the concave curvature selective unit the offsets are rotated by precise 180^°^ from those used for the acute convex and medium curvature selective unit - this makes the excitatory afferent inputs precisely concave at every position in the receptive field. For V4 unit selective for high curvature throughout the receptive field (“V4 High Curve’ in Table 11), to better match the recordings the offset had to be reduced by 45^°^ from the perfect convexity used by the acute convex position selective and medium curvature selective units. For the V4 high curvature selective unit, the excitatory inputs do not seem to be perfectly convex, but seem to be offset somewhat - this can be observed in Figure 5.

For the low curvature selective V4 unit based on [Nandy et al., 2013] the pattern connection rules were different from the rules for the curvature selective units:

1. Excitatory connections based on orientation were constant throughout the receptive field and matched the baseline orientation of the V4 unit. A constant rotational offset of just 11.25^°^ was applied to the 157.5^°^ interior angel corner (nearly straight) V2 unit connections to align them with ultralong V2 units and the orientation selectivity of the V4 unit.
2. For excitatory connections to ultralong afferent inputs from V2 there is a broad tuning so that inputs that have orientation selectivity 22.5^°^ and 45^°^ away from the preferred orientation also made excitatory connections. For inputs from the 157.5^°^ interior angel corner (nearly straight) V2 unit only the two orientations matching the preferred orientation formed excitatory connections (including the 11.25^°^ offset on the orientation, there are two 157.5^°^ interior angel corners that are equally close to the preferred orientation. See Figure 2.3 for illustration.)
3. The strength of excitatory connections to afferent ultralong inputs depended on the afferent unit’s orientation selectivity. The weight shown in Table 10 was assigned to the ultralong input that has the preferred orientation and the connection strength dropped by 25% for every 22.5^°^ that the V2 afferent selectivity differed from the V4 unit preferred orientation: The weights on the inputs 22.5^°^ from the preferred orientation were 75% of the weight on the preferred orientation, and weights on the inputs 45^°^ from the preferred orientation were 50% of the weight on the preferred orientation.

The convolution filters resulting from these rules were presented previously in the Results sections as follows:

- Acute Convex Position: Figure 4, Section 2.1
- Medium Convex Position: Figure 8, Section 2.2
- Concave Position: Figure 8, Section 2.2
- High Curve: Figure 6, Section 2.1
- Medium Curve: Figure 10, Section 2.2
- Low Curve: Figure 12, Section 2.3

#### 4.4.5 Layer 4 Pooling Models

The first layer of the model is V1 L4 which serves as an input layer to the neural network and is used as a lumped model for the eye/retina, LGN path up to V1 L4 [Vanni et al., 2020]. In this layer, the spiking probability on every trial is equal to the input intensity as a percentage of the maximum. This layer is distinct, while the V4 layers in V2 and V4 follow a common pattern.

V2 L4 plays the role of receiving excitatory input and passing it to both V2 L2/3 excitatory and inhibitory units. The L4 model for V2 also performs a pooling function: The units have 2*×* 2 receptive fields and are spaced 2 positions apart. This is known as a stride of 2 in the terminology of DCNN, and results in the dimension of the output being reduced by one half along each axis: The input to model V2 L4 is 112*×* 112, and the output is 56*×* 56.

The V2 L4 units have 8 orientations corresponding to the orientations of the V1 units orientation preferences, every 22.5^°^. (The instantiated channels are intended as a discretization of the range of possible orientation preferences and not a suggestion that there are 8 distinct unit types in area V2.) Each orientation preference of V2 L4 unit accepts inputs from one channel of the V1 L2/3 oriented band channels, and also the two corresponding edge selective channels. In terms of individual units there are a total of 12 inputs for each V2 L4 unit after pooling and combining the three different inputs. These inputs are weighted sufficiently so that any one of the 12 inputs from V1 to an area V2 L4 unit will trigger an output spike. The result is a spiking version of the maximum pooling operation in a DCNN.

The model for V4 includes a model of L4 that has the same function as the V2 L4 model, adapted to the V2 afferents: V4 L4 units take afferents from a 2*×* 2 pixel receptive field, and are staggered 2 pixels apart and the dimension of the visual maps are reduced from 56 *×* 56 to 28 *×* 28. Each unit in the V4 L4 model accepts afferent inputs from a single type (channel) of the V2 connection pattern in terms of both sub-unit separation angle and the unit’s overall orientation. Just like the V2 L4 units, the V4 L4 units have sufficient excitation so that they are guaranteed to spike if any of their four inputs from V2 units spike.

#### 4.5 Model Weight Parameter Tuning

The form of convolution filters are determined by the model theory of visual processing at the scale of the input stimuli. The precise values for the afferent connection weights for every filter had to be determined to match the recordings. As shown in Table 7 there are at total of 12 weight parameters for the V1 model, 34 for the V2 model, and 12 for the V4 model. This does not include weights on L4 and pooling units, for which the weights were hard coded by design. At a high level, the weights for V1 and V2 were tuned via stochastic optimization against the following two objectives:

1. The V2 L2/3 corner selective units respond reliably to the stimuli they prefer and as little as possible to others.
2. The V1 edge and stripe unit weights should be set such that they produce equivalent responses in later layers, when an equivalent stimuli is presented as either an outline or a silhouette.
3. The V2 L2/3 ultralong (straight selective) units respond primarily to the preferred orientation and the response decays with the angle from the preferred orientation.

The optimizations were performed using Optuna, an open source hyper-parameter tuning package [Akiba et al., 2019] that implements the Covariance Matrix Adaptation Evolution Strategy (CMA-ES) [Hansen, 2006]. The weights for V4 were tuned manually following simple heuristics.

The process of tuning the weights was performed in 3 phases:

1. Simultaneously tune the weights for the V1 units and the V2 unit with 45^°^ interior angle unit (the most acute) with the optimizer.
2. Tune the weights for all of the other V2 units from 67.5^°^ up to 180^°^ (ultralong) using the optimizer. These units are tuned separately, and in parallel, taking the V1 unit weights as fixed.
3. Manually tune the weights for the V4 units.

The following sections detail these methods: Section 4.5.1 and Section 4.5.2 explain the optimization targets for V2 and V1 respectively. Section 4.5.3 explains the simultaneous optimization of V1 and V2 45^°^ interior angle units in a combined optimization, and the tuning of the remaining V2 corner selective units; Section 4.5.4 details tuning of the V2 ultralong (straight selective) units; and Section 4.5.5 explains the tuning of the V4 L2/3 units.

#### 4.5.1 Optimization Target for V2 Corner Selective Unit Responses

The goal of V2 L2/3 pyramidal unit weight optimization is that the units are focused on their preferred stimuli - they should respond reliably to the preferred stimuli, and as little as possible to other non-preferred stimuli. Achieving this is not easy because the stimuli are composed of a common underlying set of straight oriented segments and because the inputs are highly noisy from a combination of sources (input layer failures, variable PSP amplitudes, and random timing delays at every stage as described in Section 4.1). This noise means that a unit may both fail to respond to its own preferred stimuli if its input weights are not strong enough, but it may respond to non-preferred stimuli if its input weights are too high.

This phenomena is demonstrated in the illustrations of the complete network response in the Appendix Section A Figure A1 and Figure A2. Figure A1 show the response of the entire network to a straight horizontal line stimuli. For the straight line stimuli, the strongest response of any V2 unit is the ultralong unit selective to horizontal orientation - the ultralong units whose receptive field lies along the middle of the line stimuli spike 100% of the time (Figure A1 row C, column 9). However, every other type of V2 unit also spikes at least a small percentage of the time and some spike half the time or more (e.g. Figure A row Q, columns 1, 7, 9 and 15). Specifically, V2 corner selective units in which one arm of the preferred stimuli is aligned on the horizontal arm are likely to spike some of the time, whenever noise firing in the inputs for the other arm excite unit above threshold.

Spurious responses to non-preferred stimuli are not limited to straight line stimuli: Figure A2 shows the response of the entire network to a right angle corner with arms at 0^°^ and 270^°^ to horizontal. (See Section 4.6.2 for details of the creation of the corner stimuli described in this section.) The most reliable response to the stimuli is the properly aligned V2 L2/3 unit responsive to right angle corners which spikes 100% of the time as shown in column 13, row M in Figure A2. At the same time many other units respond. There are two types of spurious responses: First, any unit with an arm at *either* 0^°^ *or* 270^°^ may respond when it is aligned with the stimuli and excitatory noise triggers the non-aligned arm. For example, see the responses of the unit with a 45^°^ interior angle on row Q, columns 3, 5 7 and 9. Second, similar or same interior angle corners that are slightly offset from the preferred orientation may also respond some of the time. For example, see columns 12 and 14, row M in Figure A2, which are the right angle corners oriented 22.5^°^ from the preferred orientation. The most strong spurious response in the network combines both of these conditions: The 112.5^°^ angle corners that share one arm each with the stimuli fire with nearly the same regularity as the aligned right angle corner, as shown in row K columns 12 and 13 of Figure A2.

With these considerations in mind, the weights of the V2 units were optimized against an error target which was a weighted combination of a term for firing less than 100% of the time to the preferred stimuli and a term for firing to any non-preferred stimuli. To tune the weights of each V2 corner selective unit a dataset was made consisting of 20 samples each of:

1. 16 rotations of a corner stimuli having the same interior angle as the V2 unit being optimized, like the 90^°^ corner shown in Figure A2A.
2. 8 rotations of the the straight line stimuli, like that shown in Figure A1A There is one stimuli in the center of the topographic map used to tune the weights which apply to the entire channel, so the maximum firing rate of any unit was used to determine the response of a given unit type to each stimuli.

To achieve the goal, the target is the mean square errors of the difference between firing at 100% to preferred stimuli and 0% to the non-preferred stimuli, with the two components calculated separately and weighted. Mathematically, let 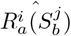 denote the maximum firing rate of any V2 unit with orientation *i* and interior angle *a* (including 180^°^) to a test stimuli with orientation *j* and interior angle *b*. Then the error optimization target to be minimized for a V2 unit with interior angle *a* is defined as:

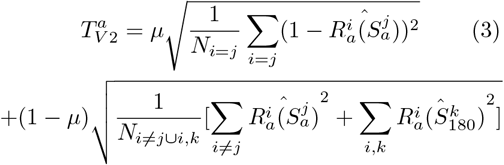

The parameter *µ* controls the weight given to ensuring the preferred stimuli triggers the right units to fire 100% of the times versus ensuring that they do not fire to similar, non-preferred stimuli. After some experimentation, the model tuning for the final study used *µ* = 0.5 in Equation 3.

#### 4.5.2 Optimization Target for V1 Edge and Stripe Units

A second aim of the weight optimization target is designed to ensure consistency of the response to outline and silhouette stimuli. As the data of [Pasupathy and Connor, 2001] only includes silhouette stimuli, and the data of [Nandy et al., 2013] only includes outline, this equivalence is assumed. Future studies should measure neuron responses to both outline and silhouette stimuli in one set of consistent observations to substantiate or revise this hypothesis. But for this modeling study the following was taken as an assumption: *The response of V2 L4 units to a shape should be as similar as possible when the shape is presented as either an outline or a silhouette*. The concept of this equivalence and the resulting error component of the optimization target is illustrated in Figure 23.

**Fig. 23.**
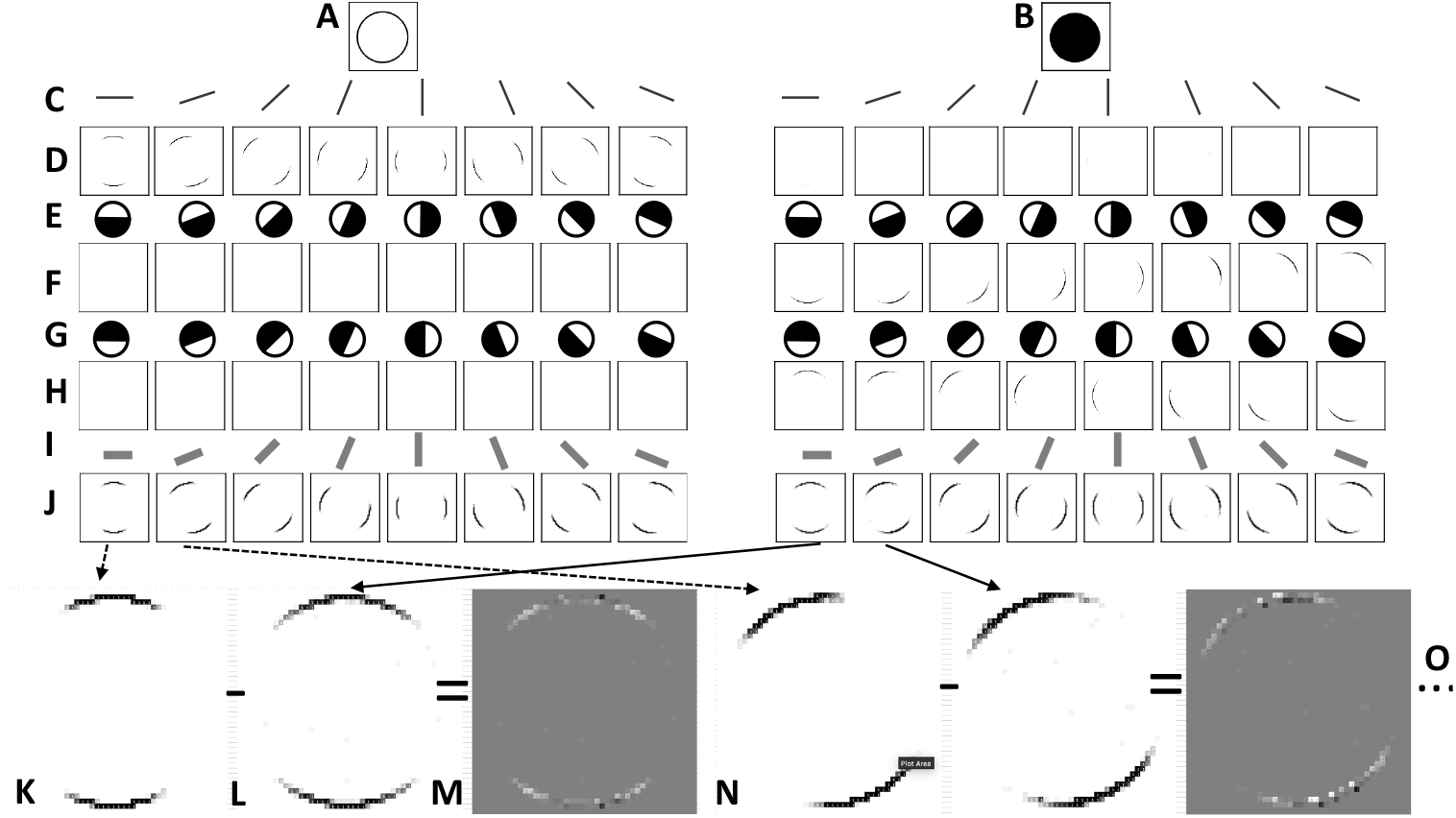
Response to Outline and Silhouette Circle from V1 to L4 V2 **A** Circle outline stimuli - resulting unit activations are on the Left side of the figure. **B** Circle silhouette stimuli - resulting unit activations are on the Right side of the Figure. **C** Pictograms representing each orientation of V1 stripe selective neuron by preferred stimuli. **D** Responses of the V1 *stripe* selective neurons - they respond well to the outline circle, and not at all to the silhouette circle. **E, G** Pictograms representing each orientation of V1 edge selective neuron. **F**,**H** Responses of the V1 *edge* selective neurons - they respond well to the silhouette circle, and hardly at all to the outlined circle. **I** Pictograms representing each orientation of V2 L4 pool unit; see Section 4.4.5 for details. **J** The response of the V2 L4 pool units - the responses for the outline circle and silhouette circle are similar but not exactly the same. **K** Zoom in on the response of the horizontal orientation selective V2 L4 pool unit to the outline circle. **L** Zoom on the response of the horizontal orientation selective V2 L4 pool unit to the silhouette circle. **M** Difference between the V2 L4 pool unit responses to the outline and silhouette circle. The mean square error is used to optimize the model weights. **N** Details of the difference between outline and silhouette responses for the V2 L4 unit with 22.5^°^ orientation preference. **O** Response differences are calculated for the remaining 6 orientation preferences.

On each iteration of the tuning algorithm, the model was presented with a data sample consisting of 20 examples of the circle outline (Figure 23A) and 20 examples of circle silhouette Figure 23B). The average response of each of the V2 L4 unit channels (Figure 23J) over the 20 presentation was then calculated for the two stimuli resulting in two matrices per orientation of the V2 L4 unit. Mathematically, use 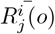 and 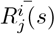 to denote the average responses of the *i*^*th*^ orientation preference and the *j*^*th*^ position in the receptive field to outline (*o*) and silhouette (*s*) stimuli. The square root of the mean of the squared differences, divided by the overall average response, was used as an optimization target to be minimized. That is,

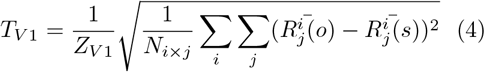

where *T*_*V* 1_ is the optimization target to be minimized and

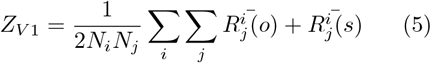

Division by the average response *Z*_*V* 1_ was used to normalize the target and make it more interpretable as this optimization target was further combined with an optimization target function for the V2 units as described in Section 4.5.3.

#### 4.5.3 Tuning V1 and V2 Corner Selective Units

The 12 parameters for the V1 model weights were tuned simultaneously with the 4 parameters for the weights of the most acute interior angle V2 L2/3 unit, having 45^°^ interior angle. The reason for this combined optimization is that the response to acute angles in V2 depends on the interaction of the responses of both the edge and line selective units in V1. This is illustrated in Figure 24: For the acute corner, the arms of the corner trigger a robust response in the V1 stripe selective units but the vertex does not (Figure 24I). Instead, the vertex of the corner triggers a response in the edge selective units(Figure 24D, column 6 and F column 4). This is because at the scale of the V1 units the vertex of the corner appears like a solid region, not a stripe. The different types of V1 units of similar orientation are then pooled in the V2 L4 unit (Figure 24J) providing a continuous view of the orientations in the image for the V2 L2/3 unit selective for a corner. As a result, correct activation of the V2 corner selective unit will require appropriate optimization of the V1 units, and the V1 units should not be optimized without ensuring correct V2 behavior in acute angle units.

**Fig. 24.**
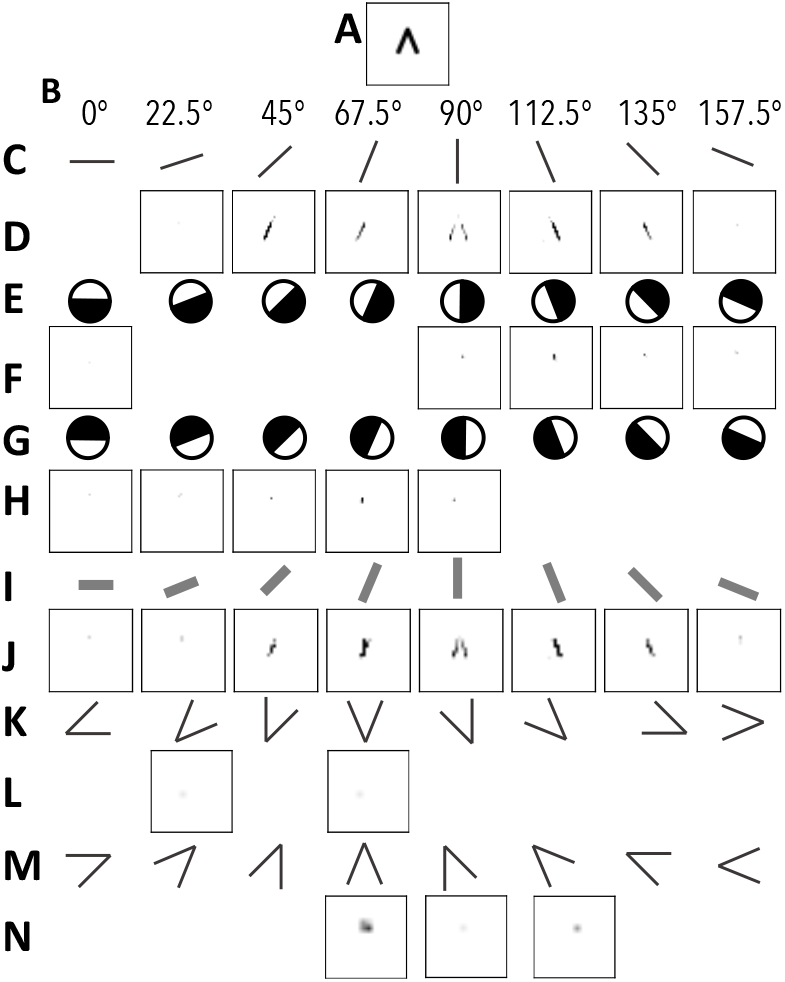
Acute corner response combines V1 Stripe and Edge unit responses.**A** A 45^°^ corner stimuli. **B** Angles of the preferred stimuli in each column. **C** Pictograms representing each orientation of V1 stripe selective neuron by preferred stimuli. **D** Responses of the V1 *stripe* selective neurons - the appropriate orientation selective units respond to the arms of the corner stimuli, but none respond at the vertex: At the scale of these units, the vertex does not appear as a stripe. **E, G** Pictograms representing each orientation of V1 edge selective neurons. For row G, add 180^°^ to the angles in row B to interpret the unit angles. **F**,**H** Responses of the V1 *edge* selective neurons: The edge selective neuron with preferred angles of 112.5^°^ (row F) and 247.5^°^ (row H) respond to the vertex of the corner. **I** Pictograms representing each orientation of V2 L4 pool unit; see Section 4.4.5 for details. The pool unit integrates the response of the edge and stripe selective units into a continuous response at orientations of 67.5^°^ and 112.5^°^. **K, M** Pictograms representing the preferred stimuli of V2 L2/3 units selective to 45^°^ corners. **L**,**M** Response of units selective to 45^°^ corners.

To achieve these goals the overall optimization target for the V2 45^°^ interior angle corner selective units was:

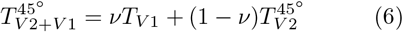

where *T*_*V* 1_ is the V1 optimization target of equation 4 and 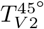 is the V2 optimization target of equation 3. *ν* is a parameter controlling the importance to give to maintaining the balance of the V1 units. After some experimentation *ν* was set to 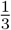 (0.33) for the experiments presented in this report and these optimizations were run for a total of 300 steps.

An illustration of the parameter tuning that resulted in the weights of Tables 8 and 9 is shown in Figure B6. The optimization is run for 300 steps and the errors *T*_*V* 1_ (equation 4) and 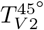 (equation 3) are reduced to around 0.35 and 0.15 respectively, with the final error 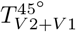 around 0.2. The initial values for the weight params are highly variable, but by the final 50-75 steps they have largely converged.

Figure B6D shows the tuning of the parameters *ρ*_*E*_ which controls the excitatory vs. inhibitory input weight balance for each variant of the 45^°^ interior angle corner selective unit. This parameter was allowed to vary in the optimization and eventually settled on values below 0.5. As described in Section 4.4.2, *ρ*_*E*_ for the V1 units was fixed at 0.4. After simultaneous tuning of the V1 unit weights and the weights of the 45^°^ interior angle V2 corner selective unit, the V1 weights are fixed and the remaining V2 unit weights were optimized in parallel. The target for this optimization was Equation 3 alone, and as there were fewer parameters to optimize these optimizations were run for only 200 steps. The optimization that resulted in the weights of Table 9 is shown in Figure B5 for the 90^°^ interior angle V2 units. As can be seen in Figure B5A, the error starts out high for firing in response to the preferred orientation 90^°^ stimuli 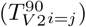,and low for the response to non-preferred units 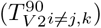- this indicates the weights are generally low and the unit did not fire enough. Figure B5B shows that *w*_*E*_ is generally increased throughout the optimization, resulting in a dramatic reduction in error for the preferred stimuli with only a modest increase in error for the response to non-preferred stimuli. The optimization settles on lower proportion of excitation *ρ*_*E*_ for the 22.5^°^ angle variant (Figure B5C.)

#### 4.5.2 Tuning V2 Ultralong Units

For the V2 ultralong units the optimization goal is slightly different from those with an interior angle less than 180^°^ . The units need to have a broad tuning with a spike rate that decays with the angular distance of the stimuli to the preferred orientations - this is a requirement for the V4 low curvature selective unit to match the recordings of [Nandy et al., 2013]. Instead of using the maximum firing rate to the stimuli, like the V2 units selective to corners, the optimization target rate uses the mean firing rate, 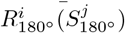,where *I* indexes the preferred orientation and *j* indexes the presented stimuli orientation. The optimization target takes the form:

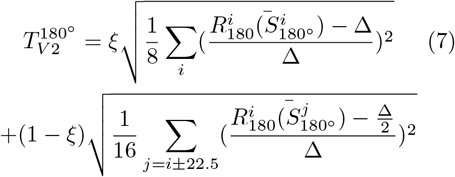

where Δ denotes the target average firing rate to the preferred stimuli. Division by Δ in Equation 7 is used as a normalization constant to make the error more interpretable as a percentage of the target rate. In words, the target is the mean square percentage error of the difference between the target average and the average response to the preferred stimuli weighted by a parameter *ξ*, plus the mean square error between the target average divided by two for the stimuli that are 22.5^°^ away from preferred, as a percentage of the preferred orientation target, with the non-preferred orientation RMSE weighted by one minus *ξ*. In principle there can be other terms in Equation 7 to penalize errors for responses to orientation greater than 22.5^°^ from the preferred orientation. But in practice the ultralong units had zero response to the stimuli more than 22.5^°^ from their preferred orientation anyway, so those extra terms can be ignored. (See Figure A1 row C, columns 9- 16 for illustration of ultralong responses to the 0^°^ stimulus.)

To match the response to stimuli in V4 there is not one specific mean firing rate Δ that is required for the ultralong V2 unit but rather a range of values that will make the simulation reproduce the recording. This is because the sensitivity to the output can be controlled by the input weights of the efferent V4 units. So given a target Δ, an allowable band was set as a range around it. Given an allowable deviation from the target, *η*, on each iteration of the algorithm the target was re-calculated as

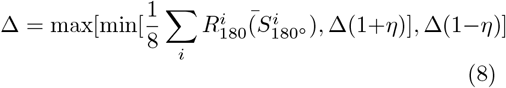

That is, the average response becomes the new target as long as the average response is within the range [Δ(1− η), Δ(1 + *η*)]. Otherwise the target is clipped to the min/max as appropriate. After some preliminary simulations it was found that a reasonable target average firing rate was around 0.0075, which in the 28 *×* 28 channel size for the V4 L4 units corresponds to approximately 6 out of 784 units firing in response to the preferred stimuli. The parameter *η* was set to 0.1, so the precise average could vary by up to 10%. The parameter *ξ* was set to 0.66, so most of the optimization weight went to matching the response to the preferred stimuli. The optimization was run for 300 steps and the simulation resulting in the weights of Table 9 is shown in Figure B7.

Figure B7 shows that the error declines from about 0.4 to 0.1 over the 300 step optimization, with generally equivalent contributions from matching the preferred and non-preferred targets (See Figure 4.5.4A). The weights are tuned to values between 0.1 and 0.25 with the highest values going to those weights on the 22.5^°^ variant afferent inputs (See Figure 4.5.4B-D). The excitatory/inhibitory balance parameters *ρ*_*E*_ fluctuated but in the end converged to similar values around 0.45 for all variants. (See Figure 4.5.4E). Figure 4.5.4F shows the final tuned average firing rates: the firing rate to the prefrred stimuli average 0.0065 although there is some variability by preferred orientation; the firing rates to the less preferred stimuli at ± 22.5^°^ from the preferred stimuli average around 50% of the magnitude of the preferred response but also vary by orientation.

#### 4.5.5 Tuning of the V4 Units

After all of the V2 unit weights were fixed by optimization, the 6 weight parameters *w*_*E*_ and 6 excitatory input ratio parameters *ρ*_*E*_ were by manual experimentation. Setting these parameters by manual experimentation was necessary because the data from V4 recordings is not available for calculation of optimization target. Also, tuning these parameters manually was rather simple given suitably balanced (optimized) inputs from the V2 units and the fact that the model is by construction suitable to reproducing the response to the stimuli. After the final V2 parameters were set, it took from 2-4 iterations of manual tuning to set the final weights for the V2 units.

### Input Stimuli

#### 4.6.1 Concave and Convex Boundary Stimuli

The boundary combination stimuli were created from the vertex data and sample code provided at [Pasupathy, 2001]. The code was ported to Python and the shapes were drawn onto a 112*×* 112 pixel image. The shapes were drawn as filled silhouettes to simulate the experiments of [Pasupathy and Connor, 2001] and were drawn as outlines with a line width of 1.5 for the experiments described in Section 2.6.3 and Section 2.6.4.

To create the concentric and rotated pair stimuli for the experiments descried in Section 2.6.4 the original set of 366 stimuli was reduced to 324 by removing 7 out of 50 configurations which were either too narrow to fit a concentric version, or the shape was symmetric or nearly symmetric to rotation. The omitted shapes correspond to vertex set numbers 0, 1, 7, 11, 12, 13 and 14 in the vertex set provided at [Pasupathy, 2001] (These are the vertex number indices *omitted* in Table C1.) Outlines of the boundary were created as described above using 1.5 pixel line width. To form the rotated pairs, the initial shape was replaced by two rotated versions +/-*π/*16 apart.

To create the concentric shapes, the initial shape was retained and a second version was created that was reduced in size and translated and/or rotated to place the reduced shape outline at a consistent distance from the original outline.

The procedure for each vertex set was as follows, referring the parameters listed in Table C1:

1. Optionally rotate the shape by ± 0.25 radians for shapes where the shape in [Pasupathy, 2001] was not aligned horizontally or vertically. Horizontal of vertical alignment made it straightforward to set the remaining parameters. These are the vertex sets with the parameter *ψ ≠* 0 in Table C1.
2. Scale the vertex set by *ζ*_*y*_ and *ζ*_*x*_ in the Y and X dimensions respectively.
3. Optionally shift the shape by *δ*_*y*_ and *δ*_*x*_ in the Y and X dimensions respectively
4. Optionally rotate the shape by Ψ radians
5. If the vertex set was rotated by ± 0.25 radians before scaling, rotate it back to its original orientation.

#### 4.6.2 Grid Of Composite Bar Stimuli

The grid of composite bar stimuli images were created following the approach described in [Nandy et al., 2013] and adapted to the neural network modeling framework. Line segments 12 pixels long were drawn in black on a white 112*×*112 pixel background. Each model stimuli consisted of 3 line segments joined at the end in which the arms had half the length of central bar. These arrangements were formed into 5 different degrees of curvature using 5 different angles: 0^°^ for straight lines, 22.5^°^ for low curvature, 45^°^ for moderate curvature, 77.5^°^ for high curvature, and 90^°^ for “C” shaped. Each stimuli was used in 16 rotated versions separated by angles of 22.5^°^. The stimuli were placed in the image on a 3*×* 3 grid, 38 pixels apart, beginning at pixel (18,18). This created a total of 5*×* 16 *×* 9 = 720 stimuli. Note that as in [Nandy et al., 2013], the straight line stimuli are redundant when rotated by 180^°^ but these duplicates were left in the stimuli set for simplicity. As described in Section 4.5, corner stimuli like the one illustrated in Figure A2 were generated to tune the parameters of the V1 and V2 models. These stimuli were created using the same techniques as the composite bar stimuli that recreated the stimuli of [Nandy et al., 2013]. The corner stimuli used for tuning consisted of just two bars having equal length, rather than 3 bars in which the center bar has twice the length of the two arms. The corner stimuli were generated with interior angles ranging from 22.5^°^ to 157.5^°^ in steps of 22.5^°^ . The tuning stimuli set consisted of only a single placement of each stimuli in the center of the model receptive field, rather than repeating the placement on a 3 *×* 3 grid.

### 4.7 Hardware and Runtime

Assuming atemporal binary inputs and outputs as in Listing 1, the tensor graph for 6 layers (3 cortical areas, 2 layers per area) has around 150 operations. This is more than suggested by Listing 1, as the actual model code must handle convolution filters of varied sizes and the input layer. The spike time simulation using the PSP equations given by Listing 2, 3 and Listing 5 was run for 30 ms discretized at 0.3 ms intervals; the resulting tensor simulation is an approximately 60,000-operation graph representing the layers at each time, plus additional operations for the PSP calculations and logic relating to the temporal simulation. (TODO: Recheck that final number.) These large tensor graphs run on GPU accelerated hardware: a simulation of a 900,000-neuron network for 30 ms takes 2.5 hours to run on a MacBook Pro with an M3 GPU and 8 minutes on an AWS g6e.xlarge GPU instance having one NVIDIA L40S Tensor Core GPU with 48 GB of GPU Memory. A high amount of GPU memory per GPU is required to process the relatively large tensor graph. As previously mentioned, the GPU memory is the key constraining factor when it comes to running longer or more detailed simulations. At the time of this writing, the Mac M3 GPU experiments resulted in significant numerical deviation from the NVIDIA results and Mac M3 runs were used for debugging only - all results presented here were run on an NVIDIA GPU.

### 4.8 PSP Maximum

The constant *Z* in equation 1 is determined by noting that the difference of exponentials in equation 1 takes its maximum at a time after the input arrival given by:

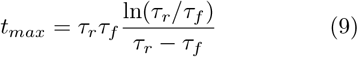

and the constant *Z* is given by

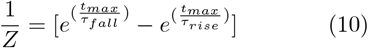

## Acknowledgments

Thanks to John Reynolds and Anirvan Nandy for allowing reproduction of their figures and for helpful discussions and suggestions.

## Code Availability

Upon publication the source code from this study will be available at http://github.com/carl24k; during the review process the source code is available by request.

## Appendix A Supplemental Results

Figure A1, A2 and A3 illustrates the response of the entire network. Figure A1 is the response to a straight line, Figure A2 is the response to a 90^°^ corner, and Figure A3 is the response to a complex shape defined by boundary elements. The sub-plots show the responses of the *L4 pool units* in V2 and V4, and the V4 units themselves. The L4 pool units in V2 serve as a concise summary of the response of V1, as it does not distinguish if an oriented segment in the stimuli is an edge of a silhouette or a line. Also, as the V4 L4 units pool the response of their inputs they are both more reliable and at a zoomed in scale in comparison to their afferents in a previous layer.

The following annotation key serves for all three figures:

- **A** Heatmap of average input intensity. In Figure A1 and Figure A2 the heatmap is zoomed in to the central 42*×* 42 units, out of 116; in Figure A3 the heatmap is zoomed to the central 90 *×* 90 units.
- **B** Pictogram representation of the preferred stimulus of the V2 L4 units, which pool the V1 Stripe and Edge selective units - see Figure 20 for the V1 unit convolution filters.
- **C** Firing rate of the V2 L4 orientation selective units. In Figure A1 and Figure A2 the heatmap is zoomed in to the central 28 *×* 28 units (out of 56); in Figure A3 the heatmap is zoomed to the central 50*×*50 units.
- **D** Pictogram for the preferred stimulus of the ultralong units.
- **E** Firing rate of the V4 L4 ultralong selective pool unit responses. The zoom is as in **C**. See Figure 21 for the convolution filters defining the V2 ultralong units.
- **F** Pictograms for the preferred stimulus of complex-shaped units having 157.5^°^ angle between sub-units. See Figure 22 for the convolution filters defining the V2 units having 157.5^°^ interior angle.
- **G** Firing rate of the V4 L4 157.5^°^ complex-shaped selective pool units. The zoom is as in **C**.
- **H** Pictograms for the preferred stimulus of complex-shaped units having 135^°^ angle between sub-units. See Figure 22 for the convolution filters defining the V2 units having 135^°^ interior angle
- **I** Firing rate of the V4 L4 135^°^ complex-shaped selective pool units. The zoom is as in **C**.
- **J** Pictograms for the preferred stimulus of complex-shaped units having 112.5^°^ angle between sub-units. See Figure 22 for the convolution filters defining the V2 units having 112.5^°^ interior angle
- **K** Firing rate of the V4 L4 112.5^°^ complex-shaped selective pool units. The zoom is as in **C**.
- **L** Pictograms for the preferred stimulus of complex-shaped units having 90^°^ angle between sub-units. See Figure 21 for the convolution filters defining the V2 units having 90^°^ interior angle
- **M** Firing rate of the V4 L4 90^°^ complex-shaped selective pool units. The zoom is as in **C**.
- **N** Pictograms for the preferred stimulus of complex-shaped units having 67.5^°^ angle between sub-units. See Figure 22 for the convolution filters defining the V2 units having 67.5^°^ interior angle
- **O** Firing rate of the V4 L4 67.5^°^ complex-shaped selective pool units. The zoom is as in **C**.
- **P** Pictograms for the preferred stimulus of complex-shaped units having 45^°^ angle between sub-units. See Figure 22 for the convolution filters defining the V2 units having 45^°^ interior angle
- **Q** Firing rate of the V4 L4 45^°^ complex-shaped selective pool units. The zoom is as in **C**.
- **R** Pictogram representation of the preferred stimulus of the V4 units selective to acute convex curvature in a position. These units are defined by the convolution filters illustrated in Figured 4.
- **S** Firing rates of V4 units selective to acute convex curvature in a position.
- **T** Pictogram representation of the preferred stimulus of the V4 units selective to medium concave curvature in a position. These units are defined by the convolution filters illustrated in Figured 8.
- **U** Firing rates of V4 units selective to acute medium concave curvature in a position.
- **V** Pictogram representation of the preferred stimulus of the V4 units selective to medium convex curvature in a position. These units are defined by the convolution filters illustrated in Figured 8.
- **W** Firing rates of V4 units selective to acute medium convex curvature in a position.
- **X** Pictogram representation of the preferred stimulus of the V4 units selective to acute convex curvature throughout the receptive field. These units are defined by the convolution filters illustrated in Figured 6.
- **Y** Firing rates of the V4 units selective to acute convex curvature throughout the receptive field.
- **Z** Pictogram representation of the preferred stimulus of the V4 units selective medium convex curvature throughout the receptive field. These units are defined by the convolution filters illustrated in Figured 10.
- **AA** Firing rates V4 units selective to medium convex curvature throughout the receptive field.
- **BB** Pictogram representation of the preferred stimulus of the V4 low curvature selective unit with 8 different preferred orientations. These units are defined by the convolution filters in Figure 12.
- **CC** Firing rates of the V4 low curvature selective units.

**Fig. A1.**
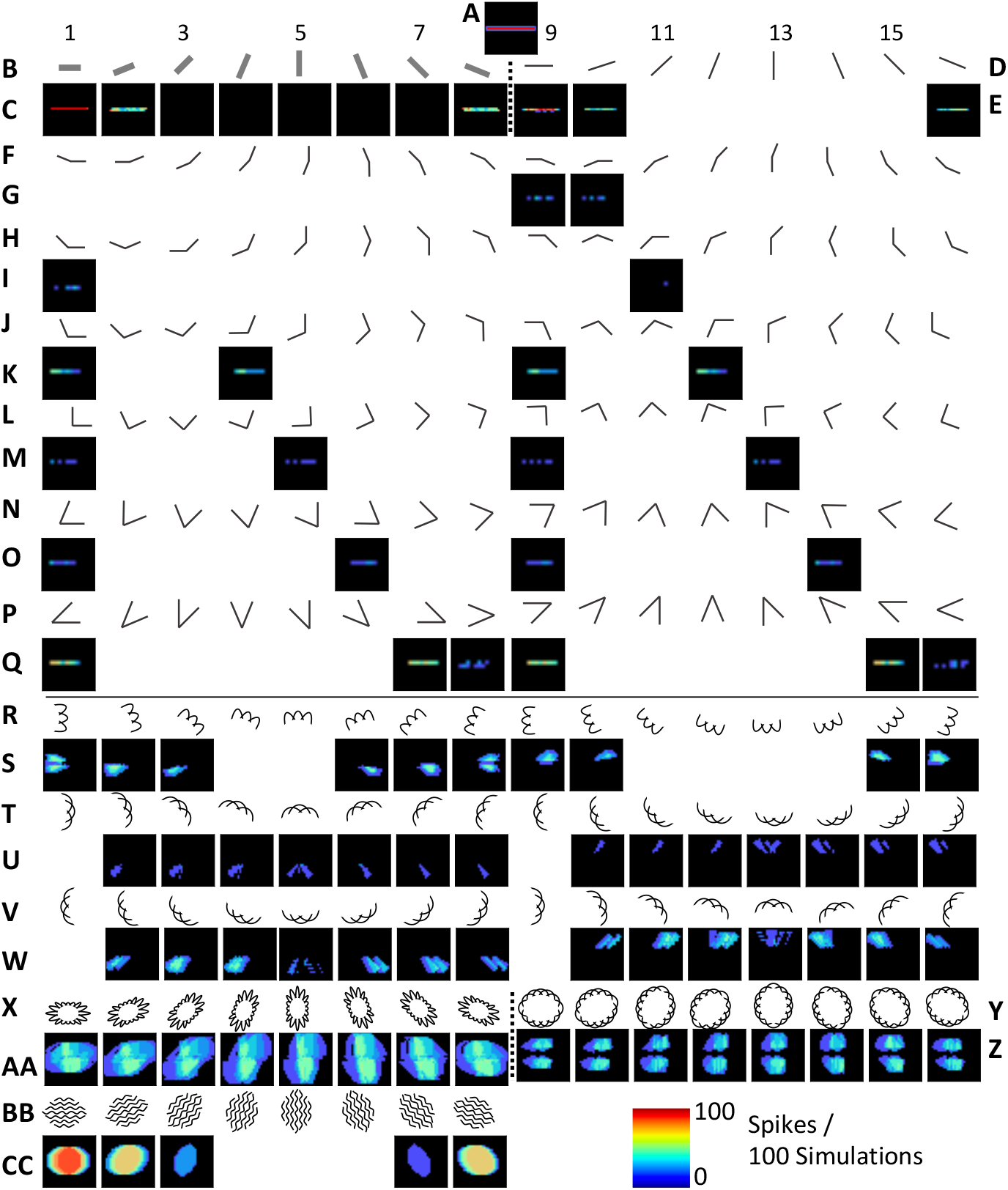
Illustration of Network Response to Low Curvature Stimuli. See text for annotation key.

**Fig. A2.**
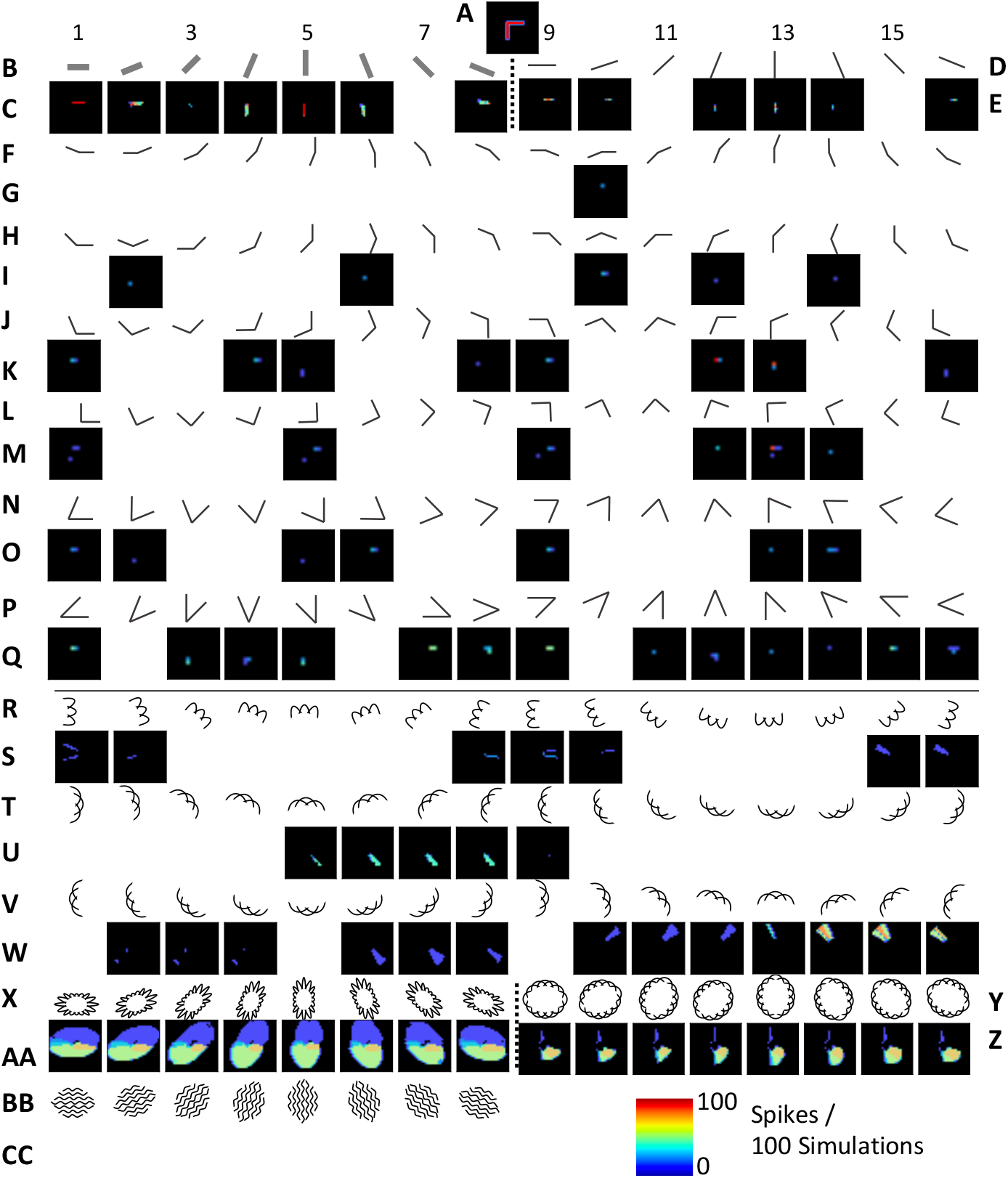
Illustration of Network Response to a Corner Shape Stimuli. See text for annotation key.

**Fig. A3.**
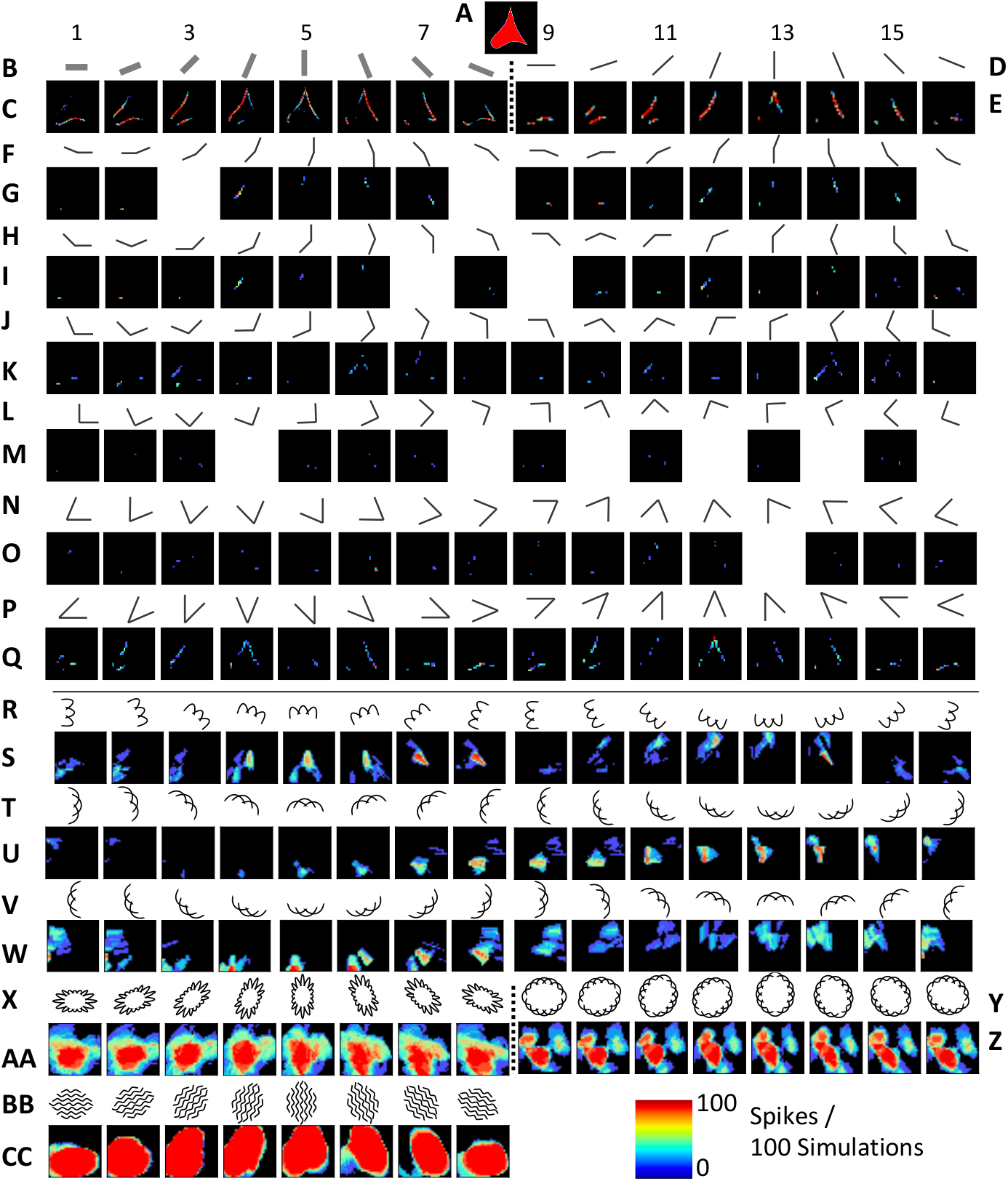
Illustration of Network Response to Complex Shape Stimuli. See text for annotation key.

**Fig. A4.**
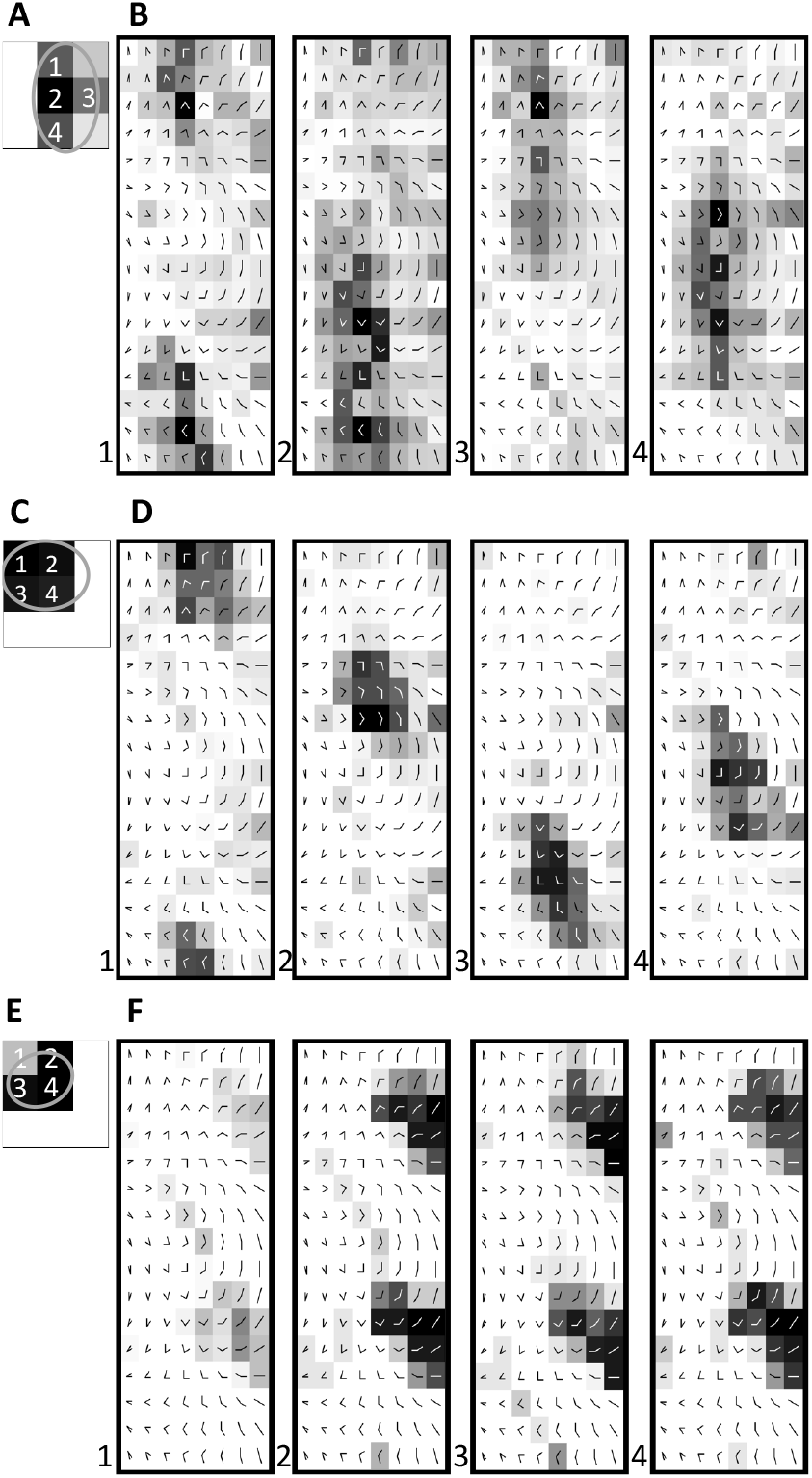
Simulated response of neurons from [Nandy et al., 2013] to stimuli of oriented corners. The stimuli of corners are constructed in a similar fashion, albeit with two bars, as described in the Methods Section 4.6.2. **A** The neuron of Figure 5 and 6 selective for high curvature. **B** The neuron of Figure 9 and Figure 10 selective for medium curvature. **C** The neuron of Figure 11 and Figure 12 selective for low curvature.

## Appendix B Weight Optimization Detail Illustrations

The figures in this section illustrate the process of tuning the convolution filter weights for the V1 and V2 units. Figure B5 illustrates the optimization of the 6 parameters of the V2 90^°^ corner interior angle unit. Figure B6 illustrates the optimization of the 11 parameters of the V1 edge and stripe selective units, and the V2 45^°^ corner interior angle unit. Figure B7 illustrates the optimization of the 10 parameters of the V2 ultralong (180^°^ interior angle) unit. For all of the figures the x-axis represents the iteration of the optimization algorithm. The y-axis of the figures is either the optimization target or the weight parameter indicated by the legend. The optimized parameters are described in the Methods Section 4.4.2, Section 4.4.3. The figures are described in more detail in the Methods Section 4.5.3 and Section 4.5.4.

**Fig. B5.**
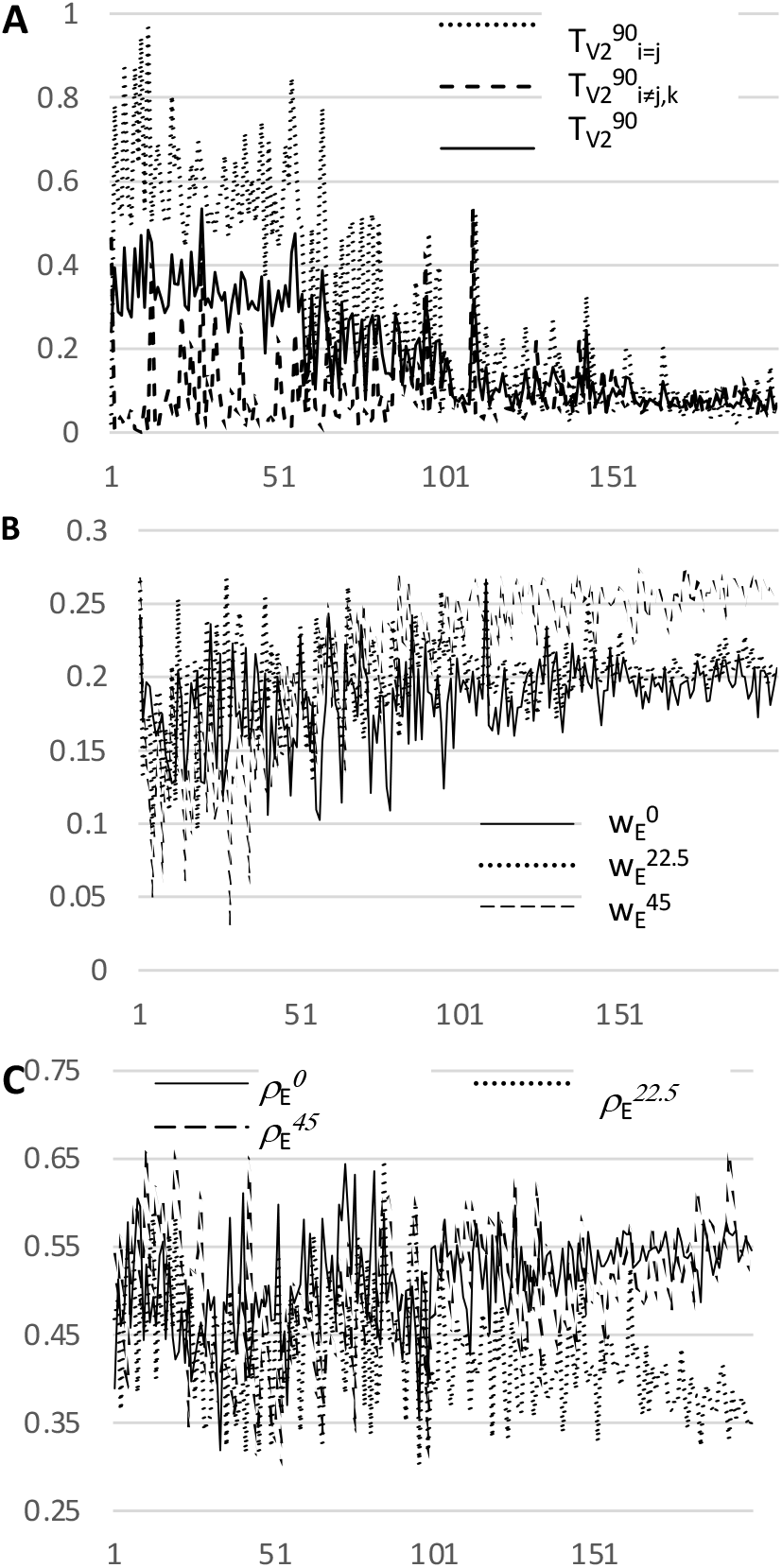
Illustration of Tuning the V2 90^°^ Interior Angle Unit Weights. For details, see section 4.5.3

## Appendix C Concentric Shape Generation Parameters

**Fig. B6.**
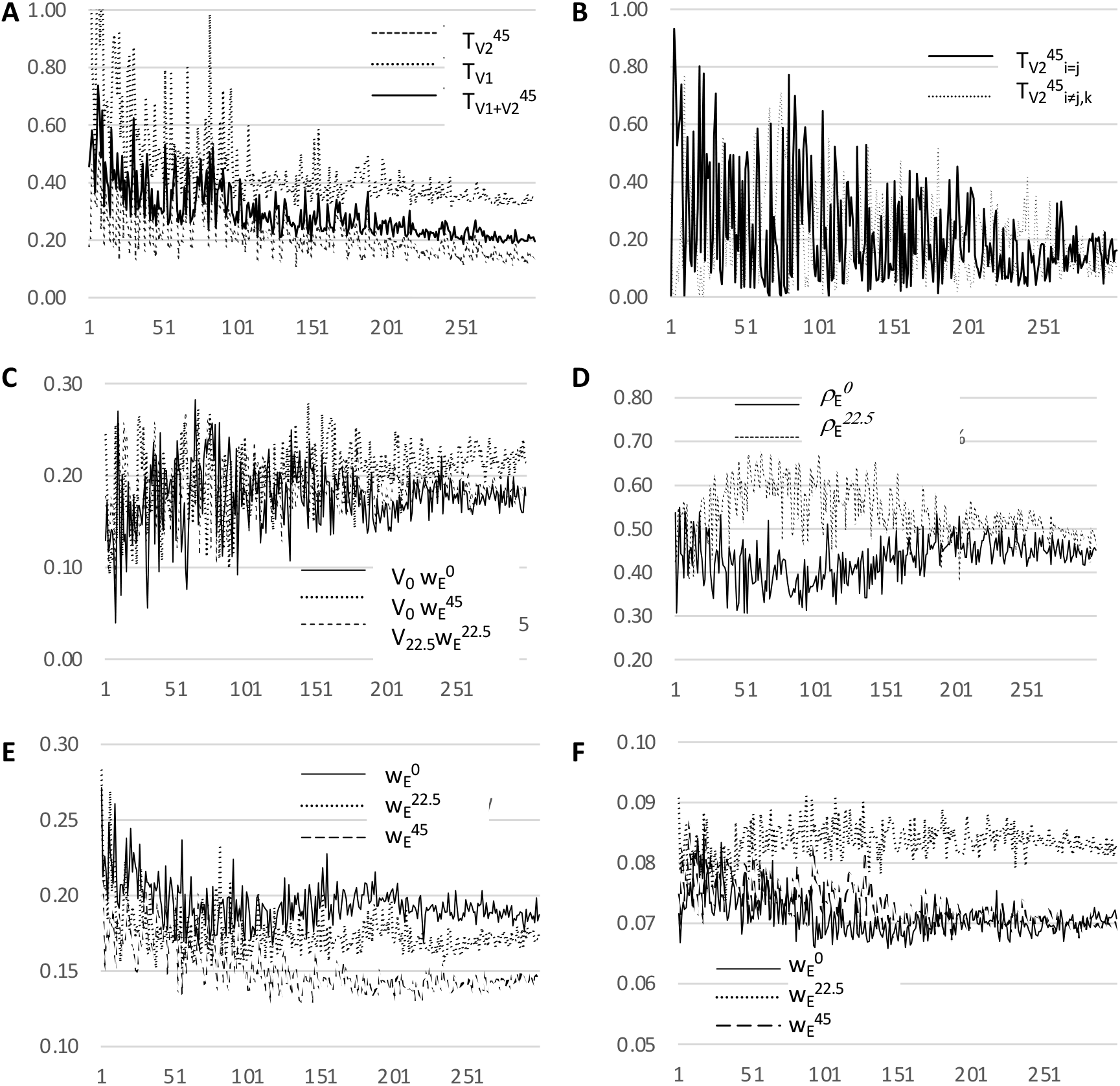
Illustration of Tuning the V2 45^°^ Interior Angle Unit Weights and the V1 Unit Weights. For details, see section 4.5.3

**Fig. B7.**
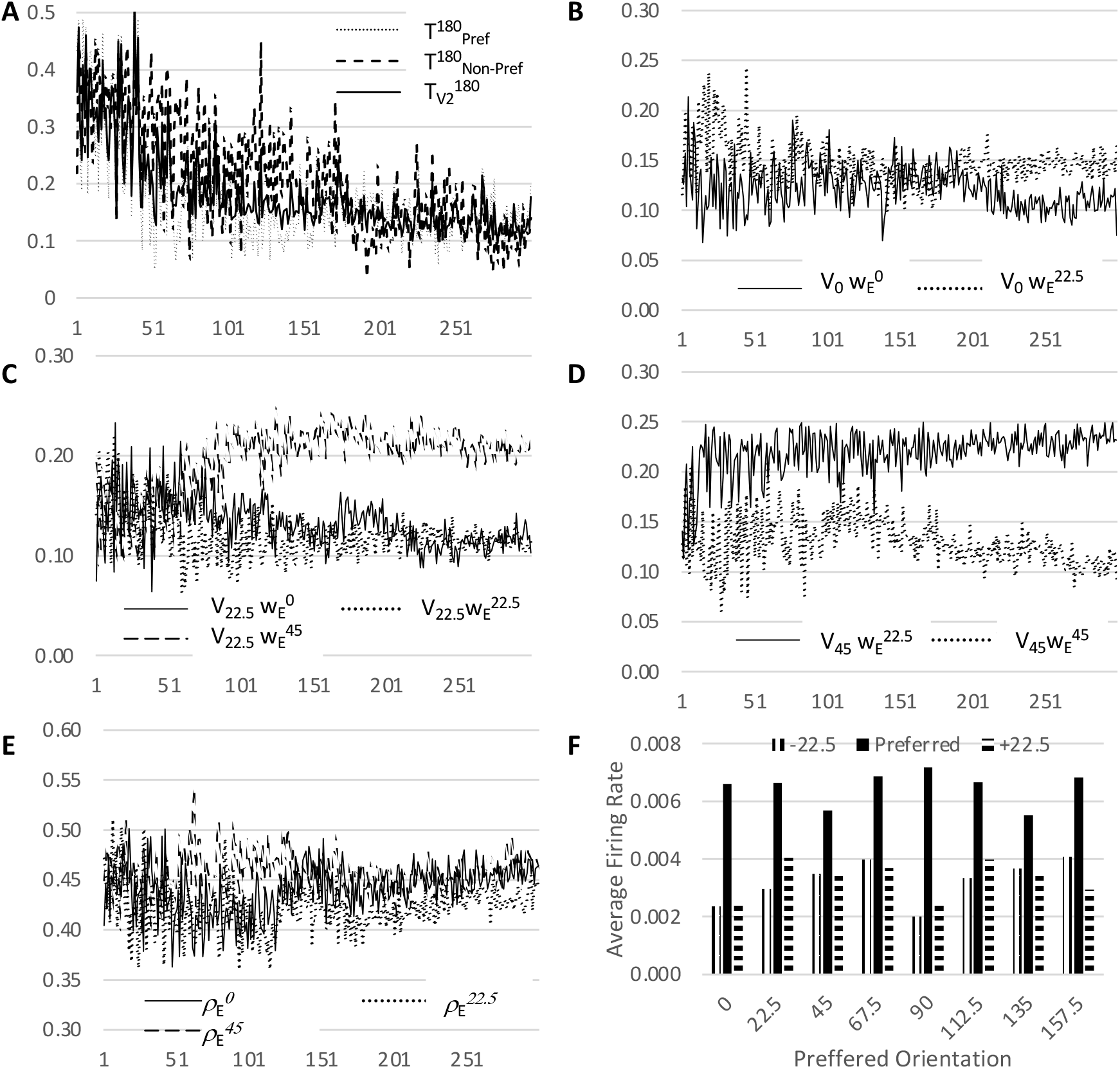
Illustration of Tuning the V2 Ultralong (180^°^ Interior Angle) Unit Weights. For details, see section 4.5.4

**Table C1.** Concentric Boundary Shape Creation Parameters.

| V# | $\zeta_y$ | $\zeta_x$ | $\psi$ | $\delta_y$ | $\delta_x$ | $\Psi$ |
| --- | --- | --- | --- | --- | --- | --- |
| 2 | 0.5 | 0.3 | 0 | 0 | 0 | 0 |
| 3 | 0.5 | 0.3 | 0 | 0 | 0 | 0 |
| 4 | 0.55 | 0.3 | 0 | 0 | 0 | 0 |
| 5 | 0.5 | 0.2 | 0 | 0 | -0.125 | 0 |
| 6 | 0.35 | 0.2 | 0 | 0 | 0 | 0 |
| 8 | 0.4 | 0.2 | -0.25 | -0.12 | 0 | 0 |
| 9 | 0.5 | 0.3 | -0.25 | -0.15 | -0.1 | 0.1 |
| 10 | 0.3 | 0.5 | -0.25 | -0.15 | -0.15 | -0.1 |
| 15 | 0.4 | 0.25 | 0 | -0.12 | 0 | 0 |
| 16 | 0.3 | 0.25 | 0 | -0.13 | 0 | 0 |
| 17 | 0.3 | 0.25 | 0 | -0.09 | 0 | 0 |
| 18 | 0.3 | 0.25 | 0 | -0.09 | 0 | 0 |
| 19 | 0.3 | 0.2 | 0 | -0.15 | 0 | 0 |
| 20 | 0.3 | 0.2 | 0 | -0.15 | 0 | 0 |
| 21 | 0.3 | 0.2 | 0 | -0.1 | 0 | 0 |
| 22 | 0.3 | 0.2 | 0 | -0.12 | 0 | 0 |
| 23 | 0.5 | 0.5 | 0 | 0 | 0 | 0 |
| 24 | 0.4 | 0.25 | 0 | 0 | 0.05 | 0 |
| 25 | 0.3 | 0.45 | 0.25 | 0.025 | 0.025 | -0.05 |
| 26 | 0.45 | 0.3 | 0.25 | -0.025 | -0.025 | 0.05 |
| 27 | 0.2 | 0.4 | 0 | 0 | -0.025 | 0 |
| 28 | 0.4 | 0.2 | 0 | 0.05 | 0.12 | 0 |
| 29 | 0.4 | 0.2 | 0 | -0.05 | 0.12 | 0 |
| 30 | 0.35 | 0.25 | 0 | 0 | 0.05 | 0 |
| 31 | 0.35 | 0.35 | 0 | 0 | 0 | 0 |
| 32 | 0.3 | 0.25 | 0 | 0 | 0.025 | 0 |
| 33 | 0.2 | 0.3 | 0 | 0 | 0 | 0 |
| 34 | 0.3 | 0.3 | -0.25 | 0 | 0.05 | 0 |
| 35 | 0.2 | 0.3 | 0 | 0 | -0.05 | 0 |
| 36 | 0.25 | 0.25 | 0 | 0 | 0 | 0 |
| 37 | 0.25 | 0.2 | 0 | 0 | 0.15 | -0.1 |
| 38 | 0.2 | 0.25 | 0 | 0 | -0.2 | 0.1 |
| 39 | 0.2 | 0.35 | 0 | 0 | -0.25 | 0 |
| 40 | 0.25 | 0.2 | 0 | -0.05 | 0.15 | -0.05 |
| 41 | 0.25 | 0.2 | 0 | -0.05 | -0.15 | 0.05 |
| 42 | 0.4 | 0.2 | 0 | 0 | -0.1 | 0 |
| 43 | 0.2 | 0.25 | -0.25 | -0.1 | 0 | 0 |
| 44 | 0.2 | 0.3 | -0.25 | 0 | 0 | 0 |
| 45 | 0.2 | 0.2 | -0.25 | 0 | -0.125 | 0 |
| 46 | 0.2 | 0.2 | 0 | 0 | -0.15 | 0 |
| 47 | 0.3 | 0.3 | 0 | -0.1 | 0.05 | 0 |
| 48 | 0.3 | 0.3 | 0 | 0.1 | 0.05 | 0 |
| 49 | 0.2 | 0.3 | 0 | 0 | 0.2 | 0 |
| 50 | 0.3 | 0.2 | -0.25 | 0 | -0.1 | 0 |

## Footnotes

1 Another element necessary for fully realistic spiking computation absent from the model of single cascades is a mechanism of synchronization that enables rapid, successive cascade computations. As will be described in detail in the Results and Methods (Section 2.4 and 4.1), the decay time constants of PSP’s at the soma are long relative to the time scale of spike integration. It is hypothesized that synchronized inhibition associated with inhibitory networks responsible for the gamma oscillation [Buzsáki and Wang, 2012] act as a “reset” mechanism for repolarization faster than passive decay implied by the time constants. Modeling such mechanisms are also beyond the scope of the present study.

## References

[Akiba et al., 2019] Akiba, T., Sano, S., Yanase, T., Ohta, T., and Koyama, M. (2019). Optuna: A next-generation hyperparameter optimization framework. In Proceedings of the 25th ACM SIGKDD International Conference on Knowledge Discovery and Data Mining.

[Bartunov et al., 2018] Bartunov, S., Santoro, A., Richards, B. A., Marris, L., Hinton, G. E., and Lillicrap, T. (2018). Assessing the scalability of biologically-motivated deep learning algorithms and architectures. arXiv preprint 1807.04587.

[Behrens and Sporns, 2012] Behrens, T. E. and Sporns, O. (2012). Human connectomics. Current opinion in neurobiology, 22(1):144–153.

[Bengio et al., 2015] Bengio, Y., Lee, D.-H., Bornschein, J., Mesnard, T., and Lin, Z. (2015). Towards biologically plausible deep learning. arXiv preprint 1502.04156.

[Bengio et al., 2017] Bengio, Y., Mesnard, T., Fischer, A., Zhang, S., and Wu, Y. (2017). Stdp-compatible approximation of backpropagation in an energy-based model. Neural computation, 29(3):555–577.

[Borst and Helmstaedter, 2015] Borst, A. and Helmstaedter, M. (2015). Common circuit design in fly and mammalian motion vision. Nature neuroscience, 18(8):1067–1076.

[Buzsáki and Wang, 2012] Buzsáki, G. and Wang, X.-J. (2012). Mechanisms of gamma oscillations. Annual review of neuroscience, 35(1):203–225.

[Cadena et al., 2019] Cadena, S. A., Denfield, G. H., Walker, E. Y., Gatys, L. A., Tolias, A. S., Bethge, M., and Ecker, A. S. (2019). Deep convolutional models improve predictions of macaque v1 responses to natural images. PLoS computational biology, 15(4):e1006897.

[Cadieu et al., 2007] Cadieu, C., Kouh, M., Pasupathy, A., Connor, C. E., Riesenhuber, M., and Poggio, T. (2007). A model of v4 shape selectivity and invariance. Journal of neurophysiology, 98(3):1733–1750.

[Carandini et al., 2005] Carandini, M., Demb, J. B., Mante, V., Tolhurst, D. J., Dan, Y., Olshausen, B. A., Gallant, J. L., and Rust, N. C. (2005). Do we know what the early visual system does? Journal of Neuroscience, 25(46):10577–10597.

[Cornford et al., 2020] Cornford, J., Kalajdzievski, D., Leite, M., Lamarquette, A., Kullmann, D. M., and Richards, B. A. (2020). Learning to live with dale’s principle: Anns with separate excitatory and inhibitory units. In International Conference on Learning Representations.

[Dale, 1935] Dale, H. (1935). Pharmacology and nerve-endings.

[Dobbins et al., 1987] Dobbins, A., Zucker, S. W., and Cynader, M. S. (1987). Endstopped neurons in the visual cortex as a substrate for calculating curvature. Nature, 329(6138):438– 441.

[Douglas and Martin, 2004] Douglas, R. J. and Martin, K. A. (2004). Neuronal circuits of the neocortex. Annu. Rev. Neurosci., 27:419–451.

[El-Shamayleh and Pasupathy, 2016] El-Shamayleh, Y. and Pasupathy, A. (2016). Contour curvature as an invariant code for objects in visual area v4. Journal of Neuroscience, 36(20):5532–5543.

[Feather et al., 2023] Feather, J., Leclerc, G., Mkadry, A., and McDermott, J. H. (2023). Model metamers reveal divergent invariances between biological and artificial neural networks. Nature Neuroscience, 26(11):2017–2034.

[Ferster and Miller, 2000] Ferster, D. and Miller, K. D. (2000). Neural mechanisms of orientation selectivity in the visual cortex. Annual review of neuroscience, 23(1):441–471.

[Fukushima, 1980] Fukushima, K. (1980). Neocognitron: A self-organizing neural network model for a mechanism of pattern recognition unaffected by shift in position. Biological cybernetics, 36(4):193–202.

[Gerstner and Kistler, 2002] Gerstner, W. and Kistler, W. M. (2002). Spiking neuron models: Single neurons, populations, plasticity. Cambridge university press.

[Girard et al., 2001] Girard, P., Hupè, J., and Bullier, J. (2001). Feedforward and feedback connections between areas v1 and v2 of the monkey have similar rapid conduction velocities. Journal of neurophysiology, 85(3):1328– 1331.

[Glorot et al., 2011] Glorot, X., Bordes, A., and Bengio, Y. (2011). Deep sparse rectifier neural networks. In Proceedings of the fourteenth international conference on artificial intelligence and statistics, pages 315–323. JMLR Workshop and Conference Proceedings.

[Hansen, 2006] Hansen, N. (2006). The cma evolution strategy: a comparing review. Towards a new evolutionary computation: Advances in the estimation of distribution algorithms, pages 75–102.

[Hoffmann et al., 2015] Hoffmann, J. H., Meyer, H.-S., Schmitt, A. C., Straehle, J., Weitbrecht, T., Sakmann, B., and Helmstaedter, M. (2015). Synaptic conductance estimates of the connection between local inhibitor interneurons and pyramidal neurons in layer 2/3 of a cortical column. Cerebral cortex, 25(11):4415–4429.

[Holmgren et al., 2003] Holmgren, C., Harkany, T., Svennenfors, B., and Zilberter, Y. (2003). Pyramidal cell communication within local networks in layer 2/3 of rat neocortex. The Journal of physiology, 551(1):139–153.

[Hu et al., 2020] Hu, J. M., Song, X. M., Wang, Q., and Roe, A. W. (2020). Curvature domains in v4 of macaque monkey. Elife, 9:e57261.

[Hubel and Wiesel, 1962] Hubel, D. H. and Wiesel, T. N. (1962). Receptive fields, binocular interaction and functional architecture in the cat’s visual cortex. The Journal of physiology, 160(1):106–154.

[Hubel and Wiesel, 1965] Hubel, D. H. and Wiesel, T. N. (1965). Receptive fields and functional architecture in two nonstriate visual areas (18 and 19) of the cat. Journal of neurophysiology.

[Jacob et al., 2021] Jacob, G., Pramod, R., Katti, H., and Arun, S. (2021). Qualitative similarities and differences in visual object representations between brains and deep networks. Nature communications, 12(1):1–14.

[Kheradpisheh et al., 2018] Kheradpisheh, S. R., Ganjtabesh, M., Thorpe, S. J., and Masquelier, T. (2018). Stdp-based spiking deep convolutional neural networks for object recognition. Neural Networks, 99:56–67.

[Kietzmann et al., 2019] Kietzmann, T. C., McClure, P., and Kriegeskorte, N. (2019). Deep neural networks in computational neuroscience. In Oxford research encyclopedia of neuroscience.

[Kim et al., 2014] Kim, J. S., Greene, M. J., Zlateski, A., Lee, K., Richardson, M., Turaga, S. C., Purcaro, M., Balkam, M., Robinson, A., Behabadi, B. F., Campos, M., Denk, W., Seung, S. H., and the EyeWireres (2014). Space-time wiring specificity supports direction selectivity in the retina. Nature, 509(7500):331.

[Kim et al., 2019] Kim, T., Bair, W., and Pasupathy, A. (2019). Neural coding for shape and texture in macaque area v4. Journal of Neuroscience, 39(24):4760–4774.

[Kisvárday et al., 1997] Kisvárday, Z. F., Toth, E., Rausch, M., and Eysel, U. T. (1997). Orientation-specific relationship between populations of excitatory and inhibitory lateral connections in the visual cortex of the cat. Cerebral Cortex (New York, NY: 1991), 7(7):605–618.

[Ko et al., 2013] Ko, H., Cossell, L., Baragli, C., Antolik, J., Clopath, C., Hofer, S. B., and Mrsic-Flogel, T. D. (2013). The emergence of functional microcircuits in visual cortex. Nature, 496(7443):96–100.

[Kobatake and Tanaka, 1994] Kobatake, E. and Tanaka, K. (1994). Neuronal selectivities to complex object features in the ventral visual pathway of the macaque cerebral cortex. Journal of neurophysiology, 71(3):856–867.

[LeCun et al., 2015] LeCun, Y., Bengio, Y., and Hinton, G. (2015). Deep learning. nature, 521(7553):436–444.

[LeCun et al., 1998] LeCun, Y., Bottou, L., Bengio, Y., and Haffner, P. (1998). Gradient-based learning applied to document recognition. Proceedings of the IEEE, 86(11):2278–2324.

[Liang et al., 2017] Liang, H., Gong, X., Chen, M., Yan, Y., Li, W., and Gilbert, C. D. (2017). Interactions between feedback and lateral connections in the primary visual cortex. Proceedings of the National Academy of Sciences, 114(32):8637–8642.

[Liu et al., 2016] Liu, L., She, L., Chen, M., Liu, T., Lu, H. D., Dan, Y., and Poo, M.-m. (2016). Spatial structure of neuronal receptive field in awake monkey secondary visual cortex (v2). Proceedings of the National Academy of Sciences, 113(7):1913–1918.

[McCulloch and Pitts, 1943] McCulloch, W. S. and Pitts, W. (1943). A logical calculus of the ideas immanent in nervous activity. The bulletin of mathematical biophysics, 5(4):115–133.

[Miller, 1994] Miller, K. D. (1994). A model for the development of simple cell receptive fields and the ordered arrangement of orientation columns through activity-dependent competition between on-and off-center inputs. Journal of Neuroscience, 14(1):409–441.

[Mueller, 1999] Mueller, B. K. (1999). Growth cone guidance: first steps towards a deeper understanding. Annual review of neuroscience, 22(1):351–388.

[Nandy et al., 2013] Nandy, A. S., Sharpee, T. O., Reynolds, J. H., and Mitchell, J. F. (2013). The fine structure of shape tuning in area v4. Neuron, 78(6):1102–1115.

[Pasupathy, 2001] Pasupathy, A. (2001). ShapeLAB Pasupathy and Connor 2001 Shapes, https://depts.washington.edu/shapelab/resources/p-c-2001.

[Pasupathy and Connor, 2001] Pasupathy, A. and Connor, C. E. (2001). Shape representation in area v4: position-specific tuning for boundary conformation. Journal of neurophysiology, 86(5):2505–2519.

[Perrett and Oram, 1993] Perrett, D. I. and Oram, M. W. (1993). Neurophysiology of shape processing. Image and Vision Computing, 11(6):317–333.

[Plomp et al., 2017] Plomp, G., Michel, C. M., and Quairiaux, C. (2017). Systematic population spike delays across cortical layers within and between primary sensory areas. Scientific reports, 7(1):1–14.

[Ponce et al., 2017] Ponce, C. R., Hartmann, T. S., and Livingstone, M. S. (2017). Endstopping predicts curvature tuning along the ventral stream. Journal of neuroscience, 37(3):648–659.

[Pospisil et al., 2018] Pospisil, D. A., Pasupathy, A., and Bair, W. (2018). ‘artiphysiology’reveals v4-like shape tuning in a deep network trained for image classification. Elife, 7:e38242.

[Price and Born, 2009] Price, N. S. and Born, R. (2009). Representation of movement. Encyclopedia of neuroscience, 8:107–114.

[Riesenhuber and Poggio, 1999] Riesenhuber, M. and Poggio, T. (1999). Hierarchical models of object recognition in cortex. Nature neuroscience, 2(11):1019–1025.

[Roe et al., 2012] Roe, A. W., Chelazzi, L., Connor, C. E., Conway, B. R., Fujita, I., Gallant, J. L., Lu, H., and Vanduffel, W. (2012). Toward a unified theory of visual area v4. Neuron, 74(1):12–29.

[Rumelhart et al., 1986] Rumelhart, D. E., Hinton, G. E., and Williams, R. J. (1986). Learning representations by back-propagating errors. nature, 323(6088):533–536.

[Seeman et al., 2018] Seeman, S. C., Campagnola, L., Davoudian, P. A., Hoggarth, A., Hage, T. A., Bosma-Moody, A., Baker, C. A., Lee, J. H., Mihalas, S., Teeter, C., et al. (2018). Sparse recurrent excitatory connectivity in the microcircuit of the adult mouse and human cortex. elife, 7:e37349.

[Serre, 2019] Serre, T. (2019). Deep learning: the good, the bad, and the ugly. Annual review of vision science, 5(1):399–426.

[Shirasaki and Pfaff, 2002] Shirasaki, R. and Pfaff, S. L. (2002). Transcriptional codes and the control of neuronal identity. Annual review of neuroscience, 25(1):251–281.

[Shuvaev et al., 2024] Shuvaev, S., Lachi, D., Koulakov, A., and Zador, A. (2024). Encoding innate ability through a genomic bottleneck. Proceedings of the National Academy of Sciences, 121(38):e2409160121.

[Singer, 1999] Singer, W. (1999). Neuronal synchrony: a versatile code for the definition of relations? Neuron, 24(1):49–65.

[Smith et al., 1997] Smith, S. W. et al. (1997). The scientist and engineer’s guide to digital signal processing.

[Srinath et al., 2021] Srinath, R., Emonds, A., Wang, Q., Lempel, A. A., Dunn-Weiss, E., Connor, C. E., and Nielsen, K. J. (2021). Early emergence of solid shape coding in natural and deep network vision. Current Biology, 31(1):51– 65.

[Thorpe et al., 2001] Thorpe, S., Delorme, A., and Van Rullen, R. (2001). Spike-based strategies for rapid processing. Neural networks, 14(6-7):715–725.

[Thorpe and Imbert, 1989] Thorpe, S. J. and Imbert, M. (1989). Biological constraints on connectionist modelling. Connectionism in perspective, pages 63–92.

[Tsao et al., 2006] Tsao, D. Y., Freiwald, W. A., Tootell, R. B., and Livingstone, M. S. (2006). A cortical region consisting entirely of face-selective cells. Science, 311(5761):670–674.

[Van den Bergh et al., 2010] Van den Bergh, G., Zhang, B., Arckens, L., and Chino, Y. M. (2010). Receptive-field properties of v1 and v2 neurons in mice and macaque monkeys. Journal of Comparative Neurology, 518(11):2051–2070.

[Vanni et al., 2020] Vanni, S., Hokkanen, H., Werner, F., and Angelucci, A. (2020). Anatomy and physiology of macaque visual cortical areas v1, v2, and v5/mt: bases for biologically realistic models. Cerebral Cortex, 30(6):3483–3517.

[VanRullen et al., 2005] VanRullen, R., Guyonneau, R., and Thorpe, S. J. (2005). spike_times make sense. Trends in neurosciences, 28(1):1–4.

[Wallis and Rolls, 1997] Wallis, G. and Rolls, E. T. (1997). Invariant face and object recognition in the visual system. Progress in neurobiology, 51(2):167–194.

[Yamins et al., 2014] Yamins, D. L., Hong, H., Cadieu, C. F., Solomon, E. A., Seibert, D., and DiCarlo, J. J. (2014). Performance-optimized hierarchical models predict neural responses in higher visual cortex. Proceedings of the national academy of sciences, 111(23):8619–8624.

[Zador, 2019] Zador, A. M. (2019). A critique of pure learning and what artificial neural networks can learn from animal brains. Nature communications, 10(1):1–7.

